# Population differences of chromosome 22q11.2 duplication structure predispose differentially to microdeletion and inversion

**DOI:** 10.1101/2025.07.04.662981

**Authors:** David Porubsky, DongAhn Yoo, Philip C. Dishuck, Nidhi Koundinya, Erika Souche, William T. Harvey, Katherine M. Munson, Kendra Hoekzema, Daniel D. Chan, Tiffany Y. Leung, Marta S. Santos, Senne Meynants, Ann Swillen, Jeroen Breckpot, Vasiliki Tsapalou, Patrick Hasenfeld, Jan O. Korbel, Peter M. Lansdorp, Joris R. Vermeesch, Evan E. Eichler

**Affiliations:** Department of Genome Sciences, University of Washington School of Medicine, Seattle, WA, USA; European Molecular Biology Laboratory (EMBL), Genome Biology Unit, Heidelberg, Germany; Laboratory of Cytogenetics and Genome Research, Centre for Human Genetics, KU Leuven, Leuven, 3000, Belgium; Terry Fox Laboratory, BC Cancer Research Institute, Vancouver, BC, Canada; Department of Medical Genetics, University of British Columbia, Vancouver, BC, Canada; Department of Human Genetics, Centre for Human Genetics, University Hospitals Leuven, Leuven, 3000, Belgium; Howard Hughes Medical Institute, University of Washington, Seattle, WA 98195, USA

## Abstract

The most common genomic disorder, chromosome 22q11.2 microdeletion syndrome (22q11.2DS), is mediated by highly identical and polymorphic segmental duplications (SDs) known as low copy repeats (LCRs; regions A-D) that have been challenging to sequence and characterize. Here, we report the sequence-resolved genomic architecture of 135 chromosome 22q11.2 haplotypes from diverse 1000 Genomes Project samples. We find that more than 90% of the copy number variation is polarized to the most proximal LCR region A (LCRA) where 50 distinct structural configurations are observed (∼189 kbp to ∼2.15 Mbp or 11-fold length variation). A higher-order SD cassette structure of 105 kbp in length, flanked by 25 kbp long inverted repeats, drives this variation and emerged in the human–chimpanzee ancestral lineage later expanding in humans ∼1.0 [0.8-1.2] million years ago. African LCRA haplotypes are significantly longer (p=0.0047) when compared to non-Africans yet are predicted to be more protected against recurrent microdeletions (p=0.00053) due to a preponderance of flanking SDs in an inverted orientation. Conversely, we identified nine distinct inversion polymorphisms, including five recurrent ∼2.28 Mbp inversions extending across the critical region (LCRA-D) and four smaller inversions (two LCRA-B, one LCRC-D, and one LCRB-D); 7/9 of these events were identified in haplotypes of African and admixed American ancestry. Finally, we sequence and assemble four families and show that LCRA-D deletion breakpoints map to the 105 kbp repeat unit while inversion breakpoints associate with the 25 kbp repeats adjacent to palindromic AT-rich regions. In one family, we observe evidence of more complex unequal crossover events associated with gene conversion and multiple breakpoints. Our findings suggest that specific haplotype configurations are protective and susceptible to chromosome 22q11.2DS while recurrent large-scale inversions help to explain why this syndrome is less prevalent among individuals of African descent.

## INTRODUCTION

The chromosome 22q11.2 region is a structurally complex region of the human genome characterized by the presence of large stretches of highly identical segmental duplications (SDs) also known as low copy repeats (LCRs). These are organized into eight SD blocks labelled LCRA-H (Shaikh et al. 2000) that predispose this region to recurrent rearrangements among various pairs of SDs (D. M. McDonald-McGinn et al. 2015). These homologous regions can misalign during meiosis via a mechanism known as non-allelic homologous recombination (NAHR) resulting in genomic rearrangements, including deletions, duplications, and inversions (Inoue and Lupski 2002). In ∼85% of cases, NAHR occurs between LCRA and D resulting in a deletion of a ∼3 Mbp long critical region (Edelmann et al. 1999). This deletion, also known as DiGeorge or velocardiofacial syndrome, is now collectively referred to as the chromosome 22q11.2 deletion syndrome (22q11.2DS) (Morrow et al. 1995). It is recognized as the most common genomic disorder in humans with a prevalence of 1 in ∼3,000 live births (Blagojevic et al. 2021). Patients may manifest with a wide range of phenotypes, including developmental delay, intellectual disability, congenital heart defects, and cleft-lip/palate. The deletion is also known to be a major genetic risk factor for schizophrenia (D. M. McDonald-McGinn et al. 2015; Marshall et al. 2017; McClellan et al. 2024).

Despite extensive research on the genetic and clinical aspects of 22q11.2DS, it is unknown why this syndrome occurs an order of magnitude more frequently than most other genomic disorders mediated by NAHR (Blagojevic et al. 2021). The structural complexity of the LCRA-D duplication blocks at the 22q11.2 locus and the resulting challenges in accurately and completely assembling this region have limited our ability to fully characterize the mechanisms driving these rearrangements. Previous studies using fluorescence *in situ* hybridization (FISH) and optical mapping (Shimojima et al. 2011; Vervoort and Vermeesch 2022) provided valuable insights into the structural diversity of this locus, but these methods lack the resolution necessary to precisely map deletion breakpoints or explore the fine-scale diversity of the underlying LCRs flanking this region. More recently, optical mapping methods were coupled with ultra-long Oxford Nanopore Technologies (ONT) sequencing approaches to locally assemble and characterize breakpoints in 22q11.2DS families (Vervoort et al. 2024; Zhou et al. 2024). While this methodology helped refine recombinant breakpoint regions, standard assembly methods failed to fully resolve the chromosome 22q11.2 region or the complexity of underlying rearrangements in either patients or population controls.

Recent advances in long-read sequencing (LRS) technologies and assembly algorithms have addressed many of the previous limitations. Combining ultra-long-read sequencing from ONT with high-fidelity (HiFi) PacBio, for example, was critical for the production of the first gapless, telomere-to-telomere (T2T) human genome reference T2T-CHM13 (Nurk et al. 2022) including resolution of all LCR regions (Vollger et al. 2022). Lessons learned from that effort have now been applied to develop new assembly algorithms (Rautiainen et al. 2023; Cheng et al. 2024), which have facilitated population-level sequencing of nearly complete T2T human genomes (Ebert et al. 2021; Liao et al. 2023; G. A. Logsdon et al. 2024). In this study, we present a detailed analysis of complete chromosome 22q11.2 from more than 130 diverse human haplotypes and six nonhuman primate (NHP) species (Yoo et al. 2025) including macaque (Zhang et al. 2025). We use these data to not only understand the full extent of human genetic diversity of this complex region at the base-pair level but to put this variation into an evolutionary context revealing human-specific and population-enriched features that drive recurrent rearrangement of 22q11.2DS. We also apply these T2T sequencing technologies to the study of chromosome 22q11.2DS patients and their families identifying distinct hotspots of recurrent inversion and microdeletion. These findings help us to pinpoint predisposed haplotypes and those protected from the disease suggesting that not all parental haplotypes nor populations have equal probability of rearrangement and disease.

## RESULTS

### Complete assembly of chromosome 22q11.2 haplotypes

In this study, we focused our analysis on a 5 Mbp region corresponding to human chromosome 22q11.2 characterized by large SD blocks denoted LCRA to LCRD with A and D being the longest and most identical to each other (**Supplementary Fig. 1**) (D. M. McDonald-McGinn et al. 2015). This region harbors ∼54 protein-coding genes (**Fig. 1a**) and was only recently fully assembled as part of the T2T-CHM13 reference (coordinates: 18-23 Mbp). The prior GRCh38 reference contained three large gaps in LCRA as well as two unassigned contigs that have been subsequently integrated into T2T-CHM13 (**Fig. 1b**). We initially evaluated the completeness of 220 phased genome assemblies generated as part of the HPRC (Liao et al. 2023) and HGSVC (G. A. Logsdon et al. 2024). Of these, 135 haplotypes were unrelated and assembled in a single continuous contig, passing our assembly quality validations (Vollger et al. 2019) (**Supplementary Fig. 2, Methods**). The set included representation of all human ancestries (AFR - African, AMR - American, EAS - East Asian, SAS - Southeast Asian and EUR - European) with comparable numbers of haplotypes of African (n=59, 43.7%) and non-African ancestry (n=76, 57.3%) (**Fig. 1c, Supplementary Fig. 2**). The total length of the region varied considerably with a relatively constant 3.66 Mbp of unique sequence and highly variable duplicated bases (988,335-2,687,425 bp) mapping to the LCR regions (**Fig. 1d, Methods**). We observed one outlier (HG00731-H1) with ∼3.91 Mbp of “unique” sequence. The subsequent analysis showed a ∼270 kbp tandem duplication of a unique sequence located distal to LCRD, likely as a result of NAHR. This duplication was previously observed at a frequency of 0.002% among both, cases with neurodevelopmental delay and unaffected individuals (Cooper et al. 2011; Coe et al. 2014) (**Supplementary Fig. 3**).

**Figure 1:**
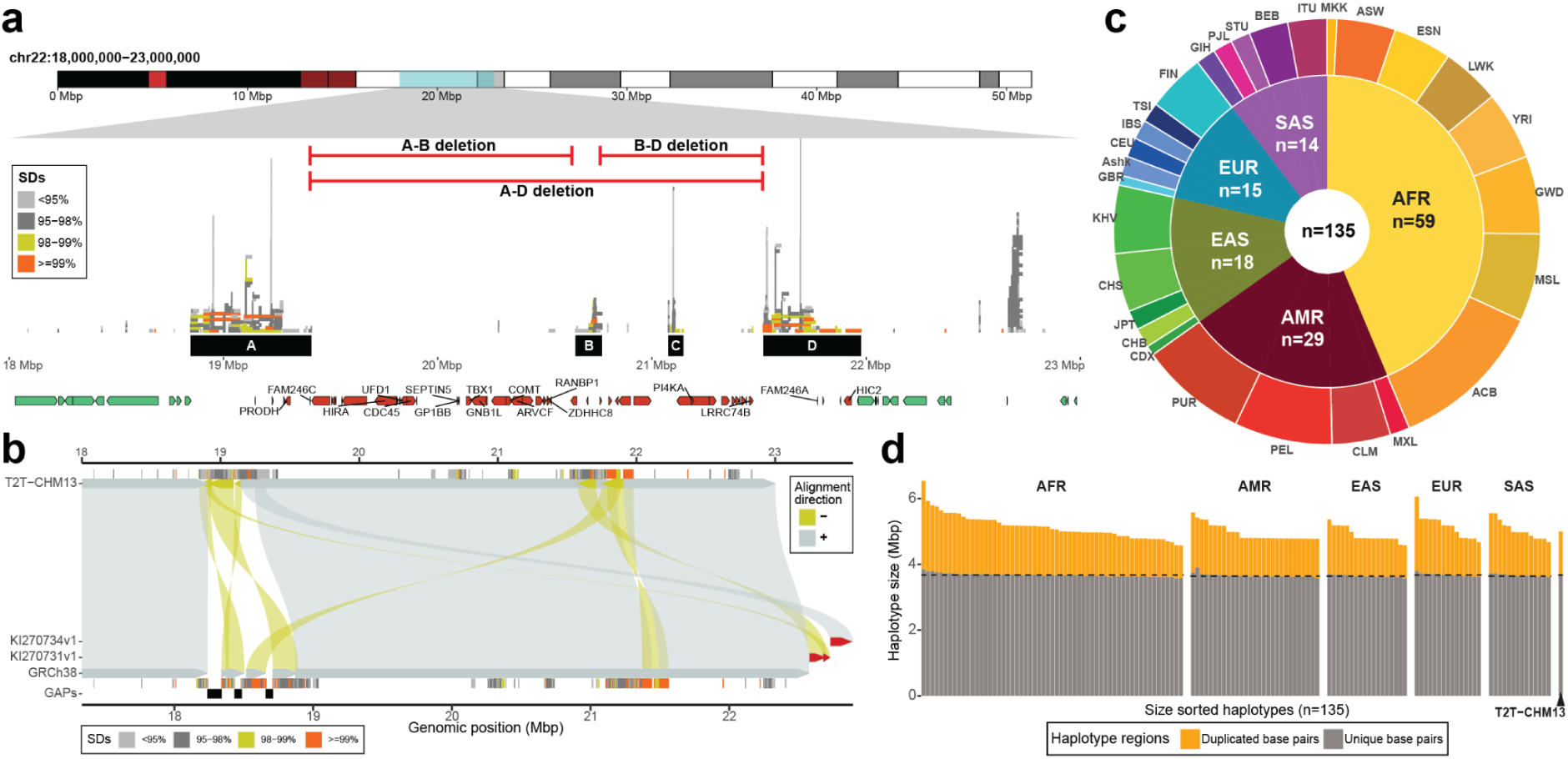
Assembly and annotation of human chromosome 22q11.2 region. **a)** Chromosome 22 ideogram (top) annotated with the critical region (chr22:18000000-23000000) (blue), pathogenic copy number variants (red horizontal lines), LCRA-D (black rectangles), segmental duplications (SDs; stacked horizontal bars colored by % identity in middle), and a subset of protein-coding genes (bottom, red and green arrowbars). There are a total of 76 protein-coding genes mapping between 18-23 Mbp of T2T-CHM13 reference of which 54 map between LCRA-D (in red). **b**) Comparison of the GRCh38 reference (query sequence, bottom) against the T2T-CHM13 reference (target sequence, top) over 22q11.2 with direct (‘+’, forward - gray) and inverted (‘-’, reverse - yellow) alignments distinguished by color and their respective orientation. Both query and target sequences depict SD annotation colored by sequence identity and gaps in GRCh38 (black rectangles). Unassigned contigs (GRCh38) are highlighted in red. **c**) A two-layer donut plot shows the counts of complete assemblies at 22q11.2 colored by major ancestries (AFR - African, AMR - American, EAS - East Asian, EUR - European, SAS - Southeast Asian). Specific 1000 Genomes Project (1KG) populations (outer layer) are indicated by an acronym (https://www.internationalgenome.org/data-portal/population). **d**) A barplot showing length variation of SD regions (orange) versus unique regions (gray); haplotypes are organized by five major ancestries. The median value of ‘unique’ base pairs is highlighted by a dashed line per ancestry. T2T-CHM13 proportions of ‘duplicated’ and ‘unique’ base pairs are shown separately.

### SD population variation and higher-order structure

We assigned SDs to LCR regions A-D for 133 haplotypes based on the canonical T2T-CHM13 structure (**Fig. 1**)—we excluded two haplotypes that carried a large inversion which complicated such assignment. Consistent with earlier optical mapping data (Demaerel et al. 2019), LCRA is, by far, the most variable (>90% of total size variability), followed by LCRD in contrast to LCRB and C, which are essentially invariant (**Fig. 2a, Supplementary Fig. 4**). The median size of LCRA is 577.15 kbp with haplotypes differing by more than 11-fold in length (189.01 to 2147.85 kbp) (**Fig. 2b**). We further decomposed LCRA into evolutionary ancestral units, termed duplicons, using DupMasker (Jiang et al, 2008). For LCRA, we identify 50 structurally distinct haplogroups of various size and structural complexity (**Fig. 2c, Methods**). There are six major LCRA haplogroups representing about half (n=74; 56%) of all haplotypes while 33 haplogroups occur as a single representative structure. Haplotypes of African ancestry are significantly longer than those of non-African ancestry (Wilcoxon test, unpaired, p=0.0047) (**Fig. 2c, inset i**).

**Figure 2:**
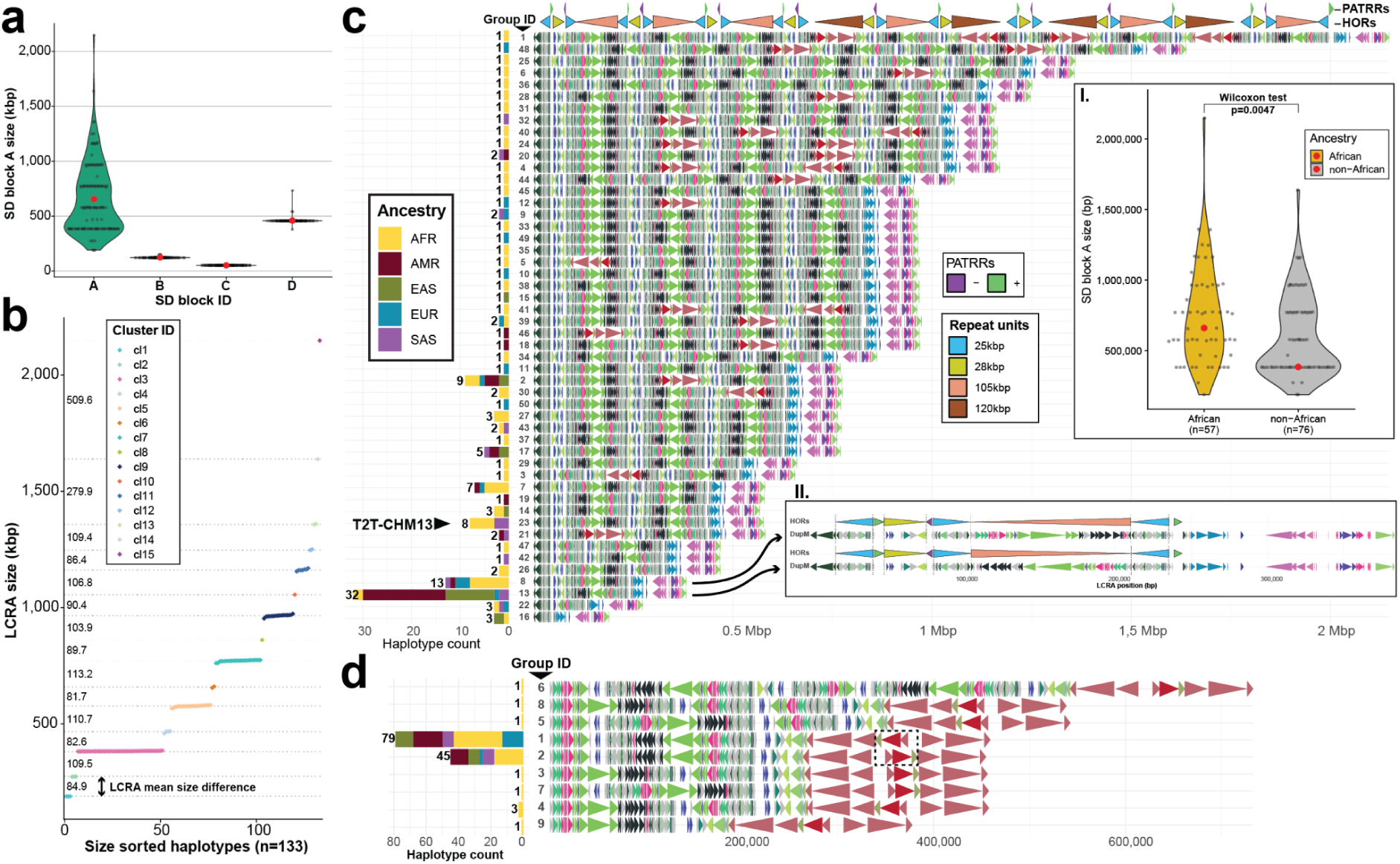
Chromosome 22q11.2 structural diversity and duplicon higher-order structure. **a**) Violin plot showing length distribution for LCRA-D along with mean SD block size (red dots). **b)** Size sorted distribution of LCRA size (colored by cluster ID) with the mean size of each cluster (horizontal dotted line) and the mean size difference between each step indicated. **c**) Duplicon architecture of 50 distinct LCRA haplogroups (colored arrowheads show size and orientation based on duplicons as defined by DupMasker). Stacked barplot shows the frequency of each haplogroup among continental population groups. On top there is an annotation of common repeat units (25, 28, 105 and 120 kbp in length) along with PATRRs. Inset **I**) Shows a significant size difference (Wilcoxon test, unpaired, p=0.0047) of LCRA among haplotypes of African and non-African ancestry; inset **II**) Zoom-in structure of the two most abundant haplogroups (8 and 13) with annotation of the 105 (red), 25 (blue) and 28 kbp long repeat units. **d**) Duplicon architecture of nine distinct LCRD haplogroups (DupMasker) and their population distribution (left stacked bar plot). A 48.65 kbp inversion distinguishing haplogroups 1 and 2 is highlighted by a dashed rectangle (**Supplementary Fig 5**).

The two most abundant haplotype groups (8 and 13) are comparatively short for LCRA (median: 383.17 kbp) but population stratified. Haplogroup 13, for example, is composed almost exclusively of haplotypes of non-African ancestry while haplogroup 8 is enriched among Africans. They differ only by a single inverted segment of the ∼105 kbp duplication unit (**Fig. 2c, inset ii**). In contrast to LCRA, distal LCRD shows far fewer structural haplogroups (n=9) with two major ones representing 93.23% (124 out of 133) of all haplotypes. These two haplogroups (1 and 2) differ only by a single inversion (∼48.65 kbp in size, flanked by ∼70 kbp SDs) at the distal portion of LCRD (**Supplementary Fig. 5**). In contrast to LCRA, the most common LCRD haplotypes are distributed similarly among African and non-African ancestries although all remaining unique haplotypes (n=9) are of African ancestry consistent with greater genetic diversity (**Fig. 2d**).

An analysis of cumulative LCRA length confirms a stepwise increase of a higher-order duplicon structure (**Fig. 2b**, **Methods**). Instead of the predicted ∼160 kbp higher-order structure (Guo et al. 2016; Vervoort et al. 2024), we refined the basic repeat unit to be of 155 kbp in length, consisting of a 105 kbp core flanked by 25 kbp long inverted repeats, (**Supplementary Fig. 6**), with base-pair resolution as opposed to the former prediction by optical mapping (Demaerel et al. 2019). In addition to these repeats, we also define two other common repeats between LCRA and D of 28 and 120 kbp in length, respectively, which adds up to a higher-order repeat (HOR) structure composed of four common repeats (**Supplementary Fig. 7, Methods**). This HOR structure can also be defined based on a set of repeating duplicon units (**Supplementary Fig. 8, Methods**) and was generally observed by fiber-FISH, albeit not at base-pair resolution (Demaerel et al. 2019). At the boundaries of 25 and 28 kbp long HORs, we locate the previously defined pockets of palindromic AT-rich regions (PATRRs) (Kurahashi et al. 2000; Vervoort et al. 2024). We define a common, non-AT-rich sequence motif overrepresented within PATRRs to help us determine the orientation of each AT-rich unit (**Supplementary Fig. 9a, Methods**). We find this motif present not only at the 22q11.2 region but also at other short acrocentric arms of chromosomes 13, 14 and 21, which are frequently associated with Robertsonian translocations (**Supplementary Fig. 9b**) (Page et al. 1996).

Parsimoniously, we explain most of the stepwise increase in LCRA size as a result of the cumulative addition of the basic HOR units (155 kbp + 28 kbp) flanked by PATRRs (∼10 kbp) via NAHR likely occurring between directly oriented 25 kbp repeats. This results in a unit increase of ∼193 kbp distinguishing the most abundant LCRA haplotype lengths (**Supplementary Fig. 10, Methods**). The net effect is that the PATRRs and HOR units create a set of embedded palindromes that predispose LCRA to recurrent insertions, deletions, and inversions among directly and indirectly oriented repeats (**Supplementary Figs. 11-12**). When assessing the sequence identity between all paralogous copies of four HOR repeats, we find most are 99% identical with one exception—the 25 kbp repeat shows significantly greater divergence (**Supplementary Fig. 13**). We tracked the reduced sequence identity of the 25 kbp unit to a GC-rich 27 bp variable number tandem repeat (VNTR) that is highly variable among paralogs and alleles and is transcribed as part of an exon of a long noncoding RNA (*FAM230*) (**Supplementary Figs. 14-15**). We provide a detailed summary of gene annotation within the 22q11.2DS region, including LCRA-D based on mapping of long-transcriptomic Iso-Seq data (**Supplementary Notes**).

### Evolutionary history of the chromosome 22q11.2 region

In order to reconstruct the evolutionary history of the chromosome 22q11.2 in humans, we compared human haplotypes to the recently completed NHP genomes (chimpanzee - PTR, bonobo - PPA, gorilla - GGO, and Bornean (PPY) and Sumatran (PAB) orangutans) (Yoo et al. 2025) (**Fig. 3a-b**). Overall, the majority (95.49% or 127/133) of human chromosome 22q11.2 haplotypes are expanded >2-fold for SD content (range: 313-2107.8 kbp; median: 687 kbp) when compared to the syntenic regions among the great apes (range: 103-464.9 kbp; median: 268.4 kbp) due to an increase in copy number of the HOR units (**Fig. 3b**). Comparing the chimpanzee haplotype to the most similar human (HG02018, EAS from Vietnam) haplotype, we observe that the proximal 25 kbp segment and large part (∼97 kbp) of the 105 kbp segment map to LCRC whereas the 105 and 25 kbp units typically reside in both LCRA and D in humans. The distal 25 kbp segment and the remaining 8 kbp of the 105 kbp segment map to LCRD in chimpanzee. This suggests a transposition from LCRC into D to constitute the full 155 kbp HOR structure as well as its association with PATRRs in humans (**Fig. 3c**). These data further support the spread and expansion of the 105 kbp long repeat unit from LCRD into LCRA where it appears in multiple copies (1-6 copies) while in LCRD it is conserved as a single unit (**Supplementary Fig. 16**). Based on the comparison to chimpanzee, we observe three ancestral human LCRA structures followed by three intermediate structures supporting stepwise addition of repeat units at LCRA (**Supplementary Fig. 17**).

**Figure 3:**
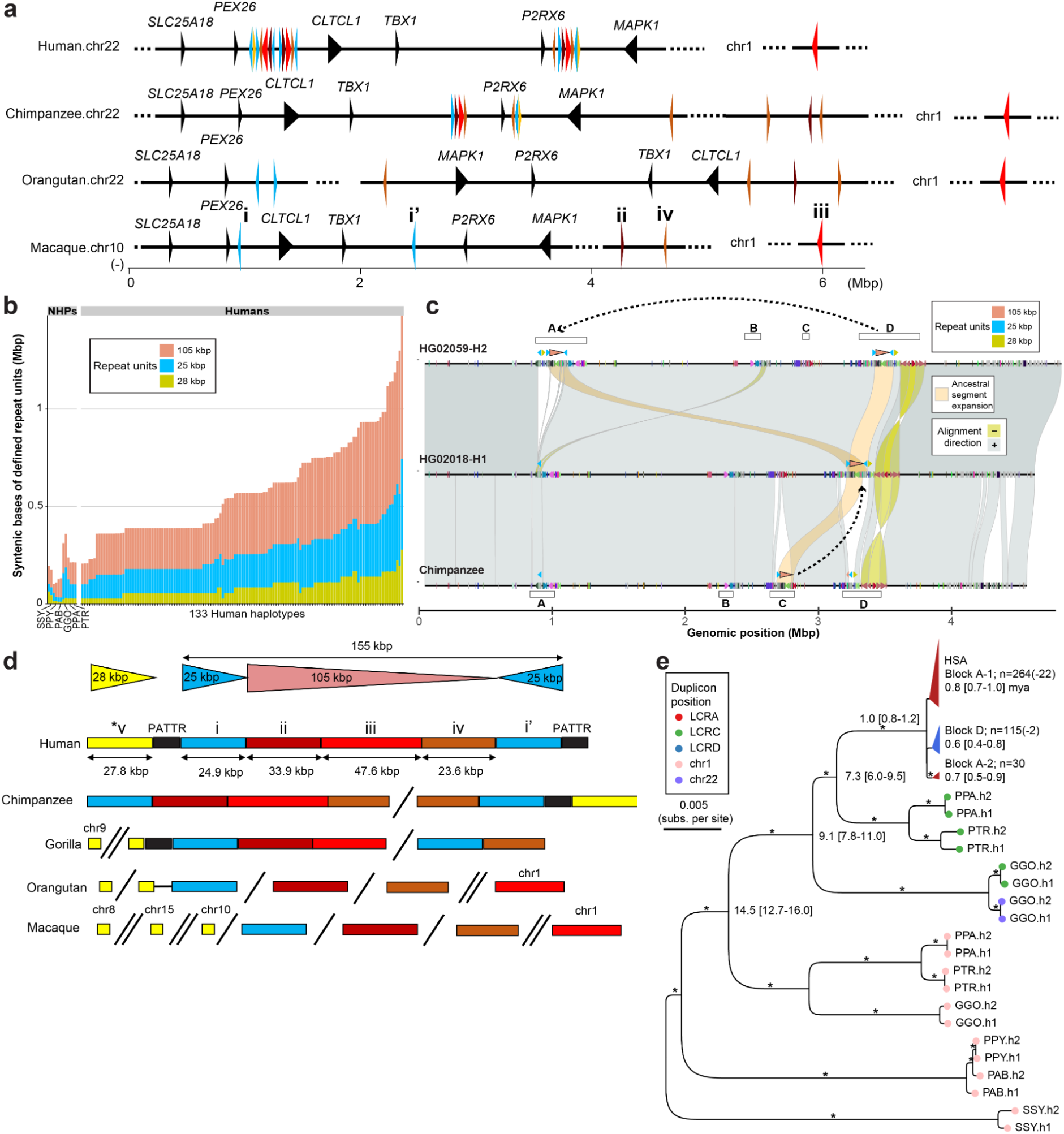
Evolutionary history of 22q11.2 and the emergence of higher-order SD structure. **a**) Origin of individual subunits coming from multiple segments of macaque and great apes (Bornean orangutan, chimpanzee and human). Syntenic genes are indicated. **b**) Stacked barplot showing the total size in base pairs for the 25 kbp (blue), 28 kbp (yellow) and 105 kbp (light red) repeat units in humans and nonhuman primates (NHPs). **c**) Miropeats-style plot showing alignments (direct - gray, ‘+’ and inverted - yellow, ‘-’) between chimpanzee (AG18354_PTR-H1, bottom) and two human samples HG02018-H1 (middle) and HG02059-H2 (top). There is a DupMasker annotation at the query and target line shown as directional arrowheads colored by unique duplicon ID. Alignments of 105 and 25 kbp long repeat units between chimpanzee and human haplotypes are highlighted by light orange color (aligned region ∼130 kbp). Positions of LCRA-D in both target and query are shown as empty rectangles. A dashed arrow shows transposition of the chimpanzee repeat unit from LCRC to D and then expansion from LCRD to A in humans. **d**) Inferred structure of the complex duplication unit composed of the five smaller duplicons. From top to bottom, the largest homologous units located in chr22 of human to macaque are shown, which differ in size. *v duplicon is represented by ∼4 kbp sequence in the gorilla, orangutan, and macaque genomes. **e**) Maximum likelihood phylogenetic tree of subunit iii (n=421 + 1 macaque outgroup), with conserved sequence length of 27.8 kbp out of 47.6 kbp total length. The human copies include 274 and 135 originated from LCRA and D, respectively, which are collapsed in the tree as two clades for A and one clade for D group. The number of sequences corresponding to each group is indicated with the parenthesis numbers representing non-A or -D block copies within each clade. The tree was outgrouped with the single copy ortholog of haploid macaque genome (not displayed), which was used to locate the root of the ape phylogeny. Ortholog copies from the apes include single or expanded copies from 12 haplotype assemblies of chimpanzee (PTR), bonobo (PPA), gorilla (GGO), Bornean orangutan (PPY), Sumatran orangutan (PAB), and siamang (SSY). Bootstrap support greater than 95 is indicated by *.

To investigate deeper phylogeny of the duplication, we tracked the synteny across primates for 25 kbp and 105 kbp, which forms 155 kbp basic unit (2*25 kbp and 105 kbp), 28 kbp, and 120 kbp units. We identified the underlying smaller duplicons independently (marked i-v in **Fig. 3d** and vi-viii in **Supplementary Fig. 18**) based on synteny among the apes using the macaque genome as an outgroup (Zhang et al. 2025). While duplicons i, ii, and iv originate as singleton or doubleton syntenically on chromosome 22 (**Fig. 3a**), others such as duplicon iii map to a single copy on chromosome 1 (human synteny) and, thus, is predicted to have duplicatively transposed interchromosomally from chromosome 1 to chromosome 22 in the common ancestor of human, chimpanzee, and gorilla. Unlike other duplicons, we find that duplicon v is relatively young and shared between human and *Pan* lineages but is missing from gorilla and more diverged genomes. Tracing the source sequence of duplicon v in macaque, we find more than half of the sequence was untraceable while the remaining half is composed of three ∼4 kbp units, all of which map to different chromosomes (chr8, chr10 and chr15). Comparing primate genomes, we observe a gradual increase in higher-order duplication complexity as we span the primate tree from macaque, gorilla, *Pan* and then to human with the data suggesting that the higher-order structure arose by a series of events (translocation, transposition, and juxtaposition of the smaller duplicon units as illustrated in **Fig. 3d, Supplementary Fig. 19, Methods**). Notably, we also find that more than 94% of duplication blocks present at chromosome 22q11.2 demarcate the boundaries of inversions between humans and NHPs suggesting the formation of these higher-order structures was frequently accompanied by inversion toggling (Porubsky et al. 2022) of the unique sequence bracketed by SDs (**Supplementary Fig. 19**).

Wherever possible, we constructed a phylogenetic tree for each of the smaller duplicons, including duplicon iii, to estimate the time when more complex SD structures formed (**Fig. 3e; Supplementary Figs. 20-30**). Calibrating based on macaque–human divergence time of 28.8 million years ago (MYA), we estimate, for example, that the duplicative transposition (duplicon iii) from chr1 to chr22 along with the juxtaposition of i-iii duplicons occurred 14.5 [12.7-16.0] MYA consistent with the comparative analysis (**Fig. 3e; Supplementary Fig. 19**). The segment further fused with duplicon iv ∼7.3 [6.0-9.5] MYA giving rise to the ∼130 kbp structure present in both human and *Pan* lineages. The complete higher-order structure arose specifically in the human lineage and we estimate the 25 kbp repeat unit began to duplicate 2.1 [1.7-2.5] MYA—two of three human haplotypes with LCRA structures most similar to chimpanzee (HG00621-H2 - EAS from China and HG02018-H1 - EAS from Vietnam discussed above) form a distinct outgroup when compared to all human copies (**Supplementary Figs. 17 and 29**). We estimate that LCRA and D diverged even more recently, 1.0 [0.8-1.2] MYA with the majority of subsequent expansions and conversions occurring during the emergence of *Homo sapiens* for LCRA (coalescent time of 0.7 [0.5-0.9] and 0.8 [0.7-1.0] MYA) and LCRD (coalescent time of 0.6 [0.4-0.8] MYA) (**Fig. 3e**).

### Population stratification of large-scale chromosome 22q11.2 inversions

The 22q11.2 region has been subjected to a large number of structural rearrangements, especially large inversions distinguishing humans and NHPs (**Supplementary Fig. 30**). We specifically focused on the identification of large-scale inversion polymorphisms between LCRs A-D in humans whose existence has been controversial (Demaerel et al. 2017). We also considered inverted duplications that change the orientation of HOR units within the LCRs but do not affect the flanking sequence. For this purpose, we utilized both the assemblies from Human Pangenome Reference Consortium (n=192) and pooled Strand-seq data (n=271) for validation and genotyping of inversions (Porubsky et al. 2022). In total, among 539 samples we identified 11 large inversions (mean size 1.9 Mbp) and 6 inverted duplications (mean size 131.5 kbp) (**Supplementary Table 1**, **Supplementary Fig. 31**). The former includes seven carriers of the largest (∼2.28 Mbp) A-D inversion extending across the chromosome 22q11.2DS critical region. Based on haplotype analysis, five of these are predicted to be, in fact, independent, recurrent mutations and, thus, each large inversion is extremely rare, occurring at a frequency <1% in the population (**Fig. 4a, Supplementary Fig. 32**). We also observe inversions of LCRA-B (1.7 Mbp, HG01695-EUR, HG01175-AMR), C-D (0.44 Mbp, HG03139-AFR), and LCRB-D (0.99 Mbp, HG03471-AFR) —all which were resolved at the sequence level (**Fig. 4a, Supplementary Fig. 33**). Notably, these large-scale inversions are restricted to African and admixed American populations and we observe a significant enrichment among African-Americans (Fisher’s exact test two-sided; p=0.02862, odds ratio: 5.262964) (**Supplementary Table 1**, **Supplementary Fig. 34**). With respect to inverted duplications, we identified six commonly inverted sites within LCRs of which four reside within LCRA and two within LCRD (**Supplementary Fig. 35a-c, Methods**). These smaller inverted duplications range in length from ∼12 kbp to up ∼208 kbp but are much more common (33.1% to 39.8% allele frequency among 133 haplotypes) when compared to the large inversions (**Supplementary Fig. 35d**). There was one exception, however, which was a proximal inversion mapping to LCRD present in only eight haplotypes (6% allele frequency–all of African ancestry).

**Figure 4:**
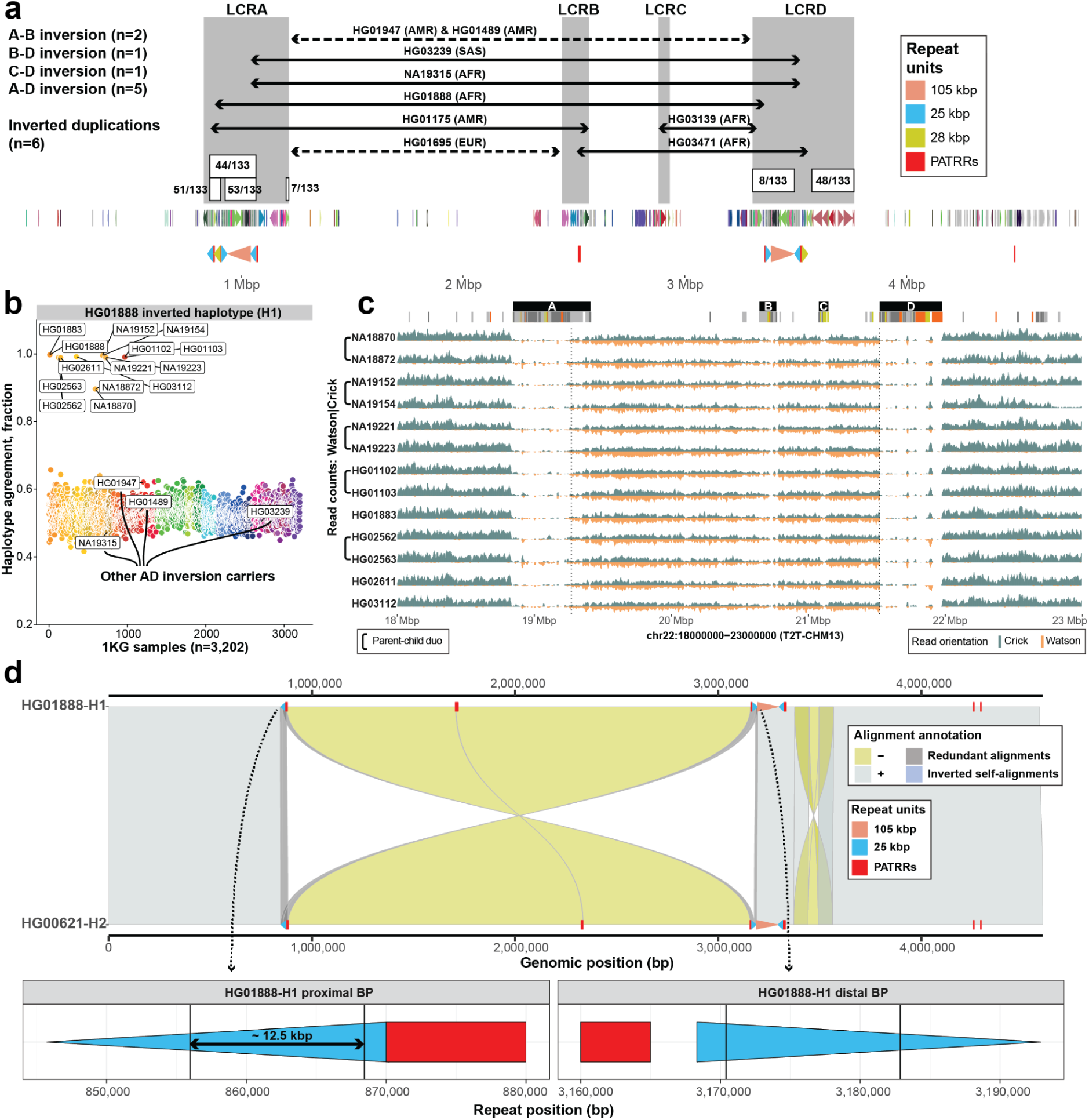
Chromosome 22q11.2 human inversion polymorphisms and breakpoints. **a**) Overview of all inverted ranges and their frequency with respect to a simple 22q11 haplotype (HG01890-H1). The gray rectangles highlight positions of LCRs A-D. The solid black arrows mark assembly-resolved inversions while dashed black arrows marked Strand-seq supported inversions. Each range is marked by a sample name where a given inversion was detected. Inverted duplications found within LCRA and D are marked by white rectangles with a number denoting predicted inversion frequency among 133 haplotypes. Below there is a duplicon annotation shown as directional arrowheads colored by a unique duplicon ID. Last, positions of common repeat units (25, 28, and 105 kbp) are shown as colored arrowheads. Positions of PATRRs are shown as red rectangles. **b**) Predicted carriers of inverted haplotype HG01888-H1 among 1KG sample panel (n=3,202) (Lalli et al. 2025). Each dot represents a single sample colored by a population ID for the five major ancestries (AFR - African, AMR - American, EAS - East Asian, SAS - Southeast Asian, and EUR - European). Samples labelled with their IDs (n=13 + HG01888) are those predicted as possible carriers of an inverted haplotype (HG01888-H1) as a measure of haplotype concordance. **c**) Read-coverage profiles of Strand-seq data over the 22q11.2 region summarized as binned (bin size: 10 kbp step size: 1 kbp) read counts represented as bars above (teal; Crick read counts) and below (orange; Watson read counts) the midline. Dotted lines highlight the inversion region between LCRA and LCRD. Here, equal coverage of Watson and Crick counts represents a heterozygous inversion as only one homologue is inverted with respect to the reference (T2T-CHM13). Above there is an SD annotation and marked boundaries of SD blocks (A to D) as black rectangles. **d**) Miropeats-style plot showing the alignments (direct ‘+’ - gray, inverted ‘-’ - yellow) between direct (HG00621-H2) and inverted (HG01888-H1) haplotypes between LCRA and LCRD. Positions of 105 (red) and 25 (blue) kbp long repeat units are shown on top of the query and target sequence along with PATRRs shown as red rectangles. Redundant (overlapping) alignments at the inversion boundaries are highlighted in dark gray color. Below there is a visualization of predicted inversion breakpoints (∼12.5 kbp region) within inversely oriented 25 kbp long repeat units in proximal (LCRA) and distal (LCRD) regions.

We selected one of the large A-D inversions (HG01888-H1) originally discovered using Strand-seq data (Porubsky et al. 2023) and subsequently assembled in this study (**Fig. 4a**) for population genotyping against the full 3,202 from 1000 Genomes Project (1KG) diversity panel (Byrska-Bishop et al. 2022; Lalli et al. 2025). Using simple genotyping concordance for single-nucleotide polymorphisms (SNPs) identified on this haplotype, we predicted 13 samples with this inverted haplotype (**Fig. 4b, Methods**). We tested for the presence of the inversion using Strand-seq confirming all 13 samples as *bona fide* heterozygous carriers of this large inversion (**Fig. 4c**). Among the 13, five correspond to parent–child duos confirming germline transmission of the event (as opposed to potential somatic rearrangements). Thus, we identified eight unrelated carriers for an estimated worldwide allele frequency of 0.16% (8 out of 5,008 haplotypes from 2,504 unrelated individuals) of this particular inverted haplotype. There was, however, almost complete stratification of this inversion between African and non-African samples with 7/8 being identified from 1KG samples collected from the African continent (**Fig 4b).**

The inverted haplotype in a single non-African sample was of admixed ancestry (PUR-HG01102/3) and we determined that this segment is ancestrally of African origin (**Supplementary Fig. 34**). We repeated the genotyping for the remaining four LCRA-D inversions, but each was predicted to occur as a single occurrence as opposed to multiple carriers of HG01888 inversion (**Supplementary Fig. 36**). Similarly, we predict other large inversions (A-B, B-D and C-D) as a single occurrence, except for the A-B inversion in HG01175 predicted to have one additional carrier (HG01074). (**Supplementary Fig. 37**).

In order to map the breakpoints of the inversions and inverted duplications, we characterized the selected completed assemblies (**Methods**). In cases, such as HG01888-H1 and NA19315-H2, it was necessary to generate additional LRS data to fully assemble the region (**Fig. 4a**). Detailed analysis of HG01888-H1, for example, refines the breakpoint to a ∼12.5 kbp segment mapping within inversely oriented 25 kbp repeat units adjacent to PATRRs (**Fig. 4d, Supplementary Fig. 38, Methods**). Similarly, we mapped A-to-D inversion breakpoints in samples NA19315 and HG03239. Despite these inverted haplotypes being more complex than in HG01888, we narrowed down the inversion breakpoint to the vicinity of the 25 kbp repeat unit adjacent to PATRRs (**Supplementary Fig. 39**). Characterizing the inversion that occurred between LCRB and D in sample HG03471 predicts that the breakpoint occurs specifically between PATRRs as there is no copy of 25 kbp repeat unit in LCRB (**Supplementary Fig. 33**). In case of inverted duplications, we found all but one of the LCRA inverted duplications (n=3) being flanked by 25 kbp long repeats. In contrast, proximal and distal inversions in LCRD are mediated by longer 40.7 and 71 kbp long inverted repeats, respectively (**Supplementary Fig. 35**).

### Predisposition to 22q11.2DS predicted to differ by ancestry

Because the majority of chromosome 22q11.2 recurrent microdeletions arise as a consequence of NAHR between LCRA and LCRD driven in part by the 105 kbp higher-order structure (Vervoort et al. 2024), we examined the orientation, the length, and percent identity of the SDs for the 133 sequence-resolved haplotypes. Focusing initially on the 105 kbp core duplicon structure, we find that about a third of all haplotypes (n=43) carry exactly one copy of the 105 kbp repeat unit in both LCRA and D, differing only by an inversion of the 105 kbp repeat unit in LCRA. This renders the 105 kbp repeat units in either a direct (n=31) or an indirect (n=12) orientation between LCRA and D (**Extended Data Fig. 1a, Supplementary Fig. 40**) making them, theoretically, either predisposed or protected for A-to-D deletion, respectively. We extended this analysis to all 133 human haplotypes and tracked the orientation of all self-aligning sequences between LCRA and D (**Methods**). Focusing on self-alignments of 20 kbp and longer, we observe that with increasing alignment length the identity also increases. As expected, the vast majority (86.6%, 291 out of 336) of large (≥120 kbp) alignments involve 105 kbp core structure (**Supplementary Fig. 41**).

Considering only the longest self-alignments (≥120 kbp), we classified and rank ordered 22q11.2 haplotypes as either predisposed to LCRA-to-D inversion or A-to-D deletion (**Extended Data Fig. 1b and c**) under the assumption that longer, more identical directly orientated duplications would predispose to the latter while inverted duplications would promote the former (Cooper et al. 2011; Inoue and Lupski 2002). We find that approximately half of all human haplotypes (n=71 out of 134) have only inverted (23.1%, 31/134) or direct (29.9%, 40/134) alignments between LCRA and D, thus classified as fully protected or predisposed to A-to-D deletion, respectively (**Extended Data Fig. 1c**). Of note, we find significantly more haplotypes of African ancestry are predicted as being fully protected against 22q11.2DS than those of non-African ancestry (Fisher’s exact test, two-sided, p=0.00053, odds ratio: 0.1627054) (**Extended Data Fig. 1d, Supplementary Fig. 42**) consistent with the >3-fold reduced prevalence reported for African Americans epidemiologically (D. M. McDonald-McGinn et al. 2005).

### Sequence breakpoint characterization of 22q11.2DS patients

To directly test the relationship between duplication haplotype structures and sporadic chromosome 22q11.2 deletions associated with disease, we sequenced and assembled the 22q11.2 region from four families where a *de novo* deletion had occurred in a child with developmental disabilities. Because the transmitting parent was already known in three cases (Vervoort et al. 2024), we sequenced and assembled parent–child duos while for the fourth we considered all three members of the parents–child trio (**Data availability**). Using LRS and assembly approaches, we fully assembled and performed QC analysis on the chromosome 22q11.2 region for each transmitting parent and a child (proband) (**Methods**). We compared the phased assembly of the proband (with A-to-D deletion) with both haplotypes (H1 and H2) of the transmitting parent to define at the sequence level the breakpoint of the rearrangement. Given the accuracy of the assembly, near-perfect sequence identity could be used to readily track parental haplotypes against the child’s assembled chromosome (**Fig. 5a**, as we have shown previously utilized (Porubsky, Dashnow, et al. 2025)), in order to define recombination breakpoints.

**Figure 5:**
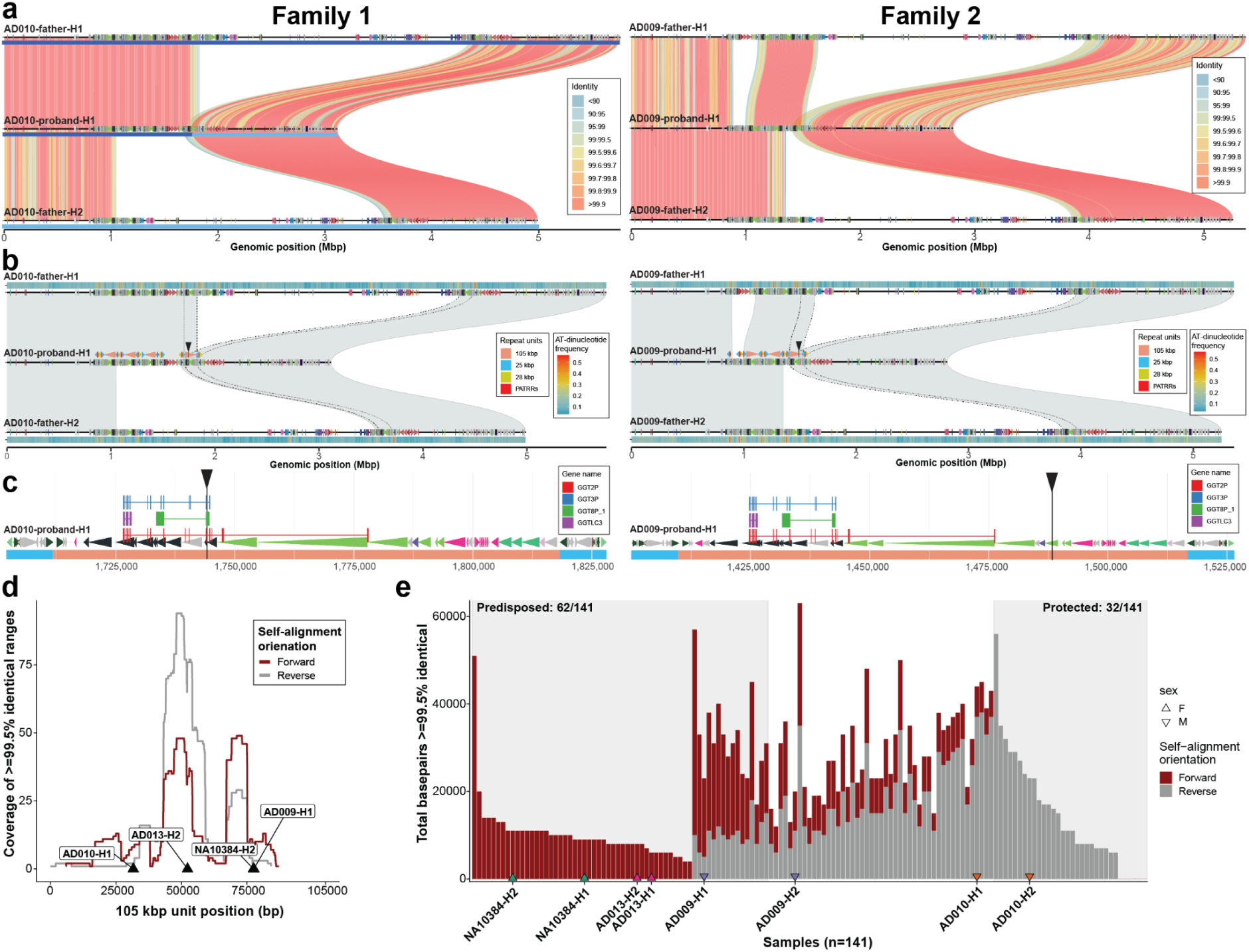
Parent–child chromosome 22q11.2 microdeletion rearrangements. **a**) Visualization of binned (bin size: 10 kbp) sequence identity between probands of family 1 and family 2 with respect to both haplotypes (H1 and H2) of a transmitting parent. The proband haplotype is in the middle while parental haplotypes H1 and H2 are at the top and bottom, respectively. Alignments are colored by sequence identity. On top of each haplotype is a DupMasker annotation depicted as arrowheads colored by unique duplicons. For family one we also highlight inherited paternal segments in the proband (H1 - dark blue, H2 - light blue). **b**) Similar to panel a, alignments shown between proband and transmitting parent are colored by alignment orientation. Redundant alignments at the deletion breakpoint are highlighted by black dotted lines. As in a) there is DupMasker annotation on top of each haplotype. In addition, for probands we also show the annotation of common repeat units. The mapped breakpoint within the 105 kbp long repeat is shown by black arrowhead. On top of parental haplotypes we also show the binned density (bin size: 5 kbp) of AT-dinucleotides for a given haplotype. **c**) Zoomed-in duplication structure at the narrowed down deletion breakpoint. Duplicons are shown as colored arrowheads. On top there are mapped exons of the *GGT* gene family. The predicted breakpoint is shown as a vertical black line. **d**) Coverage summary of the longest continuous region of high identity (≥99.5%) among all alignments between paralogous copies of 105 kbp long repeat units. We distinguish the coverage for the direct (forward, dark red) and inverted (reverse, gray) alignments. We mark the position of putative deletion breakpoints in four probands as black arrowheads. **e**) Stacked barplot of the total number of high-identity bases (≥99.5% identical) among self-alignments of 105 kbp repeat unit in forward (dark red) and reverse (gray) orientation. Shaded boxes highlight likely predisposed haplotypes as those where the majority (>0.5) of alignments are in forward orientation. Protected haplotypes are marked as those that have no alignments in forward orientation. At the base of relevant bars we mark haplotypes of transmitting parents (n=8), stratified by sex (point shape).

For family 1, a single breakpoint region was readily identified deleting ∼2.5 Mbp from H1, as a result of an unequal crossover between H1 and H2 likely when the quadrivalent was formed during meiosis I allowing for interchromosomal exchange between homologs. In this family, the deletion breakpoint occurs within the 105 kbp repeat unit (**Fig. 5b**) and we refine the breakpoint to a very narrow 36 bp range (**Fig. 5c, Methods**) mapping nearby an exon of the transcribed *GGT2* gene family. The pattern for family 2 was unexpectedly more complex, potentially involving both haplotypes although sequence flanking LCRA and LCRD would have seemingly predicted no exchange of flanking markers consistent with an intrachromosomal rearrangement involving a single homolog. Phased genome analysis, instead, suggests two breakpoints potentially as a result of a double recombination or gene conversion transposing ∼300 kbp from H2 to H1 of LCRA with a transition breakpoint in close proximity to 105 kbp repeat unit. The second breakpoint associated with the 2 Mbp deletion once again maps to the 105 kbp repeat unit (**Fig. 5b**), but in this case it is not associated with the *GGT2* gene family. We validated the reported haplotype switches using proband HiFi reads mapped back to the assembly and observed that mapped reads at each breakpoint are supportive of a given assembly (**Supplementary Figs. 43-44**). In family 3, we observe a nearly identical pattern of haplotype exchange between parental homologs as in family 1, consistent with unequal crossover within the 105 kbp repeat. Here, we narrowed down the deletion breakpoint to a 845 bp region (**Supplementary Fig. 45**). With respect to family 4 (the publicly available parent–child trio; **Data Availability**), we successfully assembled both parental haplotypes, but the proband (NA10382) haplotype carrying the LCRA-to-D deletion remained fragmented, preventing precise breakpoint definition as described above. We were able, however, to successfully pinpoint the putative deletion breakpoint based on split-read mapping using the transmitting parent’s chromosomes as a reference (NA10384) (**Supplementary Fig. 46a-b**). Taken together, all four families have deletion breakpoints mapping to the 105 kbp repeat unit albeit at different locations (**Fig. 5d**) with one family show a more complex pattern involving double recombination or gene conversion.

Finally, we projected all haplotypes of AD-deletion transmitting parents (n=4) against our initial population genetic analysis of fully assembled haplotypes (n=133) where we had previously predicted predisposition to or protection against the chromosome 22q11.2 deletion (**Methods**) based on the length, orientation, and percent identity (≥99.5%) of alignments between LCRA and D. Although we find the highest identity sequence between inversely oriented alignments cluster in the middle of the 105 kbp repeat unit, we find the highest identity regions between directly oriented repeats in two locations (around 50 kbp and 74 kbp). We found all deemed deletion breakpoints mapped within or in the vicinity of these high-identity regions within the 105 kbp repeat (**Fig. 5d**). When rank ordering parental haplotypes on our scale of AD-deletion predisposition, we find three out of four parents carry at least one haplotype predicted to be predisposed to AD-deletion suggesting that the sequence structure offers predictive power but is not absolutely deterministic of rearrangement susceptibility (**Fig. 5e**).

## DISCUSSION

Using T2T genomes, we provide a comprehensive view of the evolution and population diversity of one of the most structurally dynamic, disease-associated regions of the human genome with the aim of providing new mechanistic insights into its origin and mutational liability. Complete sequencing of human chromosome 22q11.2 revealed two remarkable features of human genetic diversity, namely, the rapid expansion of the higher-order structure of LCR copy number polymorphism and the low-level frequency of large-scale inversions. The discovery of large, rare, and recurrent inversions adds chromosome 22q11.2 microdeletion syndrome to the growing list of genomic disorders where inversion polymorphisms associate with hotspots of recurrent deletion and duplication (Porubsky et al. 2022). The frequent evolutionary and ongoing inversions are tightly coupled to the restructuring of the SDs although parental haplotypes do not carry nor directly predispose to rearrangement in the case of 22q11.2 microdeletions (Gebhardt et al. 2003). Previous studies based on fiber-FISH, optical mapping, and limited LRS (Demaerel et al. 2017; Pastor et al. 2020; Vervoort et al. 2024) reported few inversions yet never to this extent and most of the earlier reports of large inversions have been challenging to confirm (Demaerel et al. 2018). Here, we sequence resolve six distinct large inversions (ranging in size from 0.44 Mbp to ∼2.28 Mbp). We further validate a subset by orthogonal Strand-seq analyses of the corresponding cell lines and confirm germline transmission for eight events by examining parent-child trios (**Supplementary Table 1**). We find that inversion breakpoints cluster near PATRRs associated with the 25 kbp repeat unit (Kehrer-Sawatzki et al. 1997; Babcock et al. 2007).

The larger inversion polymorphisms are each exceedingly rare (<1%) but occur recurrently. For instance, we find the A-D inversion occurs independently in five different individuals (two African, two American, and one Southeast Asian ancestry). Remarkably, almost all large-scale inversions occur among individuals of African and admixed American ancestry. Population screening using genotype concordance of SNPs underlying the inverted haplotypes shows that one of the most common inversions, LCRA-D in sample HG01888 (haplotype 1), occurs at only 0.16% (8/5,008) allele frequency and is exclusive to African and admixed American individuals from the 1KG. In most other cases the inversions are exclusive to individual families suggesting that such inversions (unlike chromosome 8p23 and 17q21.31 inversions) (Koolen et al. 2006; Antonacci et al. 2009; Porubsky et al. 2023) fail to reach high frequency in the population possibly due to the action of purifying selection. These recurrent inversions affect dozens of unique genes mapping to the critical region and potentially the patterns of linkage-disequilibrium and gene expression. Although we focused on LCRA-D duplications, it is likely that this inversion toggling persists for other SD blocks along chromosome 22. For example, we document a large novel inversion in a single sample of African ancestry (NA20129) that extends from LCRD to LCRH positioned ∼3.5 Mbp away (**Supplementary Fig. 47**).

The other remarkable feature was the polarized variation of LCRA compared to the other duplication blocks (LCRC-D). We show that “normal” human LCRA haplotypes can differ by 11-fold with respect to SD content—placing it among the most copy number variable euchromatic regions in the human genome (Sudmant et al. 2015; Ebert et al. 2021). Previous studies using fiber-FISH and optical mapping predicted extensive tandem variation in LCR copy number (Shimojima et al. 2011; Guo et al. 2016; Pastor et al. 2020; Demaerel et al. 2019; Vervoort and Vermeesch 2022). Sequence-based resolution and duplicon characterization, however, reveal a much more complex and nuanced pattern of alternating tandem and palindromic HOR structures. Consistent with recent genome-wide surveys (Jeong et al. 2025), we observe some of the highest copy number (and therefore largest LCRA regions) among individuals of African descent. In our sample of 50 distinct LCRA structures, we find that 11 out of 15 of the longest LCRA blocks occur among individuals of African origin. Of note, these expansions do not always correlate with a predicted increased probability for NAHR. Using simple assumptions regarding length, percent identity, and repeat orientation, our analysis predicts that individuals of African ancestry are more likely to be predisposed to inversions as opposed to large-scale *de novo* deletions (Fisher’s exact test; p=0.00053; odds ratio: 0.1627054, Africans are ∼3 times less predisposed to 22q11.2DS) (**Extended Data Fig. 1d, Supplementary Fig. 42**). Thus, despite the larger size of LCRA and higher-copy number of SDs, Africans are more likely to be fully protected against 22q11.2DS than those of non-African ancestry because of the preponderance of inversely oriented repeats between the LCR blocks. This sequence structure prediction matches closely an epidemiological observation made 20 years ago where 22q11.2DS deletions were found to be three times less common among patients of African-American ancestry (D. M. McDonald-McGinn et al. 2005). This reduced prevalence was subsequently shown not to be the result of phenotypic ascertainment biases and was hypothesized to be the result of differences in the LCR structures (D. McDonald-McGinn et al. 2023).

The human-specific expansion of LCRA centers around an ∼160 kbp SD block first described as a *GGT-*containing core duplicon (Jiang et al. 2007) and then later as a module harboring long noncoding lncRNA, *FAM230A/B* genes (Delihas 2018; Pastor et al. 2020). Based on our sequence analysis, we redefine the boundaries of this repeat block into 105 kbp core flanked by two 25-kbp inverted repeats that itself largely overlap with FAM230 lncRNAs (**Supplementary Fig. 14**). Our complete sequencing of human genomes (Ebert et al. 2021; Nurk et al. 2022; Liao et al. 2023) as well as NHPs (Yoo et al. 2025) confirms that this core duplicon expanded exclusively in the human lineage (Vervoort et al. 2021). We estimate that this expansion occurred very recently during the evolution of the *Homo* genus 750 thousand years ago (kya; CI 1,060 kya to 600 kya) spreading copies predominantly to LCRA and to a far lesser extent to LCRD). In orangutans and gorillas, this 105 kbp HOR structure does not exist; it is fragmented into at least three sections and distributed to other chromosomes with the fully constituted segment emerging only in Homo-*Pan* ancestral lineage. Among chimpanzees and bonobos, we find a single copy of the 105 kbp unit mapping to LCRC suggesting that this likely represents the ancestral location that was subsequently transposed and expanded in the human lineage to LCRA and D. Given the central role of this sequence in mediating breakpoints, it may help to explain the observation that 85% of microdeletions occur between LCRA and D where the density is highest (Blagojevic et al. 2021). Given the central role this 105 kbp duplicon plays in mediating NAHR and the 22q11.2DS, it follows from the comparative analysis that the predisposition to this deletion syndrome likely has been recently acquired during human evolution.

The accurate sequence assembly afforded by LRS allowed us to characterize in detail the breakpoints of both naturally occurring inversions in the general population as well as rearrangements associated with the most common form of the 22q11.2DS. We find that inversion and microdeletion breakpoints cluster to distinct regions in the LCR blocks—with inversion and inverted duplication breakpoints mapping to the 25 kbp repeat unit adjacent to PATRR sequences (Kehrer-Sawatzki et al. 1997; Kurahashi et al. 2000) while LCRA-D microdeletion patients map more broadly to the 105 kbp repeat unit. A recent study of 15 families using optical mapping and fiber-FISH confirmed that 13/15 families with LCRA-D deletions map to the ∼160 kbp duplicon (termed RL-AD1), which represents one of the largest and most identical repeats between LCRA and D (Vervoort et al. 2024). Inversion breakpoints, in contrast, map to a much smaller 25 kbp repeat unit adjacent to PATRRs first described as being associated with most common non-Robertsonian constitutional translocation in the human genome t(11:22) (Kurahashi et al. 2010). These two features of instability are related as the PATRRs appear to demarcate the edges of the core duplicon potentially helping to explain its hyperexpansion in the human lineage. LRS of recombinant haplotypes between the parent and child with 22q11.2DS, however, suggests a more complex mutational process than simple NAHR. In one of four LCRA-D cases, multiple exchange events involving both parental homologs are required to resolve the long-range haplotypes. These data suggest either double recombination events or long-range gene conversions mediated by human-specific core duplicons and the need to sequence characterize more families and patients with this rearrangement.

Finally, the complete sequence of 22q11.2 region along with characterization of the associated SNP haplotypes and SD structures should provide a powerful resource to further advance studies of human genetic disease. Several genome-wide association studies, for example, have focused on the identification of the genetic modifiers expressing schizophrenia in 22q11.2DS (Bassett et al. 2003; Gur et al. 2017; Cleynen et al. 2021). Because of the long-standing gaps in the reference genomes, the complexity of the SDs, the recurrent and independent inversion and rearrangements in the region, and hence the lack of linkage disequilibrium (LD) with flanking SNPs, those studies excluded the SDs—which are the source of most of the genetic variation in this region. Moreover, the existence of such large-scale inversions and their effect on expression, LD or phenotypes represents a new area of investigation. The sequence-resolved structural map of the 22q11.2 region we generated should provide a framework to include the SDs and inversions in future genome-wide association studies allowing investigators to explore the contribution of this variation to 22q11.2DS phenotypic variability and neuropsychiatric phenotypes.

## METHODS

### Generation of PacBio HiFi datasets

PacBio HiFi data were generated per manufacturer’s recommendations. Briefly, high-molecular-weight (HMW) DNA was extracted from frozen blood or cultured lymphoblasts using the Monarch® HMW DNA Extraction Kit for Cells & Blood (New England Biolabs, T3050L). At all steps, quantification was performed with Qubit dsDNA HS (Thermo Fisher, Q32854) measured on DS-11 FX (Denovix) and size distribution checked using FEMTO Pulse (Agilent, M5330AA & FP-1002-0275.) HMW DNA was sheared with Megaruptor 3 (Diagenode, B06010003 & E07010003) using settings 28/31 or 28/30 (depending on original length distribution) and used to generate PacBio HiFi libraries via the SMRTbell Prep Kit 3.0 (PacBio, 102-182-700). Size selection was performed with Pippin HT using a high-pass cutoff of 12 kbp (NA19315), 15 kbp (GM10382), or 17 kbp (all others) (Sage Science, HTP0001 & HPE7510.) Samples HG01888, GM10382, GM10383, and GM10384 were sequenced on the Sequel II platform on SMRT Cells 8M (PacBio, 101-389-001) using Sequel II Sequencing Chemistry 3.2 (PacBio,102-333-300) with 2-hour pre-extension and 30-hour movies, aiming for a minimum estimated coverage of 40× in PacBio HiFi reads, assuming a genome size of 3.1 Gbp. The remaining samples were sequenced on the Revio platform on SMRT Cells 25M. Samples beginning with “JV” and NA19315 were sequenced with Revio Chemistry V1 (PacBio, 102-817-900) with diffusion loading and 24-hour movies (JV) or Adaptive Loading and 30-hour movies (NA). The remaining duos were sequenced with Revio SPRQ chemistry (PacBio, 103-520-200) with Adaptive loading and 30-hour movies. All Revio samples aimed for a minimum estimated coverage of 30× in PacBio HiFi reads, assuming a genome size of 3.1 Gbp.

### Generation of ultra-long ONT datasets

Ultra-long ONT (UL-ONT) data were generated from DNA extracted from the lymphoblastoid cell lines (LCLs) using a modified phenol chloroform extraction protocol (G. Logsdon and Chuang 2022). Briefly, 3-5 × 10^7^ cells were lysed in a buffer containing 10 mM Tris-Cl (pH 8.0), 0.1 M EDTA (pH 8.0), 0.5% w/v SDS, and 20mg/mL RNase A (Qiagen, 19101) for 1 hour at 37°C. Next, 200 ug/mL Proteinase K (Qiagen, 19131) was added, and the solution was incubated at 50°C for 2 hours. DNA was purified via two rounds of 25:24:1 phenol-chloroform-isoamyl alcohol extraction followed by ethanol precipitation. Precipitated DNA was solubilized in 10 mM Tris (pH 8.0) containing 0.02% Triton X-100 at 4°C for two days. Libraries were constructed using the Ultra-Long DNA Sequencing Kit (ONT, SQK-ULK001 and SQK-ULK114) with modifications to the manufacturer’s protocol. Specifically, ∼40 ug of DNA was mixed with FRA enzyme and FDB buffer as described in the protocol and incubated for 5 minutes at RT, followed by heat-inactivation at 75°C. RAP enzyme was mixed with the DNA solution and incubated at RT for 1 hour before the clean-up step. Clean-up was performed using the Nanobind UL Library Prep Kit (Circulomics, NB-900-601-01) and eluted in 225 uL EB. Lastly, 75 uL of the library was loaded onto a primed R9.4.1 or R.10.4.1 flow cell for sequencing on the PromethION, with two nuclease washes and reloads after 24 and 48 hours of sequencing.

### Generation of Strand-seq datasets

Low-coverage Strand-seq data were generated for 13 inversion candidate samples (GM18870, GM18872, GM19152, GM19154, GM19221, GM19223, HG01102, HG01103, HG01883, HG02562, HG02563, HG02611, HG03112). LCLs for these samples were obtained from Coriell Institute for Medical Research (Camden, NJ). After culture with BrdU for 24h, formaldehyde fixed nuclei with BrdU incorporation in the G1 cell cycle phase were enriched by FACS after staining with Hoechst 33258 and Propidium Iodide as described (Hanlon et al. 2022). Approximately 80 single-cell Strand-seq libraries were made for each sample with the cellenONE X1 instrument (SCIENION US, Tempe, AZ). Single-nuclei isolation and picoliter to nanoliter reagent dispensing into the wells of a 5184-nanowell array was performed as described with slight modifications (Hanlon et al. 2022; Porubsky, Dashnow, et al. 2025). Briefly, formaldehyde cross-linking of nuclei was reversed by heat and structural nuclear proteins digested by a heat-labile protease. Restriction enzymes AluI and HpyCH4V (New England Biolabs (Canada) Ltd, Whitby, ON) were used for fragmenting genomic DNA and Hemo KlenTaq (New England Biolabs) was used for A-tailing. Fragments were ligated to forked adapters, UV-irradiated, and PCR-amplified with index primers pre-spotted into nanowells before library preparation. Strand-seq libraries were size-selected with AMPure XP beads and 300 to 700 base-pair fragments were gel purified prior to PE75 sequencing on the NextSeq 550 (Illumina, San Diego, CA).

### Generation of pooled Strand-seq datasets

To efficiently generate low-coverage single-cell Strand-seq libraries from hundreds of human LCLs, samples were processed in pools of 40 different human LCL samples from various human populations. The pools were made by Coriell Institute for Medical Research by culturing individual LCLs and mixing equal numbers of viable cells that were frozen in a single pool of 40 samples. Strand-seq libraries were made from single cells from such pools after 24-hr culture with BrdU. After FACS enrichment of nuclei with BrdU, nanoliter library preparation was done in the same way as individual cell lines. Typically around 2000 single-cell libraries were made from each pool of 40 samples. Strand-seq libraries were size selected with AMPure XP beads and fragments were gel purified prior to PE75 (250 to 450 bp) or PE150 (450 to 1000 bp) sequencing on the AVITI (Element Biosciences, San Diego, CA). After demultiplexing of single-cell Strand-seq library reads from individual wells and alignment to the human reference genome, libraries were re-assigned to individual samples within the pool of 40 samples using publicly available reference haplotype SNP data.

### *De novo* genome assembly of selected samples

For this project we assembled three individuals affected by the 22q11.2 A-D deletion along with a known transmitting parent. In addition, we assembled a family trio (mother - NA10384, father - NA10383, child - NA10382) obtained from Coriell where the child has a *de novo* A-D deletion at 22q11.2 (**Supplementary Table 2**). All above-mentioned samples were assembled using a combination of HiFi and UL-ONT reads using hifiasm (v0.19.5) assembler with the default parameters. Previously reported LCRA-D inversion in HG01888 was assembled using HiFi reads only with hifiasm (v0.16.1) assembler with the default parameters. The other LCRA-D inversion carrier (NA19315) was assembled using a combination of HiFi and UL-ONT reads using hifiasm (v0.19.5) assembler with the default parameters.

### Subsetting a *de novo* assembly sequence to a region of interest (ROI)

In total, we evaluated the completeness of 220 phased genome assemblies generated as part of the HPRC (86 haplotypes), HGSVC (130 haplotypes) (Liao et al. 2023; G. A. Logsdon et al. 2024), and four haplotypes from samples (HG01888 and HG03471) that carry inversion polymorphisms over the 22q11.2 region. Of these, 35 HPRC, 96 HGSVC, and four inverted haplotypes were unrelated and assembled in a single continuous contig, passing our assembly quality validations (Vollger et al. 2019) (**Supplementary Fig. 2**). In order to extract FASTA sequence from a 22q11.2 region (chr22:18000000-23000000) in T2T-CHM13 coordinates, we aligned all available human assemblies from the HPRC and HGSVC to the T2T-CHM13 (v2) reference using minimap2 (v2.24) with the following parameters: ‘`-x asm20 –secondary = no -s 25000`’. Next, we used rustybam (v0.1.33, 10.5281/zenodo.8106233) and its functionality called ‘liftover’ in order to subset alignments in a PAF file to a desired region. Then we used such subsetted PAF files to extract the query FASTA sequence using the R package SVbyEye (v0.99) (Porubsky, Guitart, et al. 2025) and its function ‘paf2FASTA’. We only reported those assemblies that span the ROI in a single contig while at the same time ends of the contig do not map further than 100 kbp from the region boundaries. To evaluate each assembled haplotype for possible misjoins or collapses, we aligned PacBio HiFi reads back to each haplotype using minimap2 (v2.26) with the following parameters: `-a -I 10G -Y -x map-pb --eqx -L --cs`. Such alignments were then manually evaluated for possible collapses by projecting the most and the second most abundant base at each haplotype as reported previously for NucFreq analysis (Vollger et al. 2019). Any haplotype showing an extended region where there is an observable frequency of the second most abundant base was rejected from further analysis. In total, we selected for further analysis 135 human haplotypes (of which two carry an inversion), including extra haplotypes of the NHPs (n=12), *de novo* assemblies of clinical samples (four families), and extra assembly of A-D inversion in NA19315 (**Data availability**).

### Definition of continuous LCR blocks

To define continuous LCR regions (or SD blocks) in T2T-CHM13 (v2) coordinates, we took available SD annotation from: https://s3-us-west-2.amazonaws.com/human-pangenomics/index.html?prefix=T2T/CHM13/a <u>ssemblies/annotation/</u>. Next, we subsetted the SD to our defined ROI (see above). To get a continuous set of LCRs, we collapsed gaps between consecutive SD regions that are 25 kbp in size or shorter. Then, we took only continuous blocks that contain at least 10 SDs and are 10 kbp in size and larger (**Supplementary Table 3**). To define LCR boundaries per haplotype, we first aligned each haplotype to itself using minimap2 with the following parameters: `-DP -k19 -w19 -m200 -c --eqx`. Resultant self-alignments in PAF were read using the SVbyEye (Porubsky, Guitart, et al. 2025) function ‘readPaf’. Such alignments were filtered using the SVbyEye function ‘filterPaf’ with the following parameters: `min.mapq = 0, is.selfaln = TRUE, min.align.len = 10000, min.selfaln.dist = 20000`. Continuous alignments were converted into genomic ranges such that any gaps 50 kbp and shorter were collapsed to create continuous ranges. Finally, we assigned to each continuous LCR an identifier (A-E) based on the overlaps with T2T-CHM13 LCR annotation mapped to each haplotype.

### Clustering LCRA by size

We took previously defined LCRA sizes (n=133), as described in ‘Definition of continuous LCR blocks’, and clustered them by their size using the R package ClusterR and its function ‘KMeans_rcpp’ into 15 clusters. The number of clusters was defined by visual observation of the LCRA size distribution. We then ran the ‘KMeans_rcpp’ function with the following parameters: `clusters = 15, num_init = 10000, max_iters = 1000, initializer = ’kmeans++’`.

### Defining duplicon annotation for each haplotype

We defined the duplicon structure of each human haplotype (n=135), including haplotypes from clinical samples and NHPs, using DupMasker (v4.1.2-p1) (Jiang et al. 2008). Specifically, we used the Rhodonite pipeline, available on GitHub (https://github.com/mrvollger/Rhodonite), to run separately on each selected and validated haplotype as described in ‘Subsetting a *de novo* assembly sequence to a region of interest.’

### Clustering LCRA by DupMasker annotation

We started by extracting FASTA sequences from the defined LCRA boundaries as described in ‘Definition of continuous LCR blocks.’ Next, we ran the DupMasker (Jiang et al. 2008) tool on extracted FASTA sequences to obtain boundaries of the ancestral duplicon units, which altogether represent a DupMasker annotation for each LCRA haplotype (n=133). Then we assigned each unique duplicon unit, as defined by DupMasker, a unique numeric identifier. This identifier is either a positive or a negative number in order to reflect forward and reverse oriented duplicons, respectively. We compared DupMasker annotation for all possible pairs of 133 haplotypes as pairs of numeric codes. The distance between each pair of numeric codes was calculated using the ‘seq_dist’ function from the stringdist (v0.9.12) R package with the parameter ‘method’ set to ‘osa’. All pairwise distances are organized into the distance matrix, which is then used to construct the UPGMA tree using the function ‘upgma’ from the R package phangorn (v2.11.1). By visual inspection of the clustered tree, we arrived at 50 haplogroups reported by the ‘cuttree’ function of the base R package stats.

### Definition of shared repeat segments between LCRA and D

Previously a segment of ∼160 kbp in size has been defined as a repeating unit constituting LCRA and D (Guo et al. 2016; Vervoort et al. 2024). We defined this segment using a complete sequence (T2T-CHM13 reference) of the 22q11.2 region (chr22:18000000-23000000). We aligned this sequence to itself using minimap2 (v2.24) using the following parameters: `-x asm20 -c --eqx -DP -m200`. We then extracted the sequence defined by the self-alignment boundaries (three copies in T2T-CHM13) and constructed the multiple sequence alignment (MSA) using MAFFT (v7.525) with parameter `--auto`. Starting from the MSA, we defined the longest continuous alignment with gaps ≤500 bp (**Supplementary Fig. 5a-b**). We then collapsed this region of the MSA into a consensus sequence of ∼155 kbp using ‘ConsensusSequence’ of the DECIPHER (v3.2.0) R package with the following parameters: `threshold = 0.5, minInformation = 0.5, includeNonLetters = TRUE`. This was followed by self-alignments of the consensus sequence to itself as described above. With this we defined ∼25 kbp inverted repeats at the flanks of our consensus sequence with a unique 105 kbp region in the middle (**Supplementary Fig. 5c**). We report 105 kbp and 25 kbp long consensus sequences as common repeat segments present in both LCRA and D. We found these sequence units to overlap well with previous definitions (Guo et al. 2016; Vervoort et al. 2024) of a 160 kbp repeat while adding more resolution as to which parts are unique and which are repeated (**Supplementary Fig. 5d**).

In addition to the 155 kbp long repeat unit, we also observed another frequently occurring repeat segment of about 28 kbp. We initially defined this segment in T2T-CHM13 at position (chr22:18879899-18907599). We mapped this sequence back to T2T-CHM13 to find all locations of this sequence using minimap2 (v2.24) with the following parameters: `-x asm20 -c --eqx --secondary=yes`. We then extracted the sequence from all locations in the T2T-CHM13 constructed MSA using R package DECIPHER (v3.2.0) and its function ‘AlignSeqs’. Subsequently, we report the consensus sequence of ∼28 kbp in size as described above. Lastly, we defined ∼120 kbp repeat unit observed in five copies in sample NA19129 (haplotype 1; four copies in LCRA and one copy in LCRD). As described above, we again constructed an MSA using DECIPHER (v3.2.0) and exported a consensus sequence of about 120 kbp in length. See **Data availability** for access to all consensus FASTA files for all common repeat units (25 kbp, 28 kbp, 105 kbp, and 120 kbp). Having consensus repeat units for 25, 28, 105, and 120 kbp long segments, we mapped them back to each 22q11.2 haplotype (n=135) using minimap2 (v2.24) with the following parameters: `-x asm20 -c --eqx --secondary=yes -E1,0 -p 0.75` in order to obtain their coordinates in each haplotype. Note that in the case of 120 kbp repeat unit we excluded partial alignments smaller than 100 kbp (**Supplementary Fig. 7**).

### Evolutionary history of common repeat segments

The evolution of repeat segments was analyzed with additional NHPs, including the six great apes genomes (Yoo et al. 2025) and a macaque genome (Zhang et al. 2025). The phylogeny of repeat segments was analyzed in two ways: first with the consensus sequence of each repeat unit (25, 28, and 105 kbp) conserved up to chimpanzee, and secondly, with smaller individual duplicons defined by syntenic alignment blocks compared to macaque (T2T-MFA8v1.0). The alignment to outgroup macaque and NHP genomes was performed by minimap (v2.26) (Li 2018) using the following parameter to locate orthologous copies: `-cx asm20 --secondary=yes --eqx`. MSAs of the orthologous copies of repeat segments were performed using Mafft (v7.525) (Katoh and Standley 2013) with the “--auto” option. For the phylogeny of individual duplicon units, the sequence corresponding to each unit was retained by trimming with Gblocks (v0.91b). A maximum likelihood tree was reconstructed using IQtree (v2.3.6) (Minh et al. 2020) with 1000 bootstrap. The tree was also outgrouped with single-copy orthologs from the macaque genome, and divergence time was estimated by recalibration based on divergence to the siamang and macaque genomes (19.5 and 28.8 MYA, respectively) or chimpanzee genome (6.4 MYA), following the timetree database (Kumar et al. 2022).

### Definition of PATRRs and higher-order duplicon structure

We searched for palindromic AT-rich regions (PATRRs) using the R package Biostring (v2.70.2) and its function ‘matchPattern’ by detecting all positions of ‘AT’ dinucleotides in each haplotype. Then we tiled each haplotype into 100 bp long bins and counted how many AT dinucleotides were in each bin. For each bin we transformed AT dinucleotide counts into z-score and kept only those bins with z-score ≥ 3.29 (CI: 99.9%). After this we collapsed neighboring bins into continuous ranges using the R package GenomicRanges (v1.54.1) and its function ‘reduce’. We further removed all singleton ranges of 100 bp in size. Lastly, we collapsed all neighboring ranges within 10 kbp distance into the final set of PATRRs ranges.

In the next step, we used these PATRR locations to divide the duplicon structure of each haplotype at the PATRR positions. We summarized these into five nonredundant building blocks (**Supplementary Fig. 8**) and searched for the locations of these five building blocks in each haplotype. For this we gave each duplicon a unique number that was either positive or negative for direct and reverse orientation, respectively. We then recorded the best match of each of the five duplicon structures in each haplotype. This was done by measuring the distance between the numeric codes of each building block at all possible positions in each haplotype.

### Mapping annotation on top of each haplotype

Gene annotations used in this study are reported with respect to the T2T-CHM13 reference and were obtained from a file ‘chm13v2.0_RefSeq_Liftoff_v5.1.gff3.gz’ at: https://s3-us-west-2.amazonaws.com/human-pangenomics/index.html?prefix=T2T/CHM13/assemblies/annotation/. We read the annotation file using the readGFF function from the rtracklayer R package (v1.62.0). Next, we extracted the genomic region for all gene boundaries as well as all exons for each gene. We extracted the FASTA sequence from all these regions and mapped them onto each of the haplotypes (n=135). We used minimap2 (v2.24) to map whole gene sequences (including introns) with the following parameters: `-x asm20 -c --eqx --secondary=no`. FASTA sequence from gene exons was mapped with minimap2 (v2.24) using the following parameters: `-x splice -c --eqx--secondary=yes`.

### Analysis of public Iso-Seq datasets

We re-analyzed a collection of 1.4 billion full-length cDNA reads (Iso-Seq) from Dishuck et. al. (Dishuck et al. 2025). Briefly, Iso-Seq reads were extracted based on alignment to 22q11.2 or paralogous regions of T2T-CHM13 reference and re-aligned to each assembled haplotype with minimap2 (Li 2018). We transferred GENCODE v44 gene models to each haplotype with Liftoff (v1.6.3) (Shumate and Salzberg 2021). To generate *de novo* gene models, only reads with ≥99.9% identity to the best matching haplotype that uniquely map to a single paralog (at least 2 bp alignment delta) were used as input to PacBio Pigeon and SQANTI3 (v5.2) (Pardo-Palacios et al. 2024). Tissue expression analyses considered only reads aligning with ≥99.9% identity to their best-matching haplotype.

### Mapping inversion breakpoints

To map inversion breakpoints between LCRA and LCRD in sample HG01888-H1, we first selected a human sample carrying a direct haplotype with a similar DupMasker profile (HG00621-H2). We next aligned the direct haplotype (HG00621-H2) to the inverted haplotype (HG01888-H1) using minimap2 (v2.24) using the following parameters: `-x asm20 -c --eqx --secondary=no`. We projected positions of 25 kbp repeat units and PATRRs to both direct and inverted haplotypes. We then defined putative location of inversion breakpoint as a region encompassing both 25 kbp repeat unit and adjacent PATRR in proximal (LCRA) and distal (LCRD) regions of direct and inverted haplotypes. FASTA sequence from these defined regions was extracted, orientation synchronized, and an MSA constructed using the R package DECIPHER (v2.28.0, function ‘AlignSeqs’). We extracted paralog-specific variants (PSVs) and defined the inversion breakpoint as a region where inverted haplotypes (HG01888-H1) partly match proximal and distal PSVs of the direct haplotype (HG00621-H2) (**Supplementary Fig. 38**). For more details see Porubsky et al. (Porubsky et al. 2022).

### Inversion genotyping within 1KG cohort

For this analysis, we first identified single-nucleotide variants (SNVs) in the assembled inverted (A-D inversion, haplotype 1) and direct haplotype for the 22q11.2 region in sample HG01888. PAV (v2.3.4) was used to call variants with respect to both the GRCh38 and T2T-CHM13 reference genomes. The genotyping was done against a diversity panel of 3,202 1KG samples available for both reference genomes (Byrska-Bishop et al. 2022; Lalli et al. 2025). For genotyping we selected SNV positions present in both PAV as well as the 1KG callset. Then we compared HG01888 haplotypes to all 3,202 samples (6,404 haplotypes) and calculated the fraction of matching SNVs between PAV and the 1KG allele. We selected samples whose fraction of matching SNVs was ≥5 standard deviations (z-score ≥ 5) from the population mean (average fraction of matching SNVs across all comparisons) as candidate samples to carry the inversion haplotype identified in HG01888. This approach was repeated for other A-D inversion carriers (NA19315, HG03239, HG01947, and HG01489) (**Supplementary Fig. 36**) as well as for other inversion breakpoints (A-B, B-D, and C-D) in HG03471, HG01695, HG01175, and HG03139 (**Supplementary Fig. 37**).

### Detection of smaller inversions from minimap2 alignments

In order to detect other common inversion regions outside of large A-D, A-B, and B-D inversions, we searched for inverted alignments in available human haplotypes with respect to a single reference 22q11.2 haplotype. We chose as a reference haplotype HG01890-H1 of African ancestry because this haplotype carries exactly one copy of the 155 kbp repeat unit (105 kbp plus two 25 kbp units) in both LCRA and D. We used minimap2 (v2.24) to align all uninverted human haplotypes (n=133) with respect to HG01890-H1 with the following parameters: `-x asm20 -c --eqx --secondary=no`. We then processed alignment of each human haplotype as follows: We filter out query-to-target alignments that map between non-syntenic LCR blocks. Next, we retained all inverted alignments ≥10 kbp in length. We collapsed gaps between alignments ≤1 kbp into a single region. Then we grouped alignments based on the required reciprocal overlap of ≥60% (**Supplementary Fig. 35**), and for each alignment group we report inversion boundaries as a range that is covered by half and more alignments in a given group. Last, we only retain those inversion ranges supported by at least five and more inverted alignments (**Supplementary Table 1**).

### Analysis of self-aligning regions at 22q11.2

In order to determine what regions and sequences might predispose 22q11.2 to recurrent rearrangements, we aligned each human haplotype (n=133 plus the T2T-CHM13 reference) to itself using minimap2 (v2.24) with the following parameters: `-DP -k19 -w19 -m200 -c --eqx`. To specifically target self-aligning regions between LCRA and D, we selected only alignments that are at least 2 Mbp apart. In addition, to target alignments of the 155 kbp repeat unit we also retained only those alignments at least 120 kbp in length. We then documented, for each haplotype, the total number of aligned bases in forward and reverse orientation, respectively. We used the orientation of self-aligning regions between LCRA and D as a proxy to mark each haplotype as being predisposed to inversion or microdeletion. We marked deletion-predisposed haplotypes as those having alignments only in forward (direct) orientation. In contrast, haplotypes protected against A-D deletion are marked as those that carry only alignments in reverse orientation. Results of this analysis are summarized in **Extended data figure 1**.

### Mapping A-D deletion breakpoints with assemblies

We mapped A-D deletion breakpoints in three family duos (AD009, AD010, and AD013) such that we first created all-versus-all alignments between the proband haplotype that carries the A-D deletion and both parental haplotypes (transmitting parent). We used minimap2 (v2.24) with the following parameters: `-x asm20 -c --eqx --secondary=no`. We located redundant (overlapping) alignments between proband and parental haplotypes at the deletion breakpoint, followed by extracting the FASTA sequence from these regions across all haplotypes (proband and parental haplotypes). We synchronized the orientation of these sequences and constructed an MSA using the R package DECIPHER (v2.28.0, function ‘AlignSeqs’). We then extracted PSVs and defined the deletion breakpoint as a region where the deleted haplotype partly matches PSVs from either parental haplotype. For more details on mapping changepoints in PSV profiles see Porubsky et al. (Porubsky et al. 2022).

### Mapping A-D deletion breakpoints with HiFi reads

We were not able to fully assemble the deleted haplotype in the family trio (GM10382 - proband, GM10383 - father, and GM10384 - mother) thus we decided to map the deletion breakpoint using PacBio HiFi reads. We started by aligning PacBio HiFi reads of the proband to both assembled haplotypes of the transmitting parent (GM10384 - mother) using minimap2 (v2.26) with the following parameters: `-a -I 10G -Y -x map-pb --eqx -L --cs`. We then subset only reads falling within the region of interest (corresponding to 22q11.2 region). While mapping A-D deletion breakpoints, we further kept only reads that overlap LCRA and D in a given sample. We followed by extracting all HiFi alignments with a reported supplementary alignment (SAM flag 2048) and kept those primary and supplementary alignments with reported mapping quality ≥10 and those being >2 Mbp apart from each other. The AD-deletion breakpoints were defined as positions where HiFi reads are cut and continue elsewhere as supplementary alignments.

### Evaluation of high-identity alignments between paralogous copies of 105 kbp repeats

Initially, we determined that putative AD-deletion breakpoints mapped within the 105 kbp long repeat unit. Therefore, to define the likely predisposed and protected haplotype, we searched for all possible mappings between 105 kbp segments from LCRA and D within a single haplotype. To achieve this we masked all sequences within each haplotype that does not correspond to the 105 kbp repeat unit to a string of N’s. We did this for 133 human haplotypes, including four clinical samples (8 haplotypes). Then we aligned each haplotype to itself using minimap2 (v2.24) with the following parameters: `-DP -k19 -w19 -m200 -c --eqx`. We followed by binning each pairwise alignment of the 105 kbp segment between LCRA and D into 1 kbp long bins. For each bin we calculated the percentage of matched base pairs. Last, we selected a continuous set of bins with the percentage of matched base pairs ≥99.5% and collapsed into a single region of high sequence identity. We also tracked the relative orientation of these high-identity regions between LCRA and D as either forward and reverse oriented. We used this information as a proxy to define a continuous scale of likely predisposed (forward orientation) and protected (reverse orientation) haplotypes for AD-deletion. Predisposed haplotypes were defined as those where more than half of all self-alignments are in forward orientation. To define a haplotype as protected, all self-alignments had to be in reverse orientation.

Results of this analysis are summarized in **Figure 5d,e**.

### Data Availability

HGSVC assemblies (n=65) used in this study were reported by Logsdon et al. (G. A. Logsdon et al. 2024). HPRC assemblies (n=47) used in this study were reported by Liao et al. (Liao et al. 2023). Four newly generated HPRC assemblies (HG03239, HG01175, HG03139, and NA20129) are available via NCBI (BioProject ID: PRJNA730823).

An assembly for sample HG03471 was released by Hallast et al. 2023 (Hallast et al. 2023). Newly generated long-read datasets for HG01888 and NA19315 along with corresponding assemblies are available via European Nucleotide Archive (ENA) (BioProject ID: PRJEB91688).

Strand-seq data (for 13 samples) generated for genotyping purposes have been released via ENA (BioProject ID: PRJEB91688).

Pooled Strand-seq dataset has been released specifically for human chromosome 22 and is available via ENA (BioProject ID: PRJEB91688).

Clinical samples will be available via European Genome-phenome Archive (EGA) upon publication.

FASTA sequences for 135 human haplotypes and 12 NHP haplotypes over the 22q11.2 region including FASTA sequences of all four common repeat units is available via Zenodo (doi: 10.5281/zenodo.15790793).

### Author Contributions

D.P. and E.E.E. conceptualization.

V.T., D.D.C., T.Y.L., P.H., J.O.K and P.M.L. Strand-seq data generation and quality control. K.M.M., K.H., N.K. and W.T.H. Long-read data generation and analysis support.

E.S., M.S.S., S.M., A.S., J.B. and J.R.V. Patient data collection. D.P., D.Y., P.C.D. and E.E.E. Data analysis and interpretation. D.P., D.Y. and E.E.E. developed main figures.

D.P., D.Y., P.C.D. and E.E.E. manuscript writing.

E.E.E. supervised experiments and analyses.

## Acknowledgements

We thank T. Brown for edits in the preparation of this manuscript. We also thank Mianne Lee and Trang Nguyen for technical assistance with preparation and sequencing of Strand-seq libraries. This work was supported, in part, by US National Institutes of Health (NIH) grants HG010169, HG010971, and HG007497 to E.E.E. Strand-seq work in the Lansdorp lab is supported by grants from the Terry Fox Research Institute (Program Project Grant #1074), the Canadian Institutes of Health Research (CIHR grants PJT-159787 and PWUZ GR028457), the Canada Foundation for Innovation (CFI grants #40044 and #43153), Genome Canada and Genome BC (project 323ECC) and the BC Cancer Foundation. Last, we would like to acknowledge the National Genome Research Institute (NHGRI) for funding the following grants supporting the creation of the human pangenome reference: U41HG010972, U01HG010971, U01HG013760, U01HG013755, U01HG013748, U01HG013744, R01HG011274, and the Human Pangenome Reference Consortium (BioProject ID: PRJNA730823).

E.E.E. is an investigator of the Howard Hughes Medical Institute. The following cell lines were obtained from the NIGMS Human Genetic Cell Repository at the Coriell Institute for Medical Research: GM01888, GM19315, GM10382, GM10383, GM10384, GM01883, GM02562, GM02563, GM02611, GM03112, GM18870, GM18872, GM19152, GM19154, GM19221, GM19223, GM01102, and GM01103.This article is subject to HHMI’s Open Access to Publications policy. HHMI lab heads have previously granted a nonexclusive CC BY 4.0 license to the public and a sublicensable license to HHMI in their research articles. Pursuant to those licenses, the author-accepted manuscript of this article can be made freely available under a CC BY 4.0 license immediately upon publication.

---

**Extended Data Figure 1:**
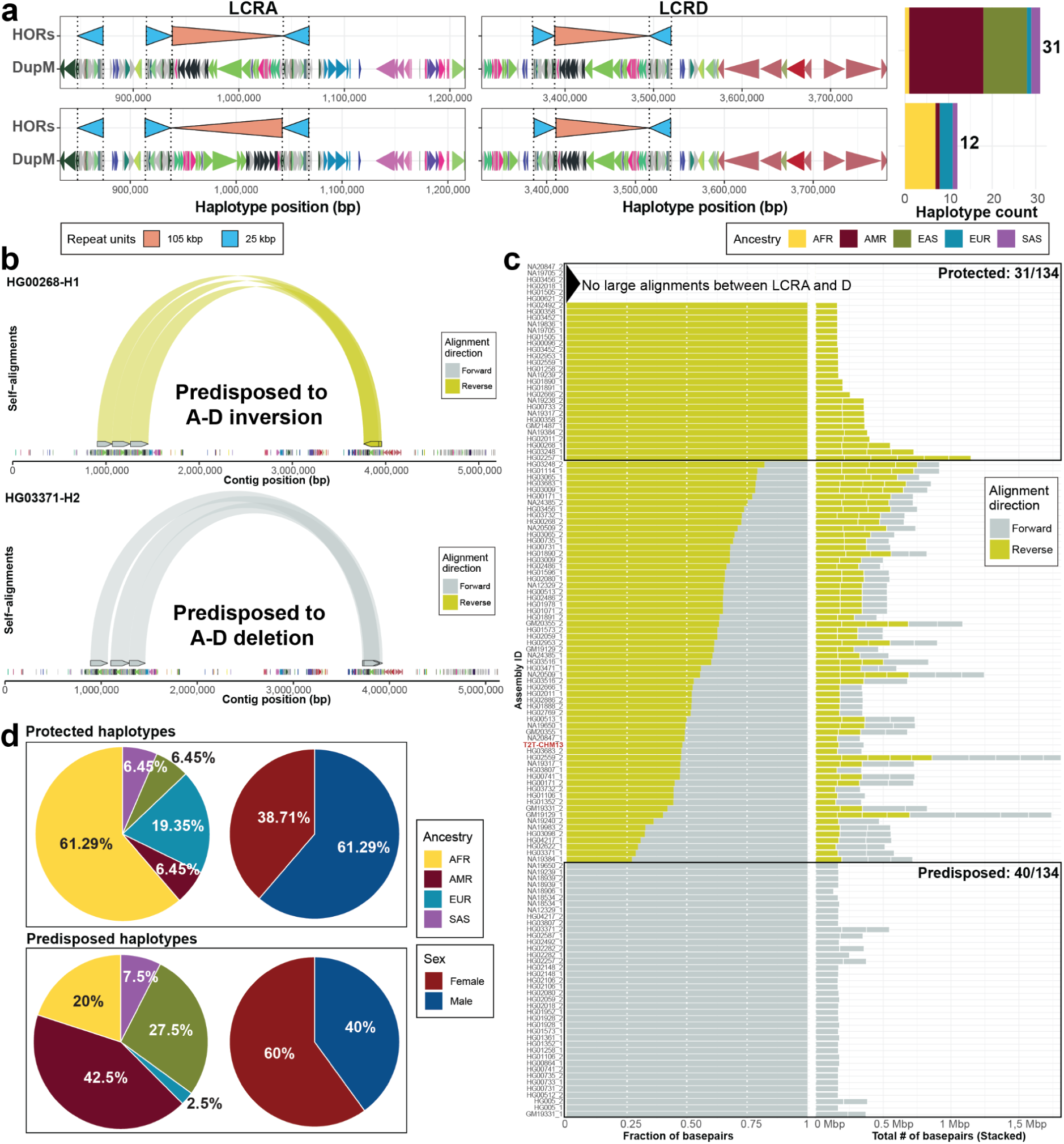
22q11.2 predisposition to recurrent rearrangements. **a**) Summary of human haplotypes (n=43) with a single copy of 105 kbp repeat unit (red) in both LCRA and D and their predisposition to A-D deletion. We show an annotation of 105 (red) and 25 (blue) kbp long repeat units as directional arrowheads. Below there is a duplicon annotation shown as directional arrowheads colored by a unique duplicon ID. Left: there is a stacked barplot showing the haplotype count for the forward (n=31) and reverse (n=12) oriented 105 kbp repeat unit in LCRA. **b**) Arc plot showing the self-aligning regions between LCRA and D for a haplotype predisposed to A-D deletion and a haplotype predisposed to A-D inversion. **c**) Summary of all self-aligning bases (>=120 kbp) between LCRA and D across human haplotypes (n=133 + T2T-CHM13). Left: The fraction of self-aligning bases in forward and reverse orientation. Right: The total number of self-aligning bases in forward and reverse orientation. Each segment represents a single alignment between LCRA and D. Predicted haplotypes protected (n=31) and predisposed (n=40) to A-D deletion are highlighted by black rectangles. **d**) Pie charts summarizing ancestry and sex for protected (n=31) and predisposed (n=40) haplotypes.

