## Supplementary materials for "Population differences of chromosome 22q11.2 duplication structure predispose differentially to microdeletion and inversion"

### Supplementary Figures 1-54 & Supplementary Notes

David Porubsky[1,2], DongAhn Yoo[1], Philip C. Dishuck[1], Nidhi Koundinya[1], Erika Souche[3], William T. Harvey[1], Katherine M. Munson[1], Kendra Hoekzema[1], Daniel D. Chan[4], Tiffany Y. Leung[4], Marta S. Santos[3], Senne Meynants[3], Ann Swillen[6], Jeroen Breckpot[6], Vasiliki Tsapalou[2], Patrick Hasenfeld[2], Jan O. Korbel[2], Peter M. Lansdorp[4,5], Joris R. Vermeesch[3,6], Evan E. Eichler[1,7]

**Affiliations**

1. Department of Genome Sciences, University of Washington School of Medicine, Seattle, WA, USA
2. European Molecular Biology Laboratory (EMBL), Genome Biology Unit, Heidelberg, Germany
3. Laboratory of Cytogenetics and Genome Research, Centre for Human Genetics, KU Leuven, Leuven, 3000, Belgium
4. Terry Fox Laboratory, BC Cancer Research Institute, Vancouver, BC, Canada
5. Department of Medical Genetics, University of British Columbia, Vancouver, BC, Canada
6. Department of Human Genetics, Centre for Human Genetics, University Hospitals Leuven, Leuven, 3000, Belgium
7. Howard Hughes Medical Institute, University of Washington, Seattle, WA 98195, USA

**Corresponding author**

Evan E. Eichler, Ph.D.
Department of Genome Sciences
University of Washington School of Medicine
3720 15th Ave NE, S413C
Box 355065
Seattle, WA 98195-5065


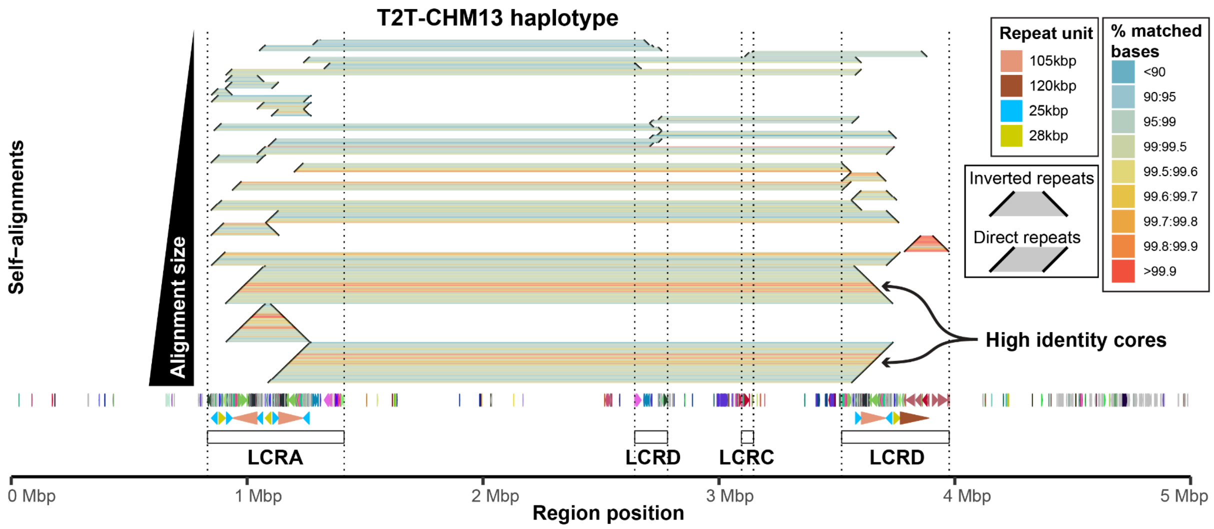


**Supplementary figure 1: T2T-CHM13 self-alignments.**
A “horizontal dotplot” visualization showing self-alignments within the T2T-CHM13 reference for the 22q11.2 region. Alignments (diagonally oriented black lines) are connected by horizontal ribbons colored by percentage of matched bases per 2 kbp long bin. Below is a DupMasker annotation of T2T-CHM13 colored by a unique duplicon ID. Further below is a repeat unit annotation shown as direction pointing arrowheads. Last, there is annotation of LCR regions A-D.

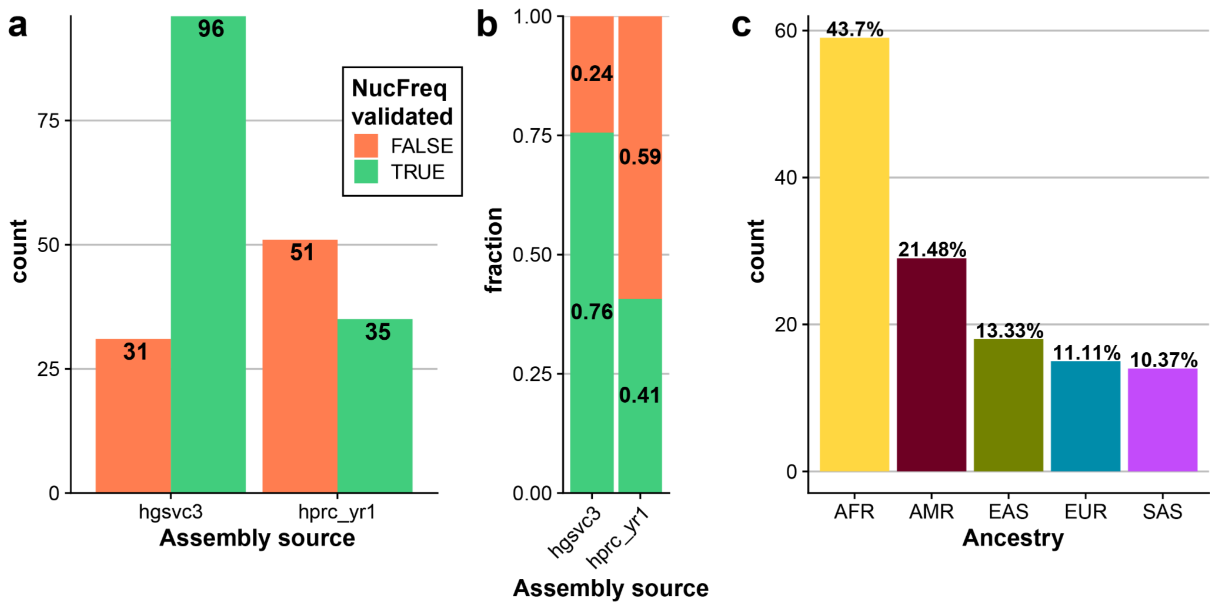


**Supplementary figure 2: Summary of complete assemblies of the 22q11.2 region.**
**a**) Barplot showing the number of complete assemblies (assembled in a single contig) and NucFreq validated assemblies (Valid, TRUE - green; invalid, FALSE - orange) stratified by assembly source (‘hgsvc3’, ‘hprc_yr1’). **b**) A stacked barplot showing the proportion of all evaluated assemblies being assembled in a single contig and validated by NucFreq. **c**) Barplot showing the counts for all NucFreq validated assemblies stratified by ancestry. **Note**: Counts in a) and b) do not include four haplotypes from two samples (HG01888 and HG03471) that carry a large-scale inversion over the 22q11.2 loci; thus, the total count reported here is 131. Counts in c) consider the full panel of 135 diverse haplotypes.

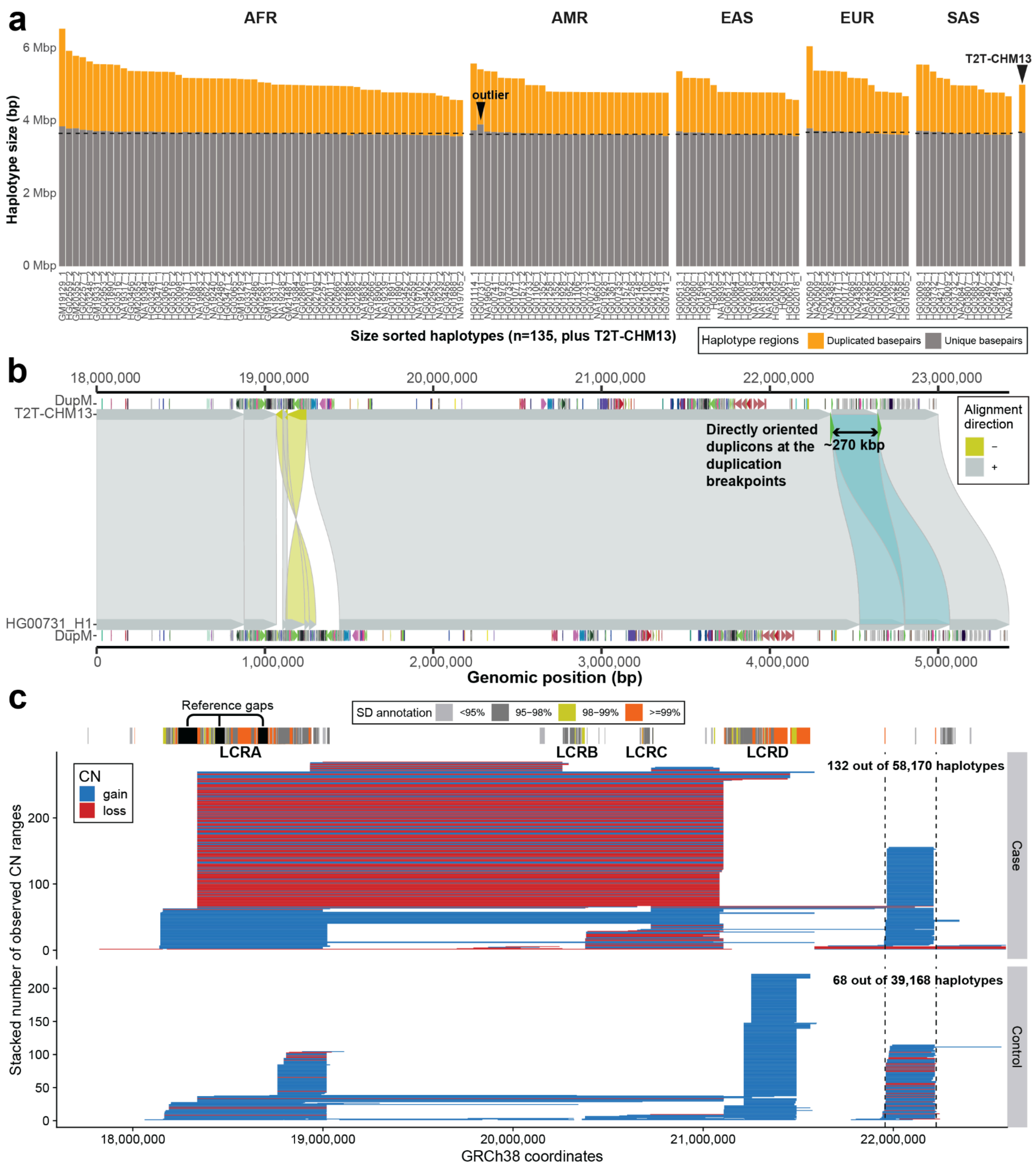


**Supplementary figure 3: Summary of complete assemblies of 22q11.21 region.**

**a**) Barplot showing the proportion of base pairs that are marked as duplicated (orange) and those that are unique (gray) stratified by major ancestries (AFR - African, AMR - American, EAS - East Asian, EUR - European, SAS - Southeast Asian). The median value of unique bases is highlighted by a dashed line per Ancestry. T2T-CHM13 proportions of unique and duplicated bases are shown separately. Each unique haplotype is marked by a sample ID and haplotype-specific (1 or 2) label. A haplotype with an excess of unique bases (HG00731_1) is marked as outlier. **b**) Miropeats-style plot showing alignments (direct - gray, '+' and inverted - yellow, '-') between query (bottom, HG00731-H1) and target (T2T-CHM13) sequences. There is a duplicon annotation ('DupM' track) specific to query and target sequences shown as directional arrowheads colored by unique duplicon ID. The duplicated sequence (~270 kbp) in HG00731 haplotype 1 is marked by light blue colored alignments. As expected, there are directly oriented duplicons (green arrowheads) at the duplication breakpoints supporting a duplication event via NAHR. **c**) Presentation of previously published data of 29,085 cases with developmental delay along with healthy controls (n=19,584) over the 22q11.2 region (Cooper et al. 2011; Coe et al. 2014). Each vertical line represents a defined copy number (CN) region marked either as a loss (red) or gain (blue) of a given region. Only CN changes of 200 kbp and longer are shown. The region of interest of a duplication defined in panel b) is highlighted by vertical dashed lines. We observe the CN gain in our region of interest among 132 cases and 68 controls.

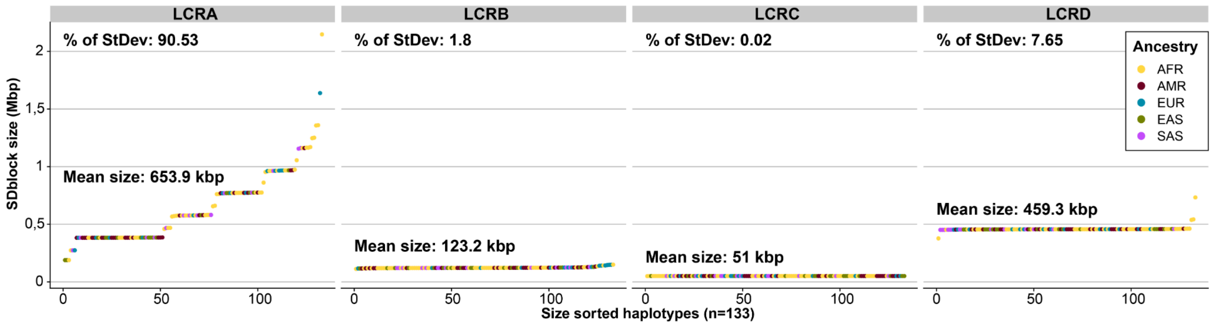


**Supplementary figure 4: Summary of LCRA-D sizes.**
Distribution of LCR sizes per haplotype (n=133) colored by Ancestry (AFR - African, AMR - American, EAS - East Asian, EUR - European, SAS - Southeast Asian). For LCRA-D we also report a mean size as well as percentage of standard deviation per LCR ('% of StDev').

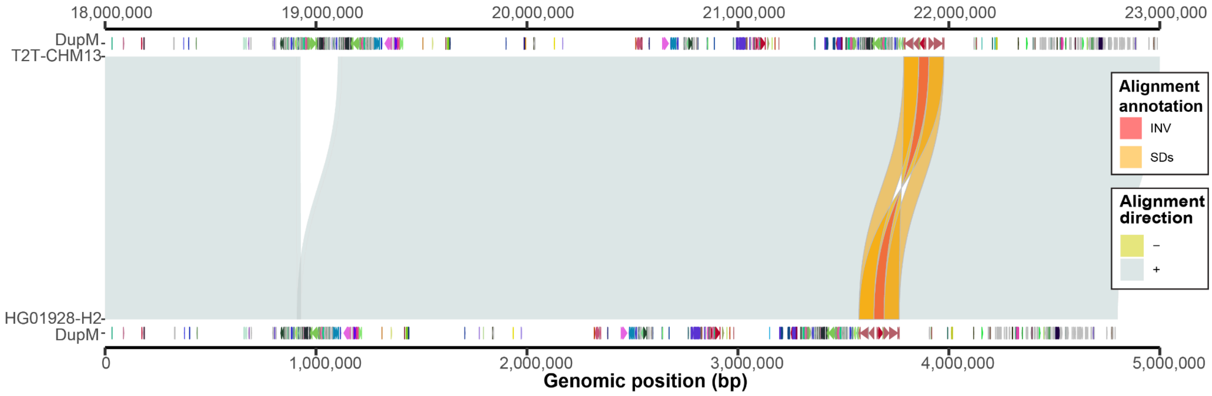


**Supplementary figure 5: Distal inversion in SD block D.**
Miropeats-style plot showing alignments (direct - gray, '+' and inverted - yellow, '-') between query (bottom, HG01928-H2) and target (T2T-CHM13) sequences. There is a duplicon annotation ('DupM' track) specific to query and target sequences shown as directional arrowheads colored by a unique duplicon ID. Inverted sequence in HG01928-H2 is highlighted as red colored alignment (48.65 kbp in size) while flanking inverted duplications (~71 kbp in size) are visible as orange colored alignments between query and target sequences.

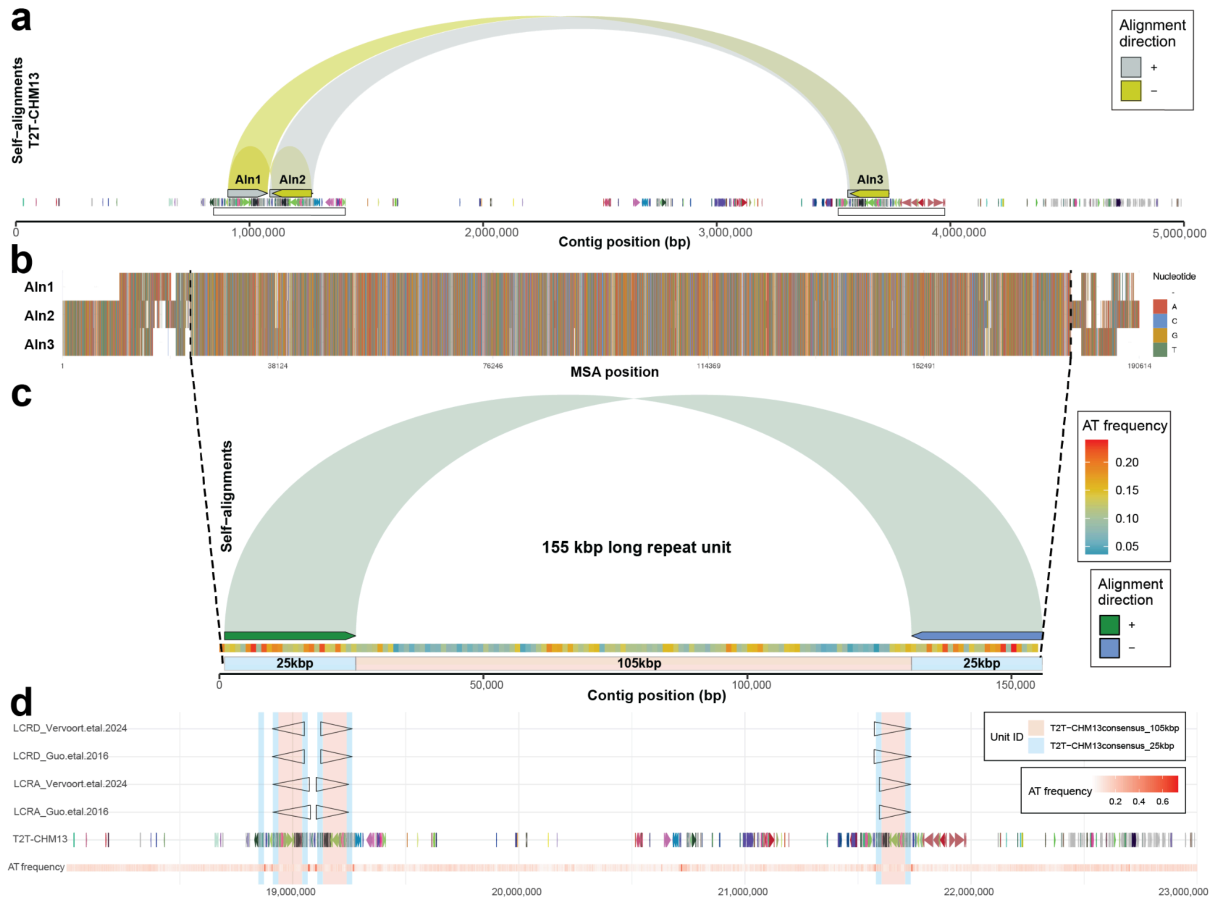


**Supplementary figure 6: Definition of common repeat unit within T2T-CHM13 at 22q11.2.**
**a**) Visualization of long (≥110 kbp) self-alignments within the T2T-CHM13 reference shown as gray and yellow arcs for direct and reverse oriented alignments, respectively. Below is a DupMasker annotation shown as directional arrowheads colored by a unique duplicon ID. Further below are white rectangles delineating LCRA (left) and LCRD (right). **b**) Multiple sequence alignment (MSA) between self-aligning regions (Aln 1-3) from (a). Vertical dashed lines highlight a region that can be continuously aligned between the three sequences including gaps ≤500 bp. **c**) Self-alignment of the consensus sequence (~155 kbp), obtained from the MSA, is shown as an arc connecting flanking 25 kbp long regions in inverted orientation. Below is a binned (bin size 1 kbp) frequency of AT dinucleotides to show there are no extended regions of AT richness. Lastly, we show colored bars highlighting 105 kbp long core and 25 kbp long flanking inverted repeats that compose the 155 kbp long common repeat unit. **d**) Mapping of previously defined common repeat units (Guo et al. 2016; Vervoort et al. 2024) (~160 kbp) to the T2T-CHM13 reference are shown as empty arrowheads. Below is a DupMasker annotation of T2T-CHM13 shown as directional arrowheads colored by a unique duplicon ID. At the bottom there is a binned (bin size 1 kbp) frequency of AT dinucleotides. Mappings of our redefined repeat unit (~155 kbp) are shown as vertical rectangles colored by 105 kbp long core (red) and 25 kbp long inverted repeats (blue).
Note: See methods section ‘Definition of shared repeat segments between LCRA and D’ for more details.

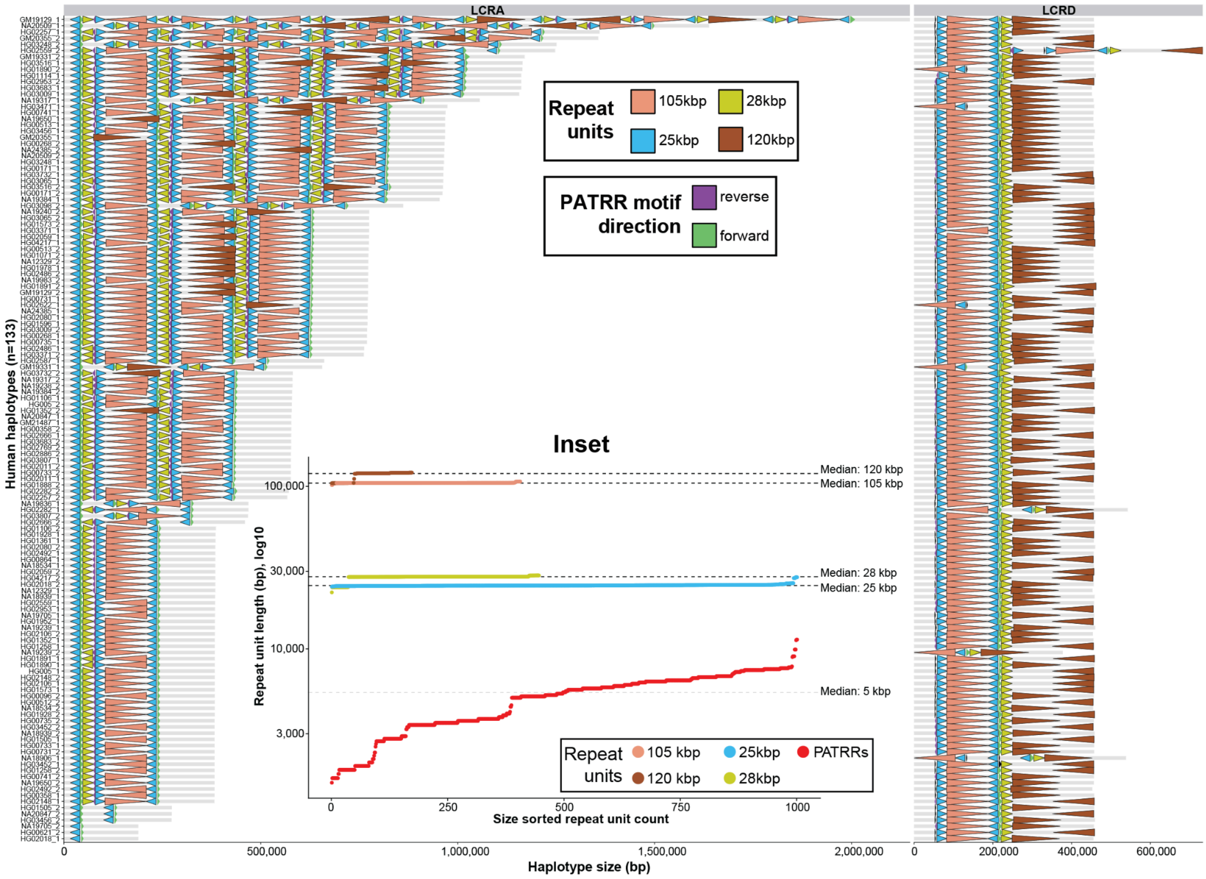


**Supplementary figure 7: Higher order structure of LCRA and D among human haplotypes (n=133).** Higher order units in this plot were defined based on self-aligning regions between LCRA and D at base-pair resolution using the T2T-CHM13 haplotype as a reference to define sequence of each unit (25, 28, 105 and 120 kbp in length). Here, each repeat unit in each human haplotype (in rows) is shown as an arrowhead to reflect the directionality of each repeat in LCRA and D (in columns). **Inset**: Median size across all paralogous copies of each repeat unit in LCRA and D.

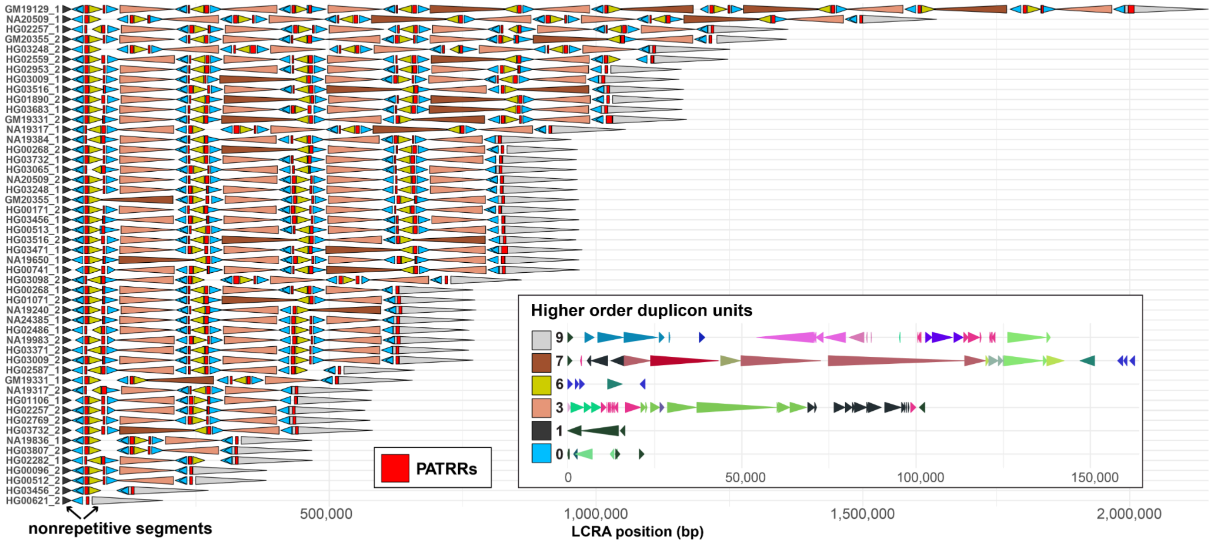


**Supplementary figure 8: Higher order duplicon structure of each representative haplotype of LCRA (n=50).**

Higher order structure was defined based on the repeated duplicon units as defined by DupMasker (**Methods**). We defined a total of six nonredundant units of which four (unit ID: 0, 3, 6, and 7) correspond to common repeats units between LCRA and D as defined in **Supplementary fig. 7**. The remaining two units (shades of gray, nonrepetitive segments) are the flanking units of each LCRA haplotype and are not present in LCRD. We also highlight positions of palindromic AT-rich regions (PATRRs) as red rectangles in each haplotype. **Inset**: Representation of six nonredundant duplicon units.

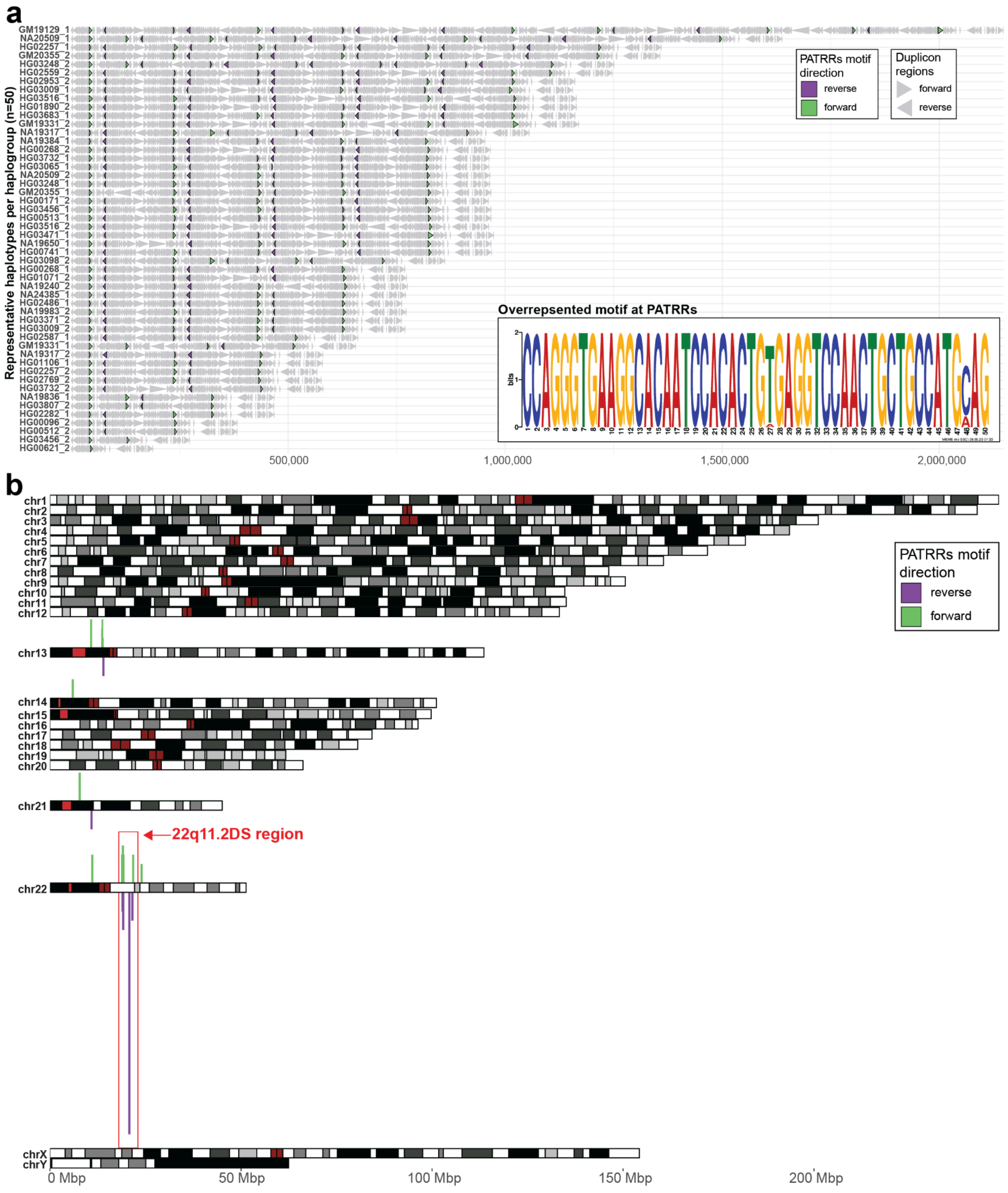


**Supplementary figure 9: Position of PATRRs in each representative haplotype (n=50).**
**a**) DupMasker structure shown as a set of directional arrowheads all colored in gray. Pockets of PATRRs are highlighted as arrowheads colored by the motif orientation (see inset) (forward - purple, reverse - green) within each haplotype as defined using a FASTA sequence for each haplotype (**Methods**). **Inset**: Sequence logo of the overrepresented motif defined among PATRRs. **b**) Frequency of PATRR motif sequence (FIMO score ≥ 50) counted in 200 kbp long bins along each chromosome from the T2T-CHM13 reference. Direction of each matched motif is recorded and counted separately as forward (purple) and reverse (green) oriented motifs.

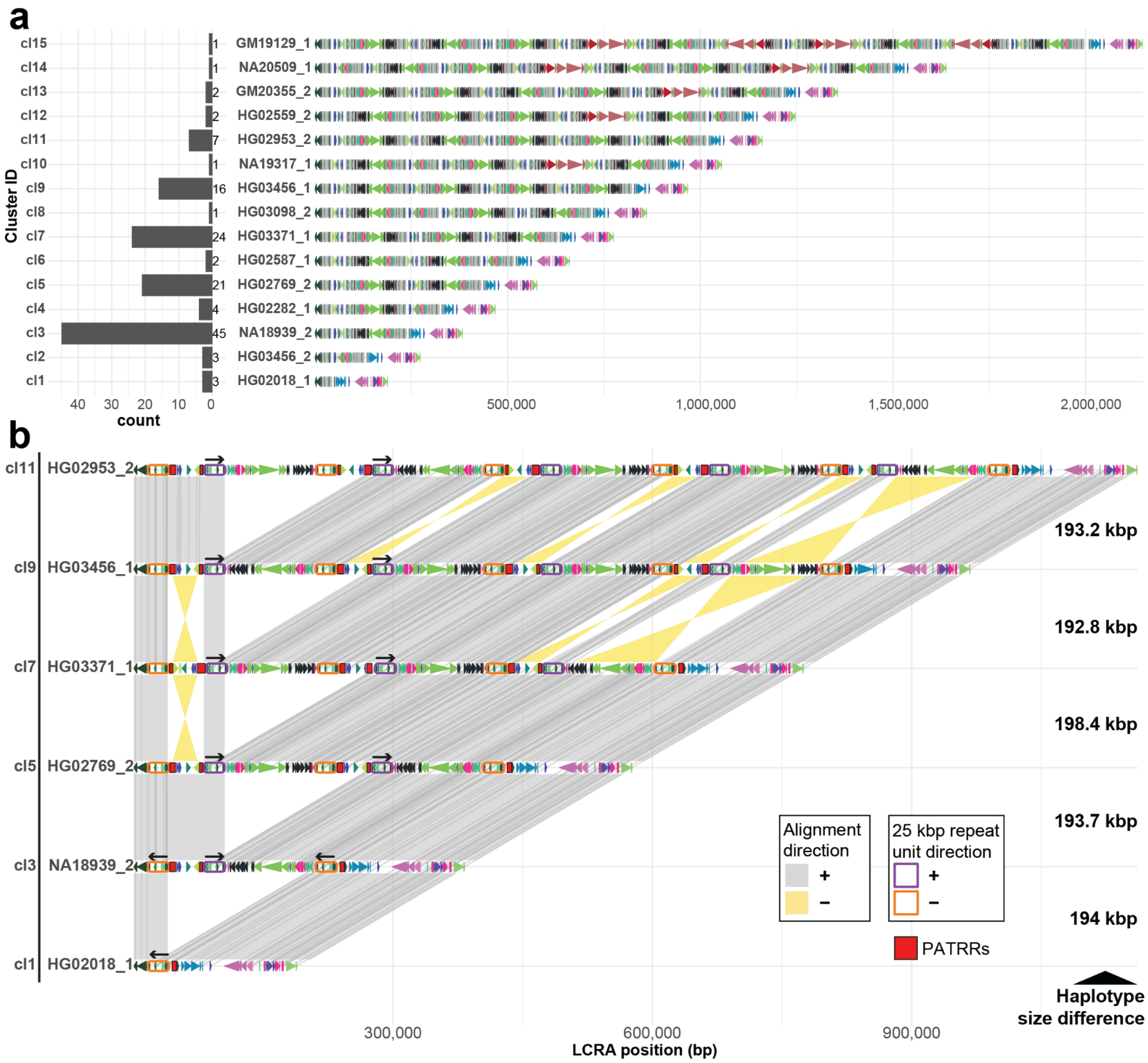


**Supplementary figure 10: Gradual size expansion of SD block A.**

**a**) Left barplot shows a count summary of all 133 haplotypes clustered by their size as shown in Figure 2b. On the right we show DupMasker annotation of manually selected representative haplotypes for each size group from (a). They are shown as directional arrowheads colored by a unique duplicon ID. **b**) Selected haplotypes from (a) that belong to size groups with the highest haplotype count. Their DupMasker structure is reported along with putative alignments between haplotypes. Position and direction of the 25 kbp long repeat unit is shown for direct (purple) and reverse (orange) oriented repeats. PATRR positions are shown as red rectangles. The number in between in each subsequent pair of haplotypes reports their size difference.

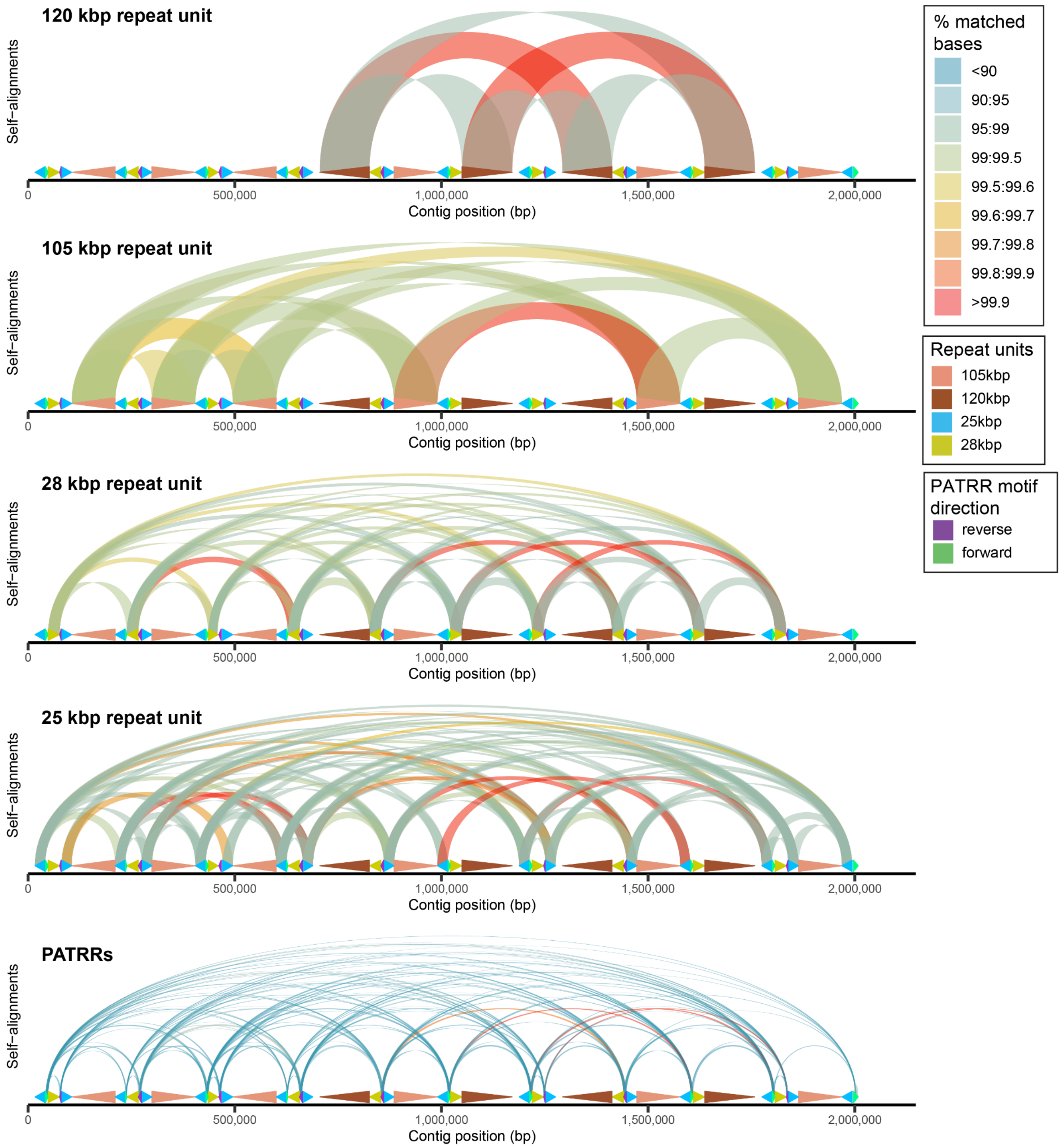


**Supplementary figure 11: Levels of palindrome complexity at LCRA.**
Visualization of alignments between defined repeat units (25, 28, 105 and 120 kbp in size) and PATRRs (~5 kbp in size) for the longest LCRA haplotype in sample GM19129-H1. Each alignment is colored by the percentage of matched bases. There is colored annotation of all repeat units and PATRRs (see legend) with arrowhead orientation reflecting the orientation of each repeat unit and PATRRs within the haplotype.

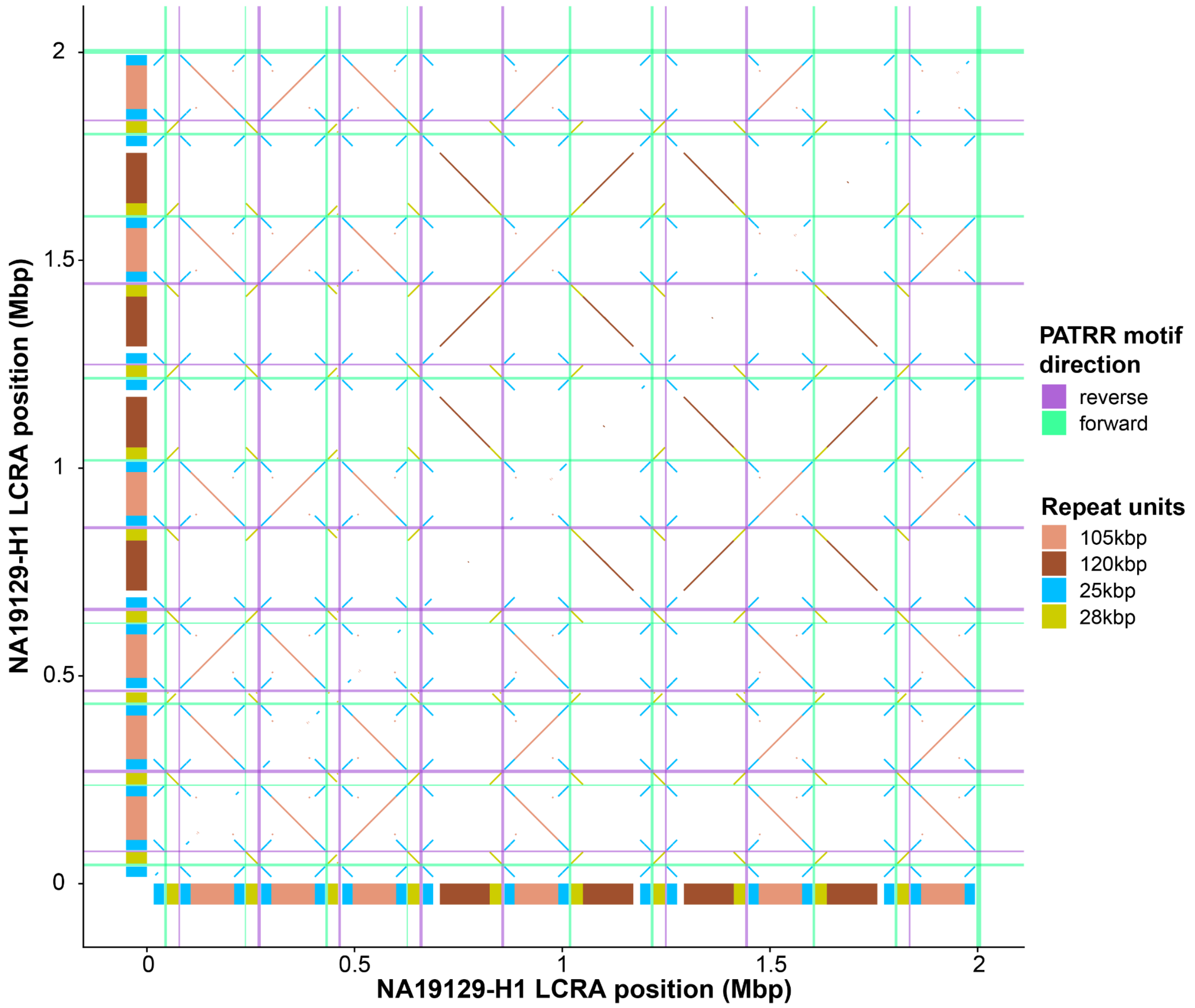


**Supplementary figure 12: Levels of palindrome complexity at LCRA.**
Visualization of alignments between defined repeat units (25, 28, 105 and 120 kbp in size) and PATRRs (~5 kbp in size) for the longest LCRA haplotype in sample GM19129-H1 as an alignment dotplot. Each alignment (diagonal lines) is colored by the repeat unit identity. PATRR positions are shown as horizontal and vertical rectangles colored by PATRR motif direction (see legend).

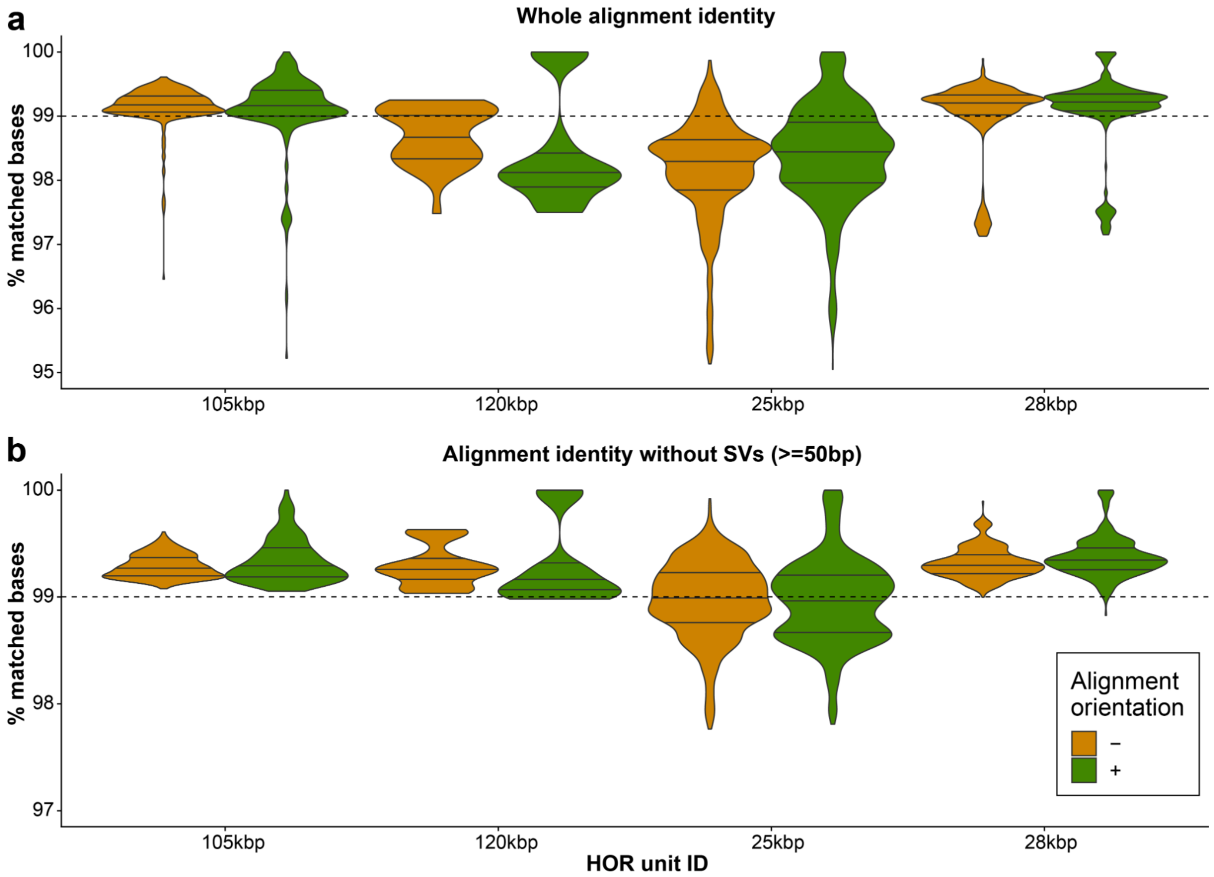


**Supplementary figure 13: Sequence identity of HOR units.**
Distribution of sequence identity (measured as a percentage of matched bases in an alignment) for all alignments between paralogous copies of each HOR unit (25, 28, 105, and 120 kbp) within each human haplotype (n=133). Panels **a**) and **b**) show the alignment identity calculated before and after removal of SVs from identity calculation, respectively. Alignment identity distributions are stratified by the alignment orientation (orange - reverse, '-' and green - direct, '+'). Three horizontal lines within each mark the distribution quantiles (0.25, 0.5, and 0.75).

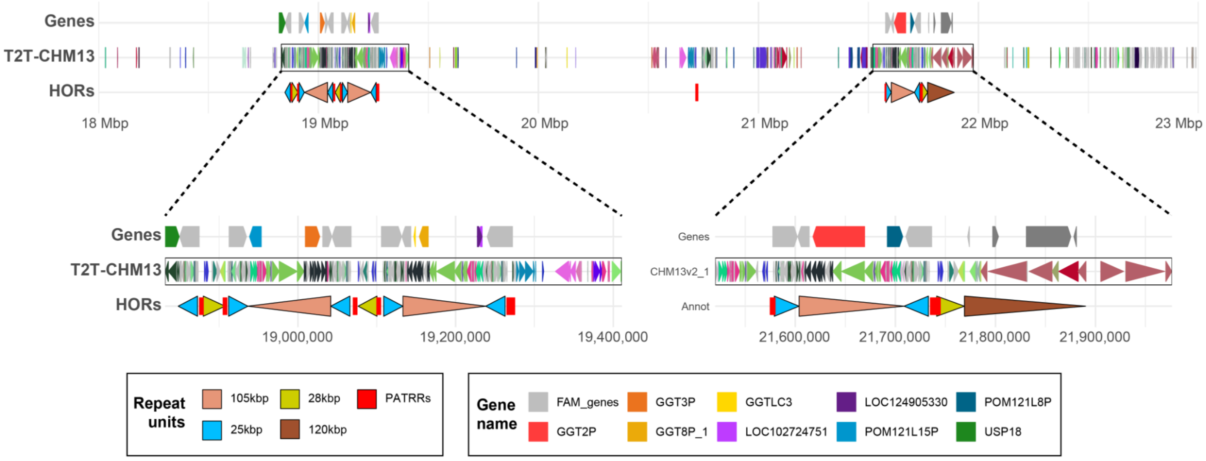


**Supplementary figure 14: Genes overlapping defined repeat units in T2T-CHM13 coordinates.**
From top to bottom row we show gene boundaries colored by gene name. Then there is a DupMasker annotation shown as a set of directional arrowheads colored by a unique duplicon ID. Last, there is a common repeat unit annotation colored by unit identity (25, 28, 105, and 120 kbp) along with PATRR positions colored as red rectangles.

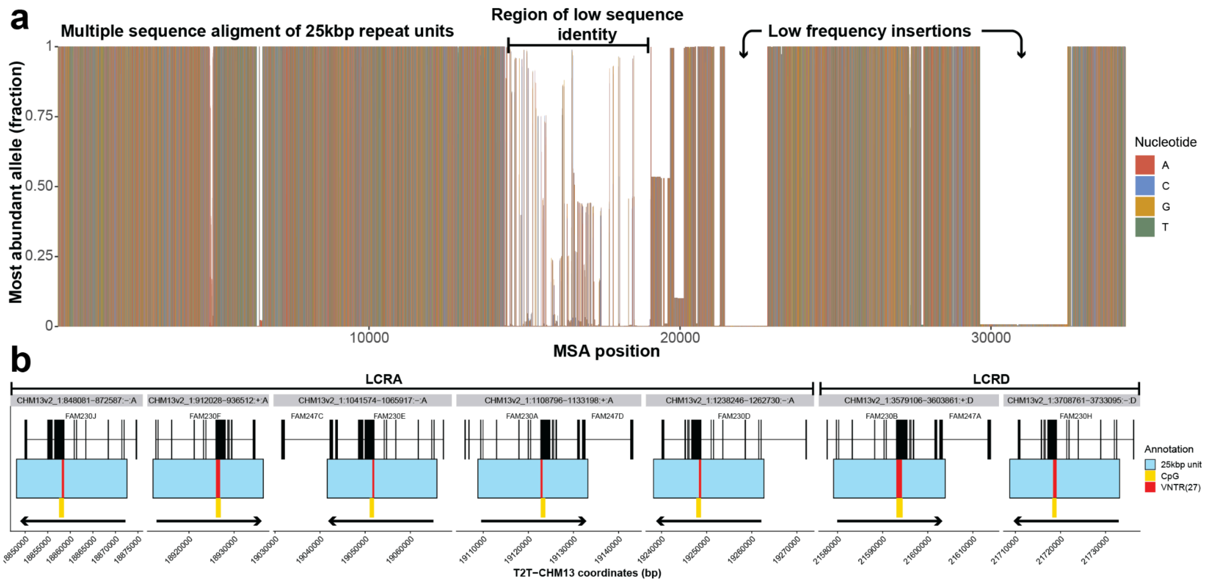


**Supplementary figure 15: Sequence identity and structure of 25 kbp long repeat unit.**
**a**) MSA using all copies (n=1021) of 25 kbp long repeat units among 134 human and 4 nonhuman primate haplotypes (chimpanzee and bonobo). The height of each bar represents the fraction of the most abundant nucleotide (allele) at any given position in the MSA. There are low-frequency insertions in the distal part of the repeat unit but those have only a marginal effect on overall sequence identity among all alignments between copies of this repeat unit. In the middle we observe an extended region of low sequence identity likely caused by the difficulty to align this region. **b**) Visualization of all copies of 25 kbp repeat units (blue rectangles) present in the T2T-CHM13 reference with respect to positions of *FAM230* genes (black rectangles - exons connected by black line). The VNTR position is marked within each repeat unit by a red rectangle followed by the annotation of the known CpG islands in T2T-CHM13 (yellow rectangles). The orientation of each 25 kbp repeat unit is shown by black arrows.

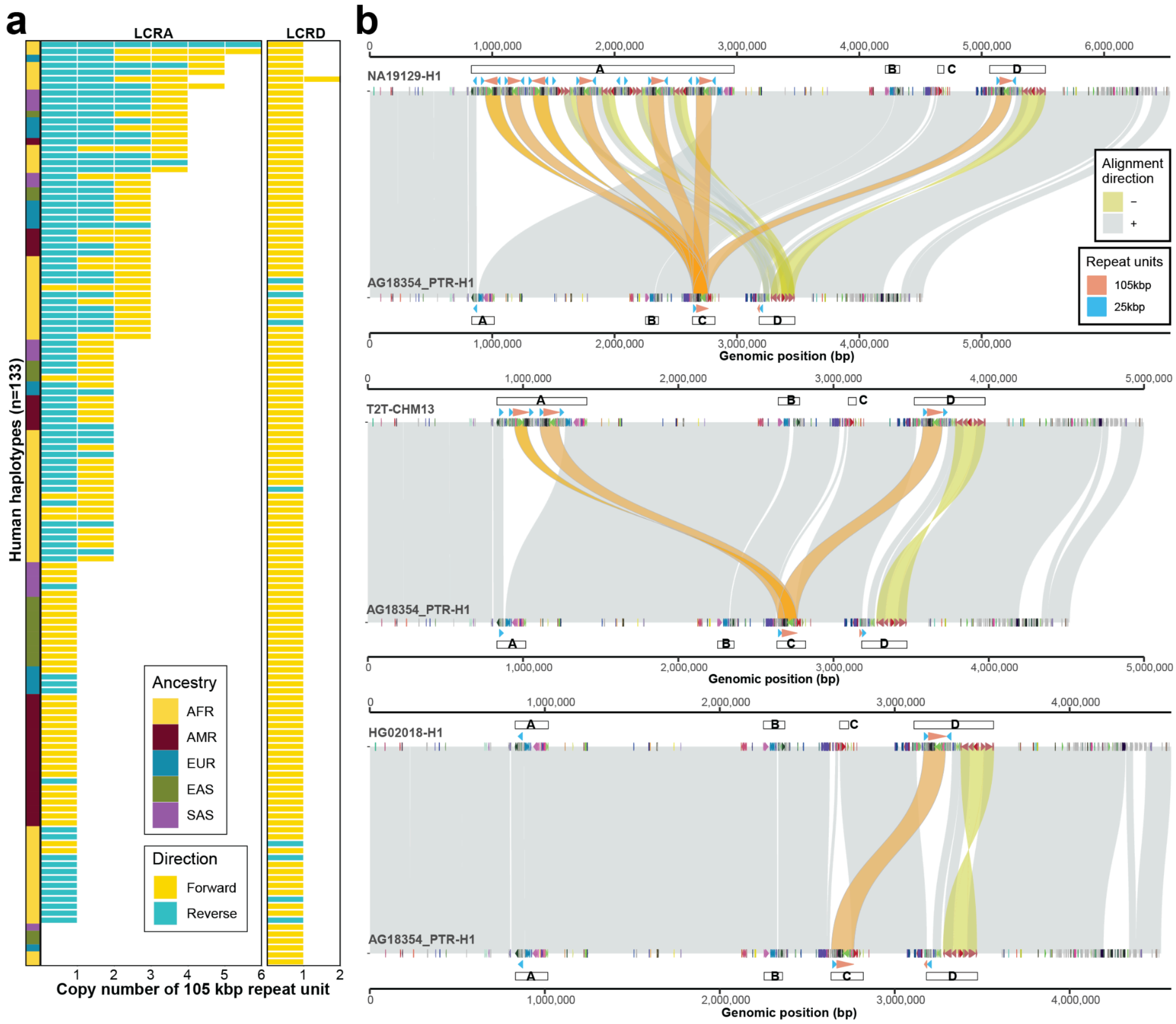


**Supplementary figure 16: Copy number of 105 kbp repeat unit in humans.**

**a**) Counts of main repeat unit (105 kbp) in each human haplotype (n=133) stratified by SD block identity (A or D) and alignment orientation (forward '+' - yellow; reverse '-' - blue). The ancestry of each haplotype is defined in the leftmost panel. **b**) Miropeats-style plot showing alignments (direct - gray, '+' and inverted - yellow, '-') between query (bottom) and target (top) sequences. The top plot shows comparison of chimpanzee haplotype 1 (AG18354_PTR_1, query) to the NA19129 haplotype 1 (target), which is a human haplotype with the highest number of 105 kbp repeat units. The middle plot shows the copy number of the 105 kbp repeat unit in the T2T-CHM13 reference while the bottom shows the haplotype with the least number of copies of the 105 kbp unit (HG02018-H1). Alignments of the 105 kbp repeat unit are highlighted by light orange color. There is a duplicon annotation track specific to query and target sequences shown as directional arrowheads colored by unique duplicon ID. Positions of LCRA-to-D are depicted by white rectangles positioned above the target and below the query sequences, respectively.

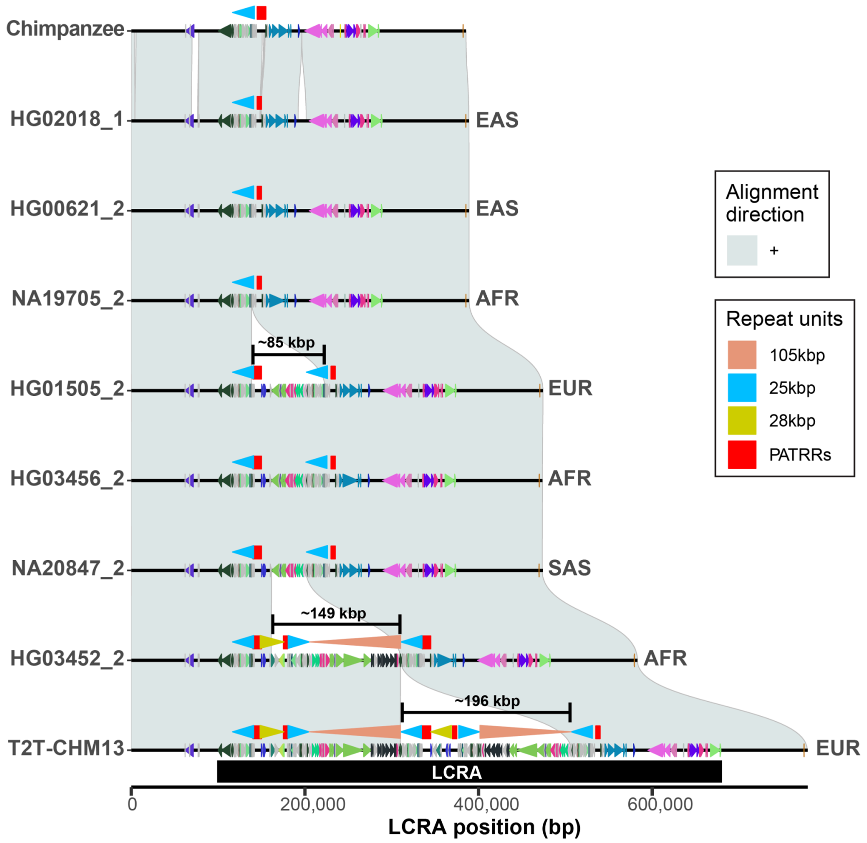


**Supplementary figure 17: Ancestral structure of LCRA in humans.**
Progressive alignments ('+' - directly oriented alignments) between ancestral LCRA structure (chimpanzee) and multiple human haplotypes most similar to the ancestral haplotype, including a human haplotype that carries a single copy of the 105 kbp repeat unit as well as the T2T-CHM13 reference that carry two copies of the 105 kbp repeat unit in LCRA. On top of each haplotype is a DupMasker annotation shown as directional arrowheads colored by a unique duplicon ID. Above each haplotype we show an annotation of various repeat units, namely 25, 28, and 105 kbp repeat units, including PATRR positions.

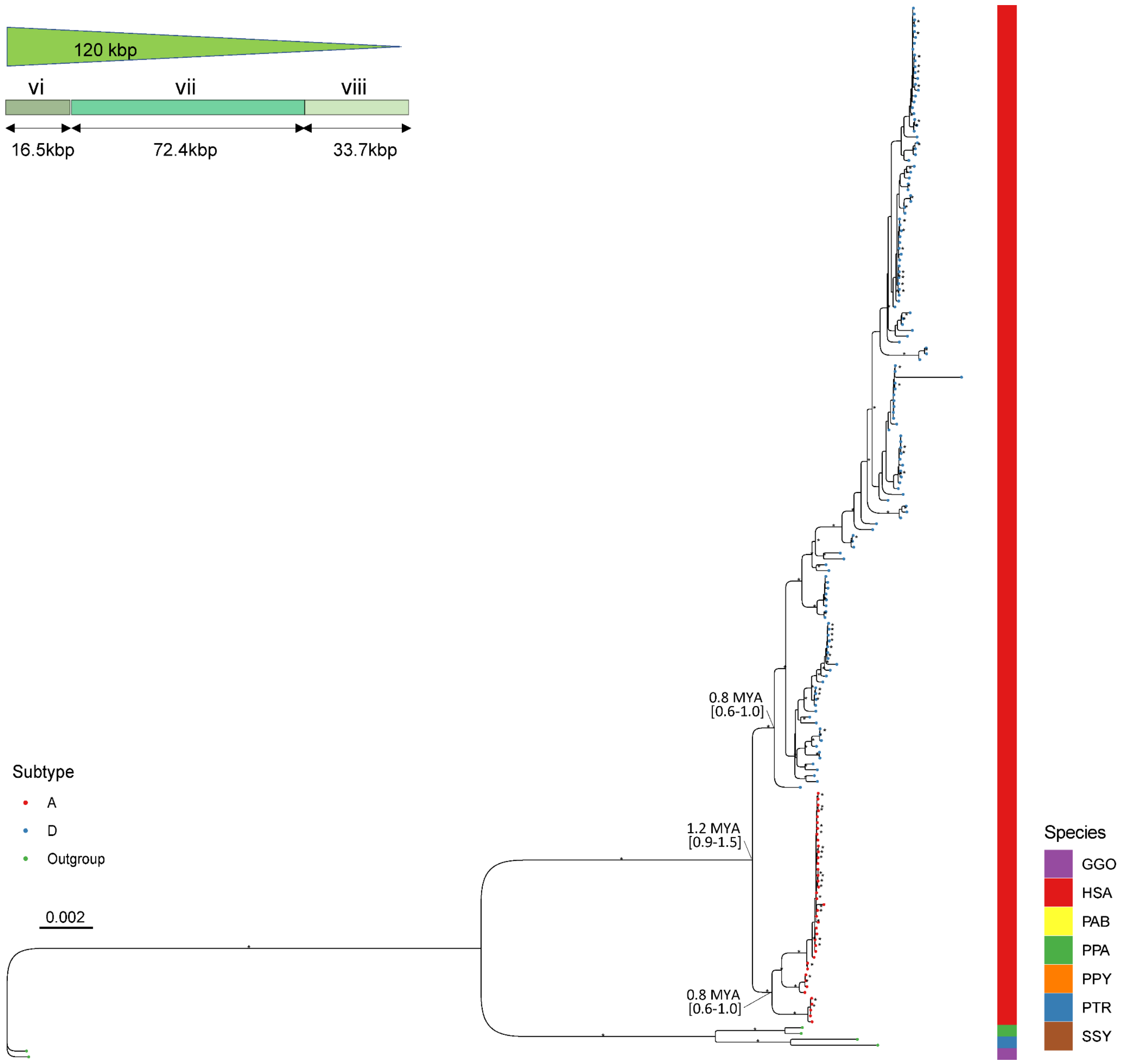


**Supplementary figure 18: Phylogenetic tree of duplicon (vi-viii) of length 120 kbp (n=180).** The tree was outgrouped with gorilla orthologs. The human copies located in LCRA and D are indicated as red and blue, respectively. The corresponding species is indicated on the right annotation track; chimpanzee (PTR), bonobo (PPA), human (HSA), and gorilla (GGO). Notable divergence times among human copies are indicated as million years ago (MYA) with the 95% confidence interval. The bootstrap value of >95 is indicated as an asterisk.

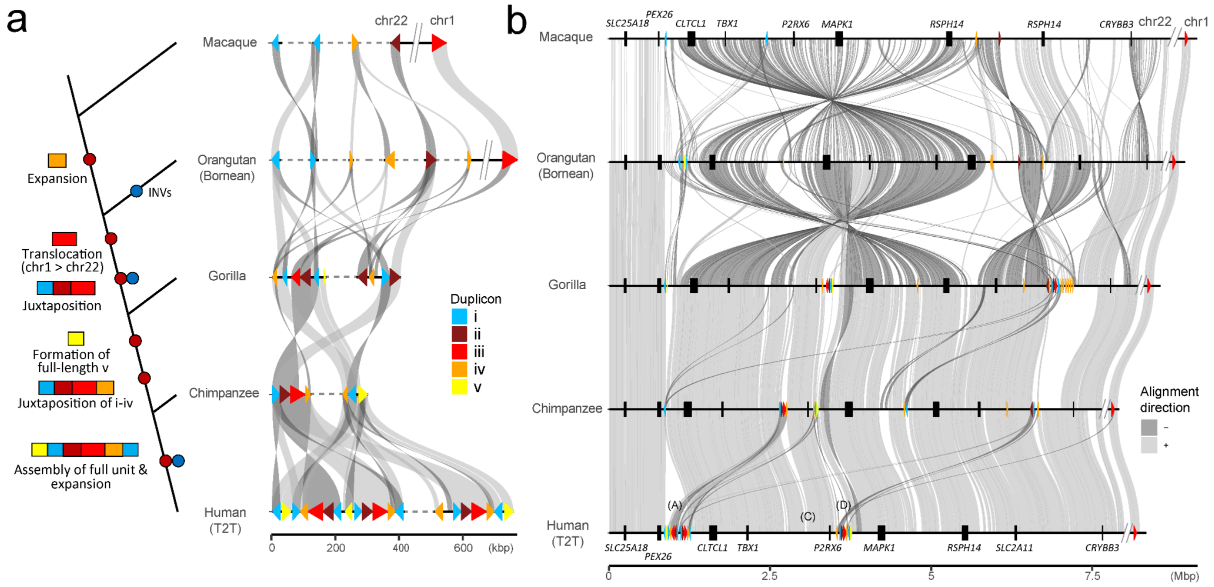


**Supplementary figure 19: Evolutionary model for formation of complex duplication. a**) Schematics of the evolution model, illustrating the emergence of complex duplication observed in human, through multiple steps: 1) expansion of duplicon iv, 2) translocation of duplicon iii from chr1 to chr22, 3) juxtaposition/transposition of complex duplication i-ii and i-iii, 4) formation of full-length duplicon v, 5) juxtaposition of complex duplication i-iv, and 6) formation of full-length human duplication and further expansion of the complex duplication. **b**) Syntenic view of chromosome 22 18-26 Mbp (T2T-CHM13 reference coordinates) region as well as original position of duplicon iii located in chr1. Syntenic genes are indicated along with the duplicons in their respective colors.

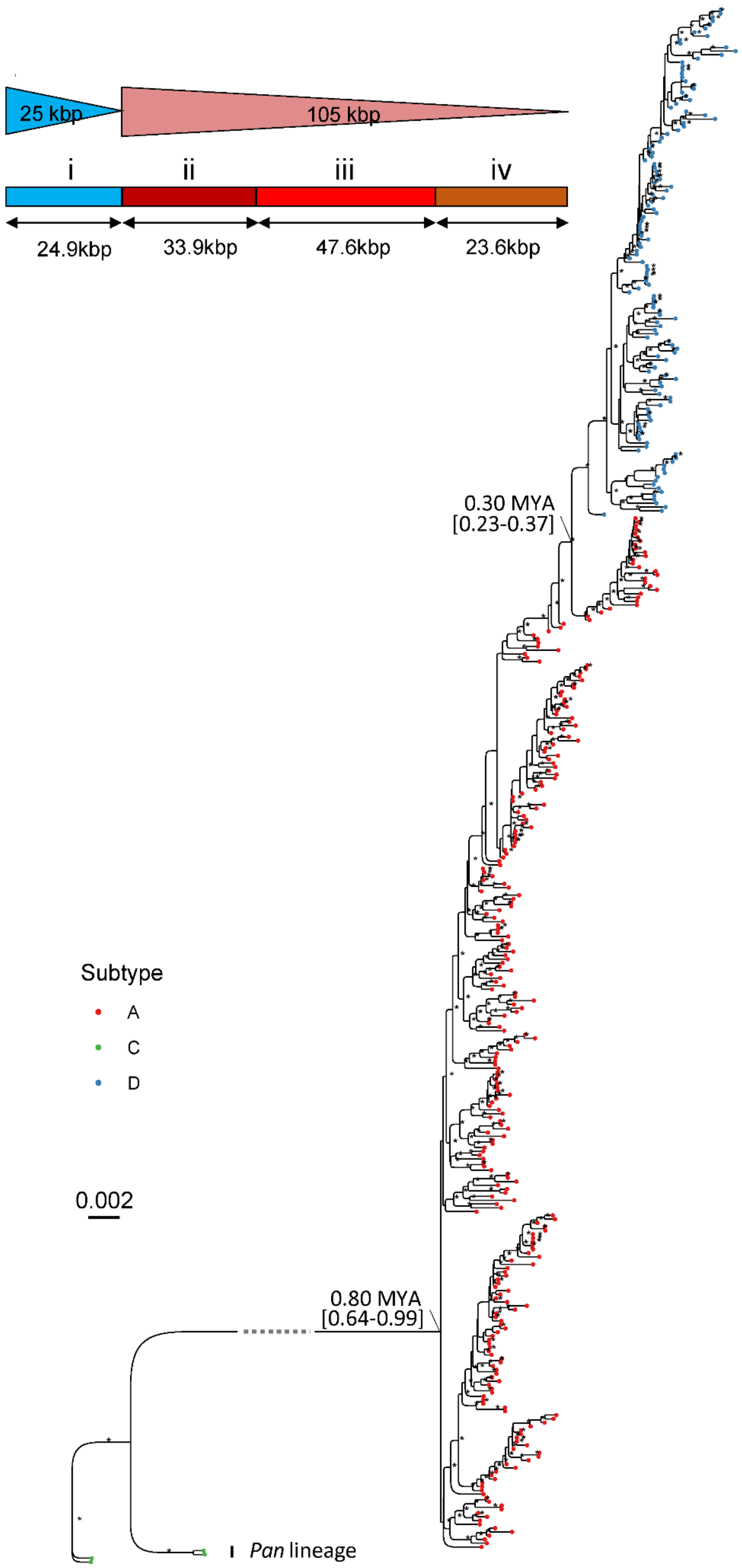


**Supplementary figure 20: Phylogenetic tree of duplicons (i-iv) of length 105 kbp (n=413).** The tree was outgrouped with chimpanzee and bonobo (*Pan* lineage) orthologs. The copies located in LCRA, C, and D are indicated as red, green and blue, respectively. Notable divergence times among human copies are indicated as million years ago (MYA) with the 95% confidence interval. The bootstrap value of >95 is indicated as an asterisk.

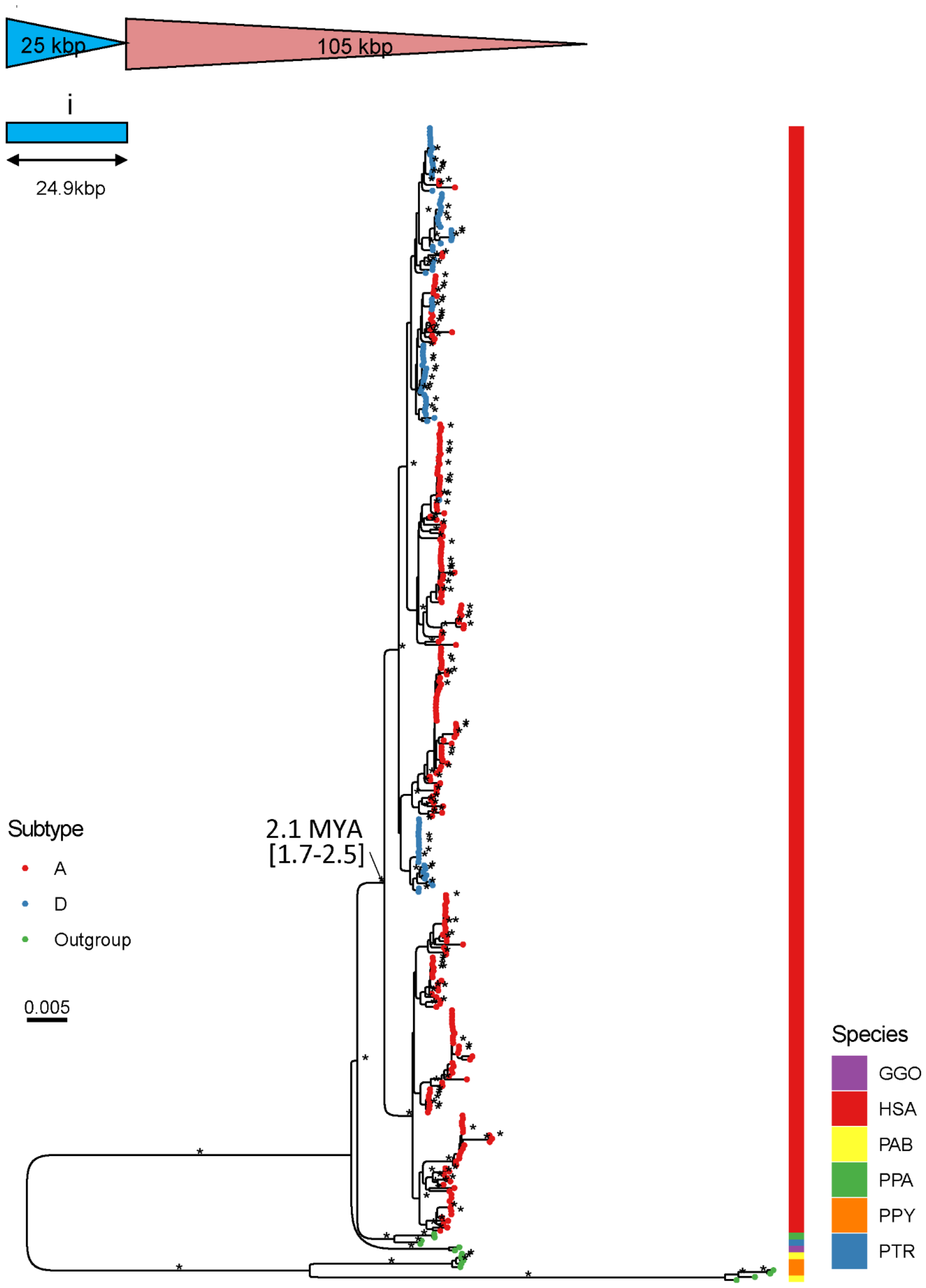


**Supplementary figure 21: Phylogenetic tree of duplicon (i) of length 25 kbp (n=1024).** The tree was outgrouped with macaque and orangutan orthologs. The human copies located in LCRA and D are indicated as red and blue, respectively. The corresponding species is indicated on the right annotation track; chimpanzee (PTR), bonobo (PPA) and human (HSA), gorilla (GGO), Sumatran orangutan (PAB), Bornean orangutan (PPY). Notable divergence times among human copies are indicated as million years ago (MYA) with the 95% confidence interval. The bootstrap value of >95 is indicated as an asterisk.

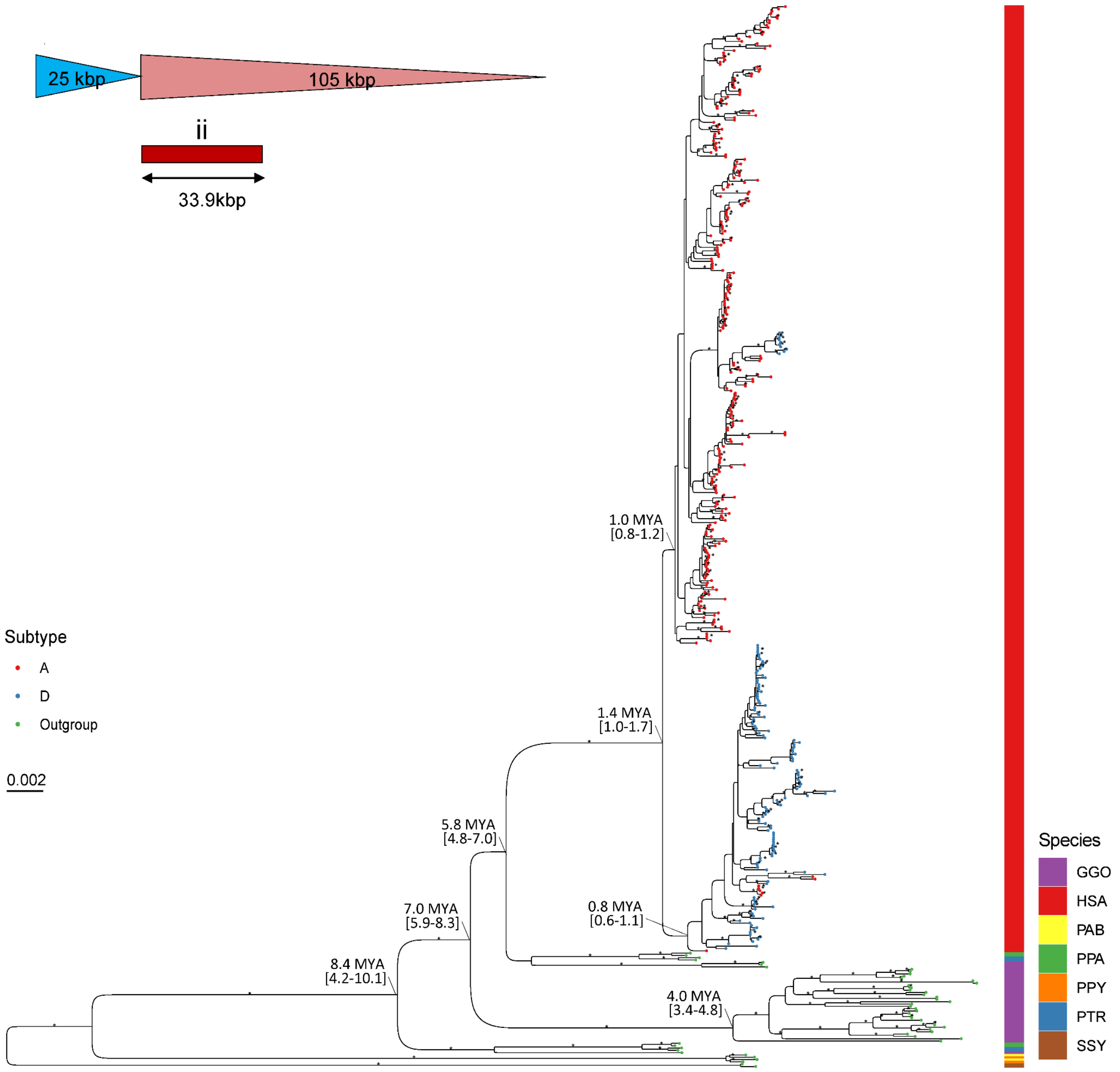


**Supplementary figure 22: Phylogenetic tree of duplicon (ii) of length 33.9 kbp (n=460).** The tree was outgrouped with single-copy macaque ortholog (absent in the tree). The human copies located in LCRA and D are indicated as red and blue, respectively. The corresponding species is indicated on the right annotation track; chimpanzee (PTR), bonobo (PPA), human (HSA), gorilla (GGO), Sumatran orangutan (PAB), Bornean orangutan (PPY) and siamang (SSY). Notable divergence times among human copies are indicated as million years ago (MYA) with the 95% confidence interval. The bootstrap value of >95 is indicated as an asterisk.

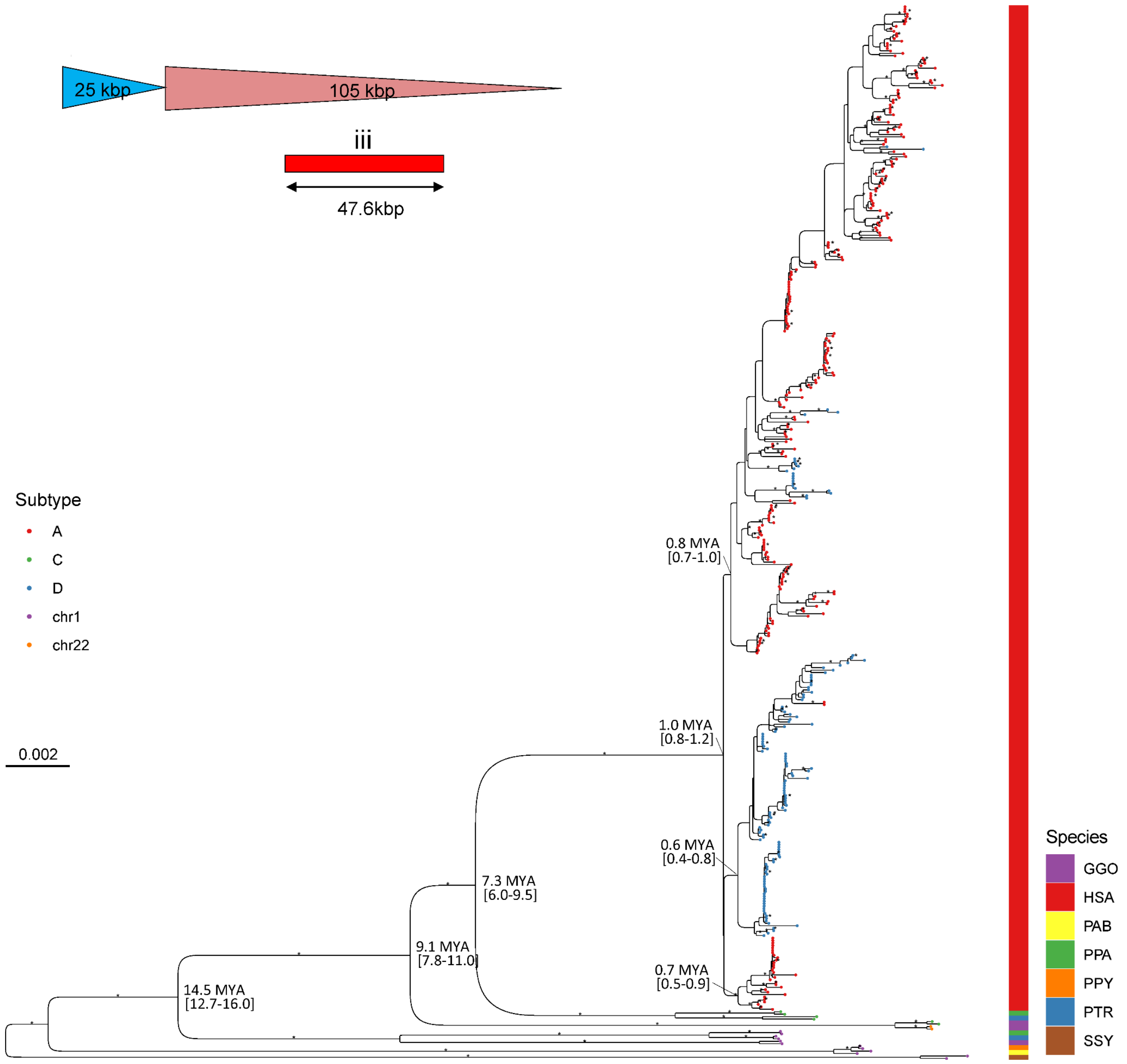


**Supplementary figure 23: Phylogenetic tree of duplicon (iii) of length 47.6 kbp (n=430).** The tree was outgrouped with single-copy macaque ortholog (absent in the tree). The copies located in LCRA, C and D are indicated as red, green and blue, respectively. Additionally, the chromosomal origin (chr1 or 22) of orthologs are indicated as violet and orange, respectively. The corresponding species is indicated on the right annotation track; chimpanzee (PTR), bonobo (PPA), human (HSA), gorilla (GGO), Sumatran orangutan (PAB), Bornean orangutan (PPY) and siamang (SSY). Notable divergence times are indicated as million years ago (MYA) with the 95% confidence interval. The bootstrap value of >95 is indicated as an asterisk.

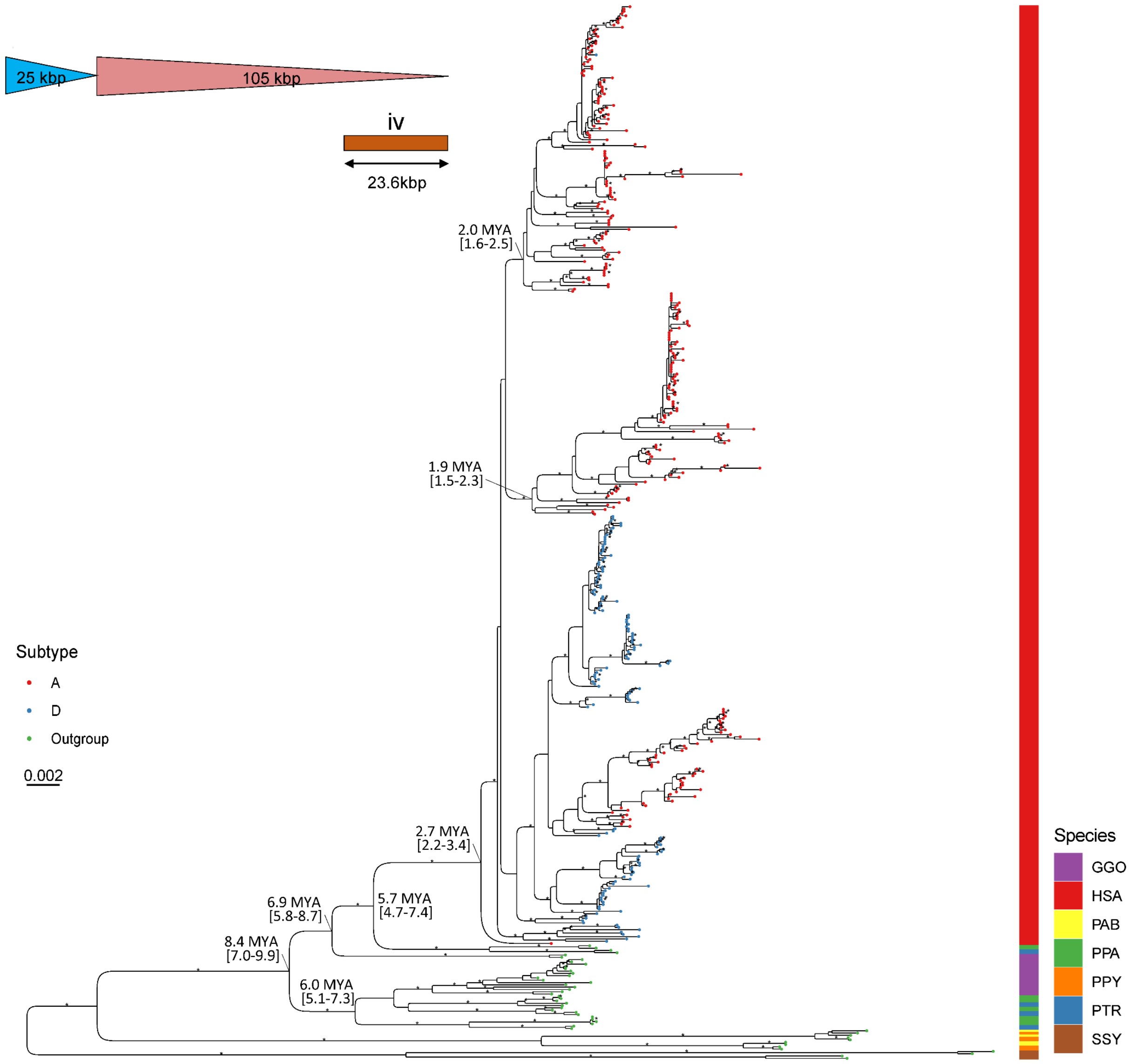


**Supplementary figure 24: Phylogenetic tree of duplicon (iv) of length 23.6 kbp (n=460).** The tree was outgrouped with single-copy macaque ortholog (absent in the tree). The human copies located in LCRA and D are indicated as red and blue, respectively. The corresponding species is indicated on the right annotation track; chimpanzee (PTR), bonobo (PPA), human (HSA), gorilla (GGO), Sumatran orangutan (PAB), Bornean orangutan (PPY) and siamang (SSY). Notable divergence times are indicated as million years ago (MYA) with the 95% confidence interval. The bootstrap value of >95 is indicated as an asterisk.

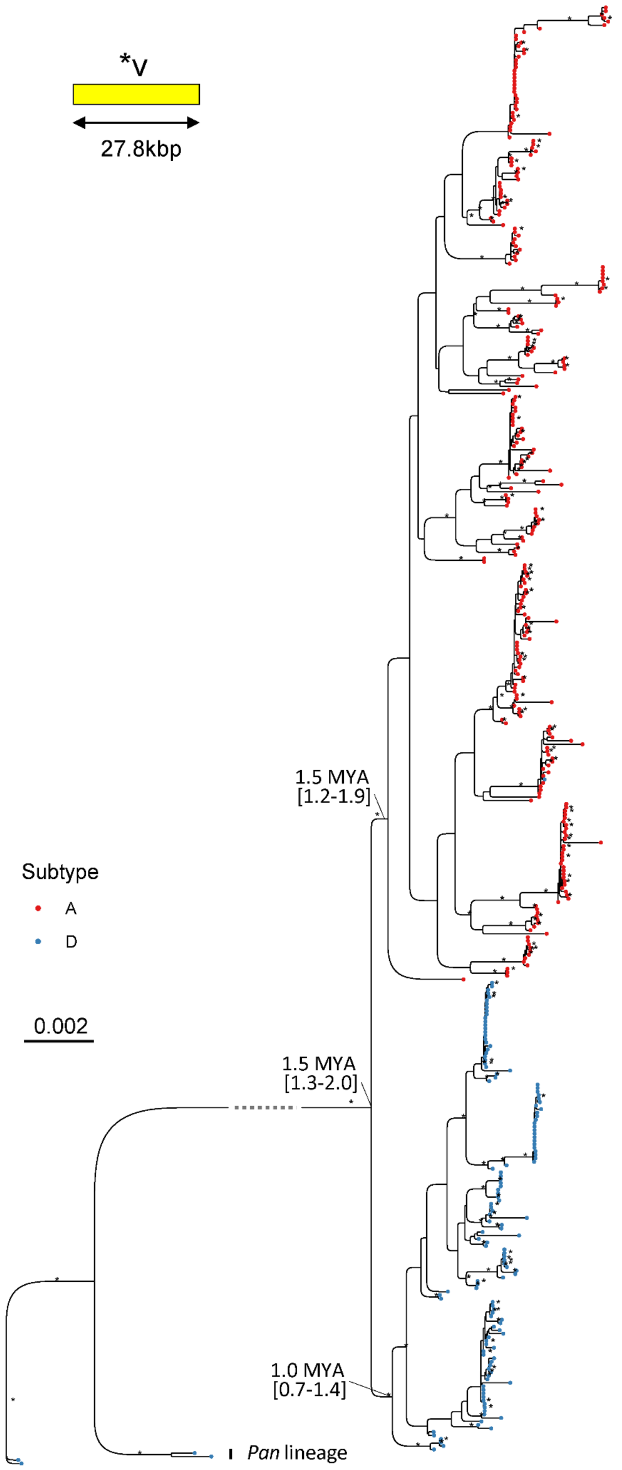


**Supplementary figure 25: Phylogenetic tree of duplicon (v) of length 28 kbp (n=416).** The tree was outgrouped with chimpanzee and bonobo (*Pan* lineage) orthologs. The human copies located in LCRA and D are indicated as red and blue, respectively. Notable divergence times among human copies are indicated as million years ago (MYA) with the 95% confidence interval. The bootstrap value of >95 is indicated as an asterisk.

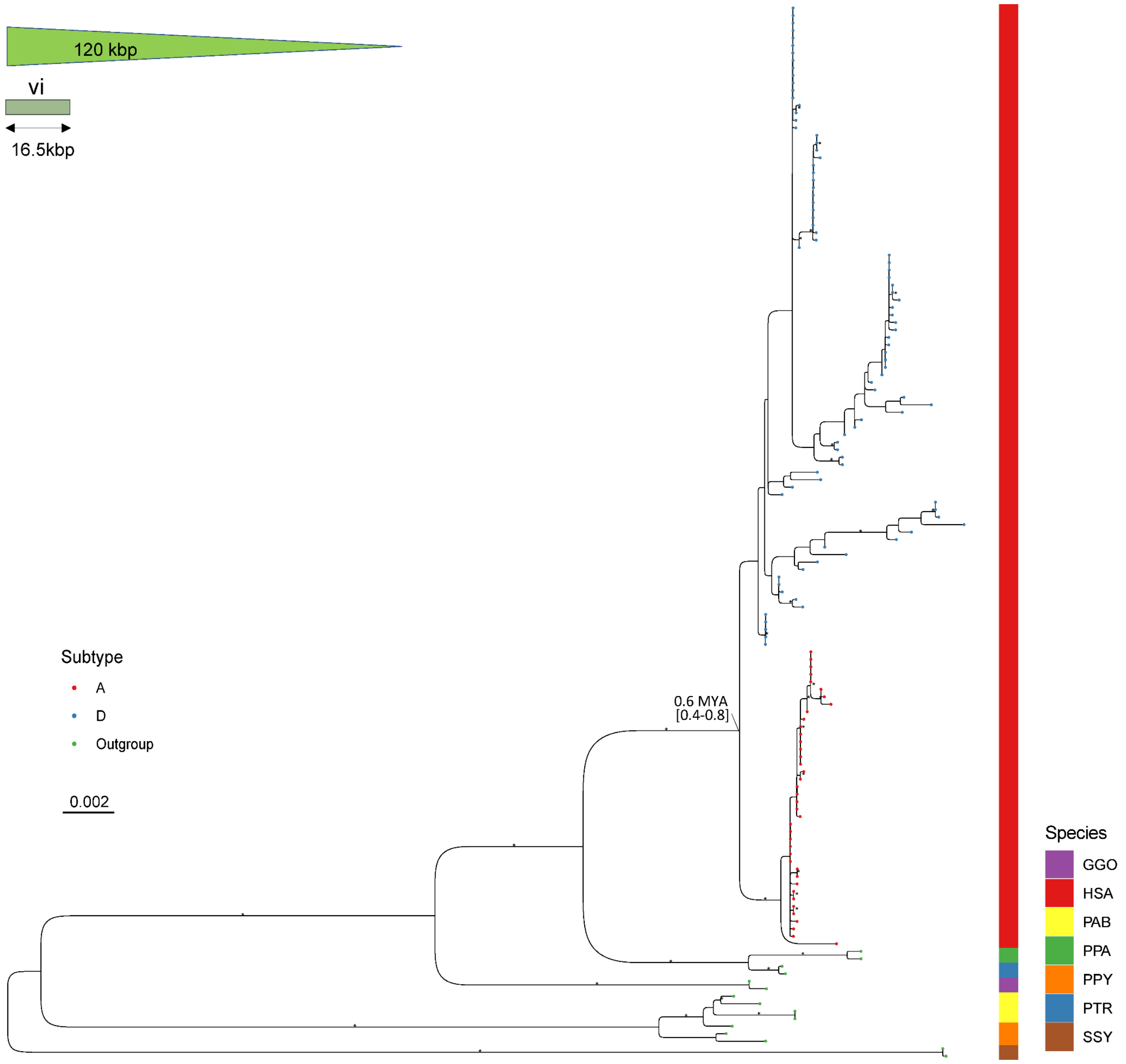


**Supplementary figure 26: Phylogenetic tree of duplicon (vi) of length 16.5 kbp (n=190).** The tree was outgrouped with single-copy macaque ortholog (absent in the tree). The human copies located in LCRA and D are indicated as red and blue, respectively. The corresponding species is indicated on the right annotation track; chimpanzee (PTR), bonobo (PPA), human (HSA), gorilla (GGO), Sumatran orangutan (PAB), Bornean orangutan (PPY) and siamang (SSY). Notable divergence times are indicated as million years ago (MYA) with the 95% confidence interval. The bootstrap value of >95 is indicated as an asterisk.

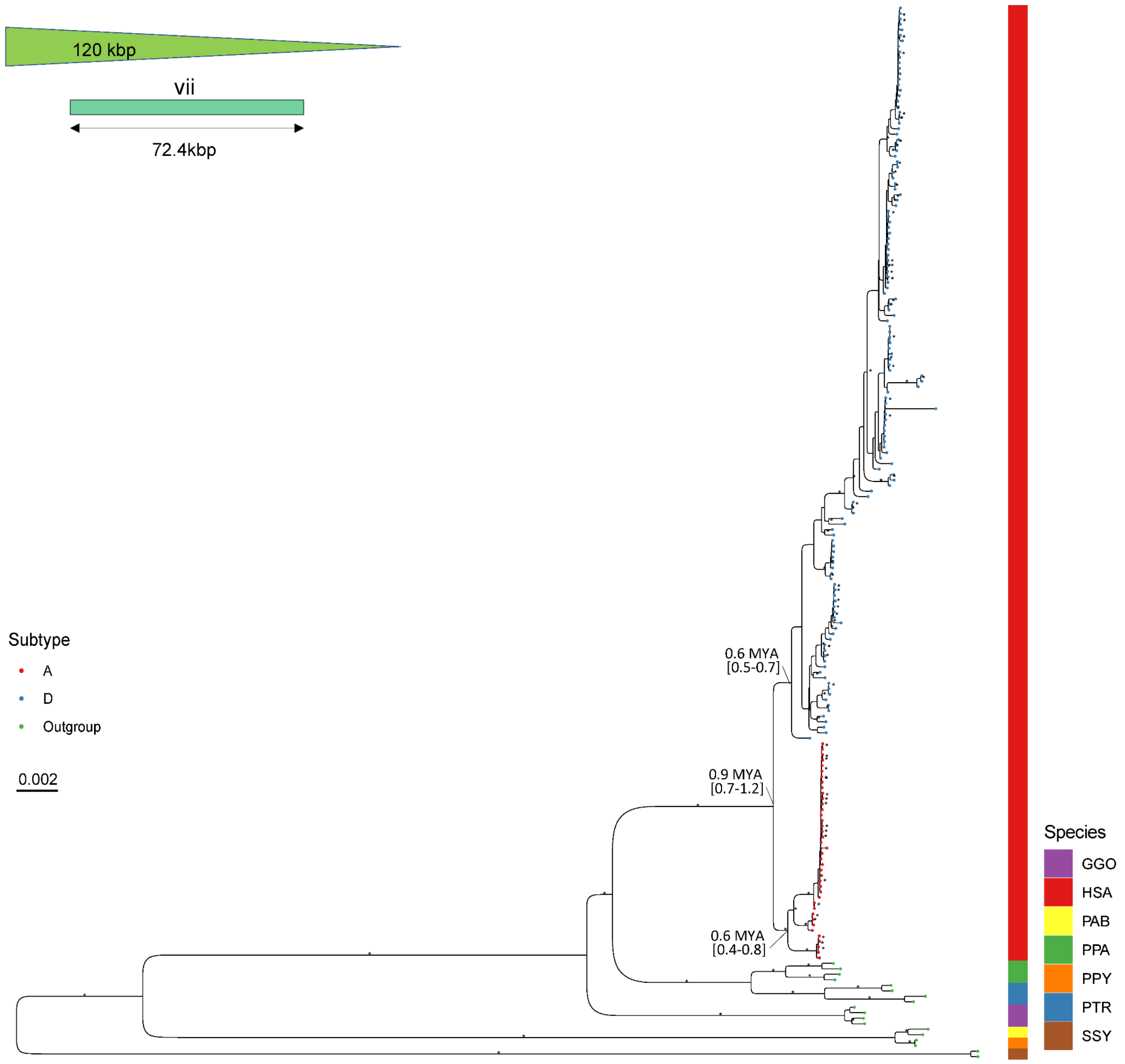


**Supplementary figure 27: Phylogenetic tree of duplicon (vii) of length 72.4 kbp (n=193).** The tree was outgrouped with single-copy macaque ortholog (absent in the tree). The human copies located in LCRA and D are indicated as red and blue, respectively. The corresponding species is indicated on the right annotation track; chimpanzee (PTR), bonobo (PPA), human (HSA), gorilla (GGO), Sumatran orangutan (PAB), Bornean orangutan (PPY) and siamang (SSY). Notable divergence times are indicated as million years ago (MYA) with the 95% confidence interval. The bootstrap value of >95 is indicated as an asterisk.

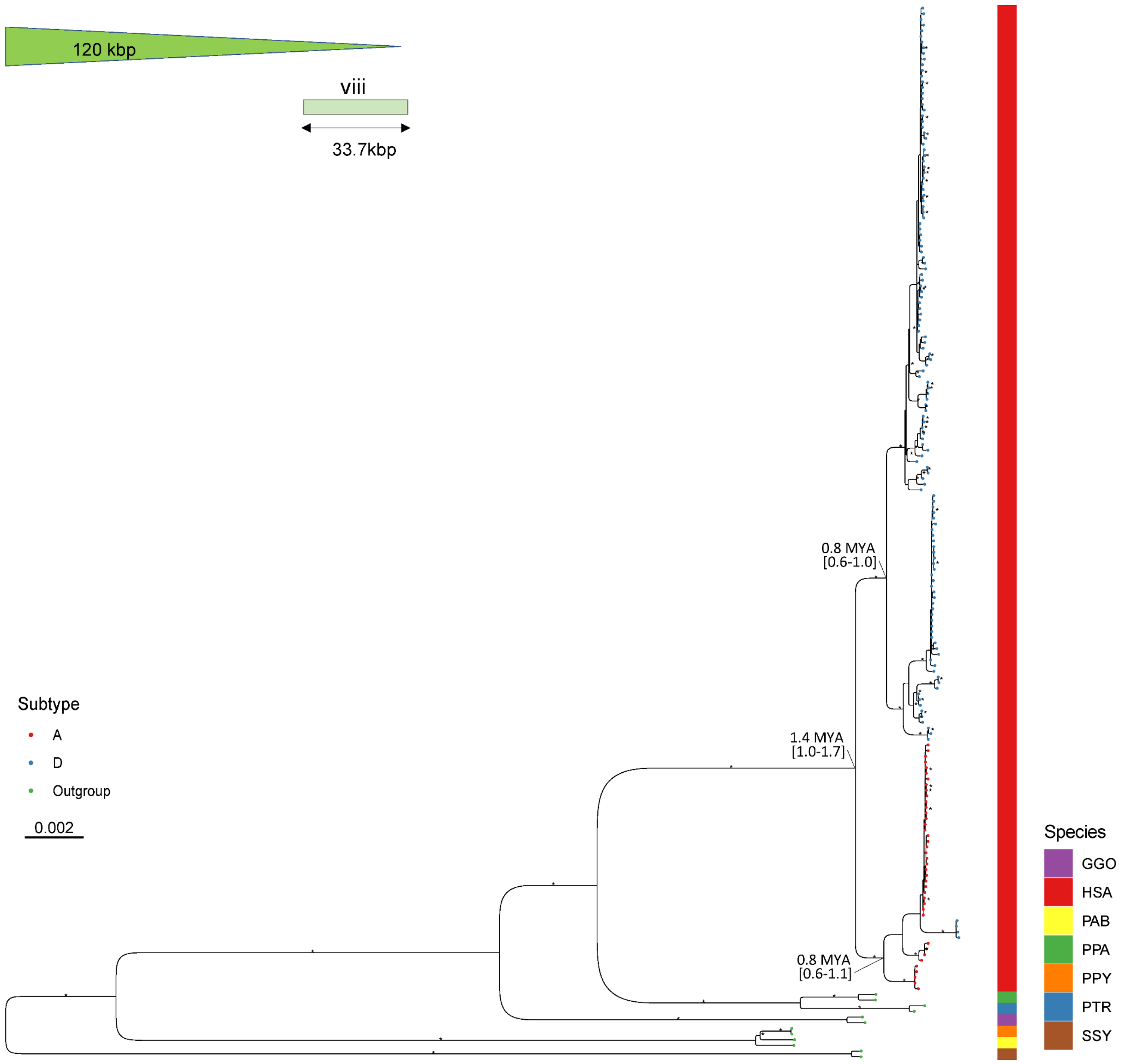


**Supplementary figure 28: Phylogenetic tree of duplicon (viii) of length 33.7 kbp (n=187).** The tree was outgrouped with single-copy macaque ortholog (absent in the tree). The human copies located in LCRA and D are indicated as red and blue, respectively. The corresponding species is indicated on the right annotation track; chimpanzee (PTR), bonobo (PPA), human (HSA), gorilla (GGO), Sumatran orangutan (PAB), Bornean orangutan (PPY) and siamang (SSY). Notable divergence times are indicated as million years ago (MYA) with the 95% confidence interval. The bootstrap value of >95 is indicated as an asterisk.

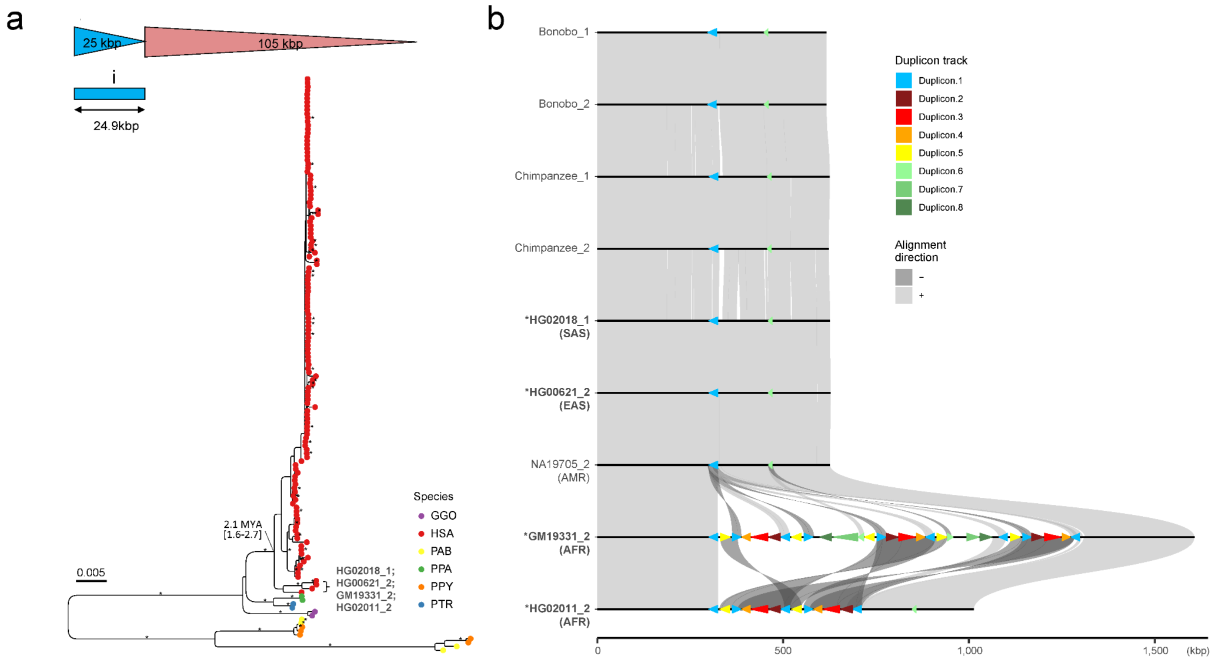


**Supplementary figure 29: Phylogenetic tree of the first proximal copy of duplicon (i) located in LCRA (n=150). a**) The tree was outgrouped with macaque and orangutan orthologs. For the outgroup orthologs, the copies located in LCRA are included. The corresponding species is indicated in different colors; chimpanzee (PTR), bonobo (PPA), gorilla (GGO), Bornean orangutan (PPY), Sumatran orangutan (PAB) and human (HSA). The deepest coalescent time among human copies is indicated as million years ago (MYA) with the 95% confidence interval. The bootstrap value of >95 is indicated as an asterisk. **b**) The first proximal copy of 25 kbp repeat unit located in LCRA among humans indicated by blue arrowhead. Duplicon structures of four putative ancestral haplotypes highlighted in the tree, including two haplotypes, HG02018_1 and HG00621_2, which show similar structure to chimpanzee and bonobo.

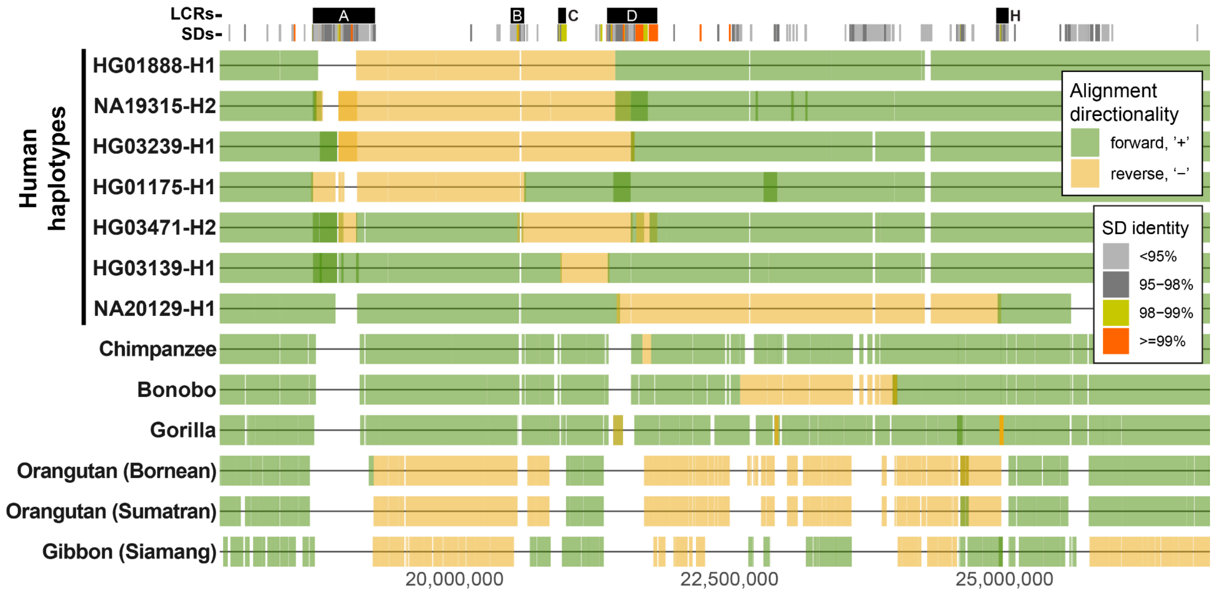


**Supplementary figure 30: Summary of large inversions detected at 22q11.2.**

Alignment directionality of fully assembled sequences across the 22q11.2 region for example human haplotypes and nonhuman primates with respect to T2T-CHM13. Forward oriented alignments are shown in green while reverse oriented alignments in yellow (designating large-scale inversions). On top there is an segmental duplication annotation (SDs) colored by sequence identity followed by annotation by LCRA-D and H as black rectangles.

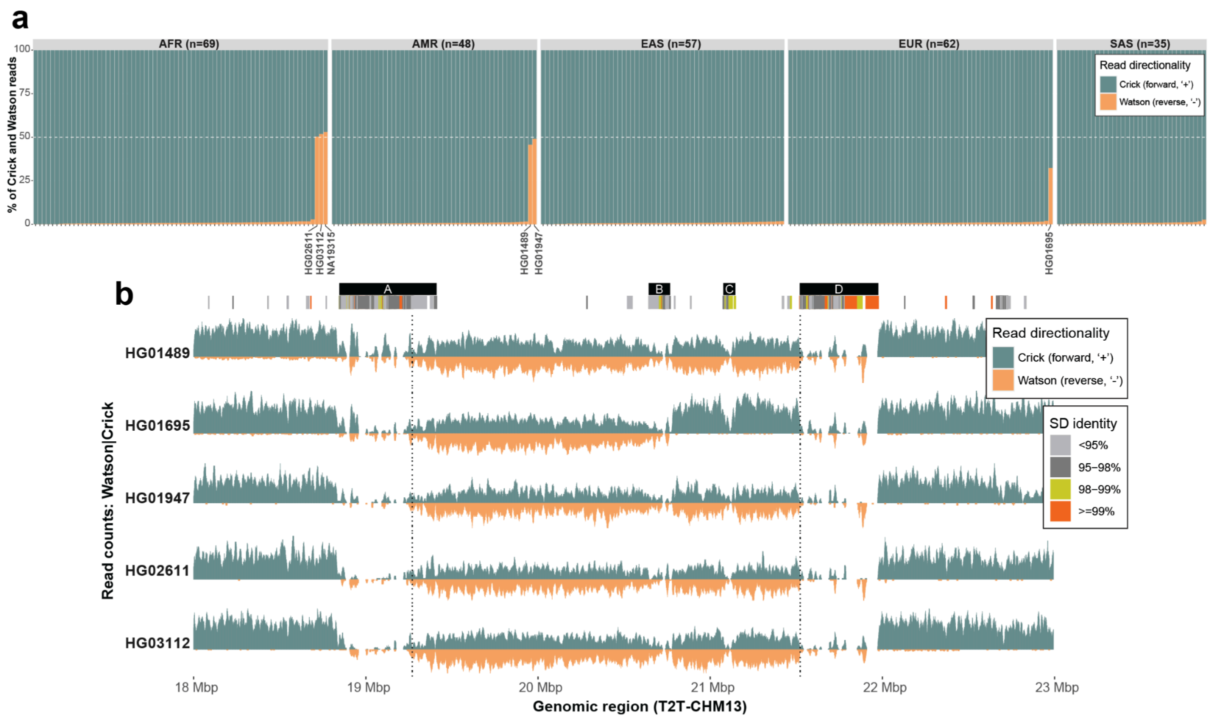


**Supplementary figure 31: Strand-seq-based inversion genotyping.**
**a**) Barplot showing the proportions of forward (Crick - teal) and reverse (Watson - orange) mapping reads to the reference genome for 271 diverse human samples. Samples with about 50% proportion of Watson reads mark heterozygous inversion. Sample (HG01695) with a little lower Watson proportion due to a short A-B inversion. **b**) Read-coverage profiles of Strand-seq data over the 22q11.2 region summarized as binned (bin size: 10 kbp step size: 1 kbp) read counts represented as bars above (teal; Crick read counts) and below (orange; Watson read counts) the midline. Dotted lines highlight the inversion region between LCRA and LCRD. Here, equal coverage of Watson and Crick counts represents a heterozygous inversion as only one homologue is inverted with respect to the reference (T2T-CHM13). Above there is an SD annotation and marked boundaries of SD blocks (A to D) as black rectangles.

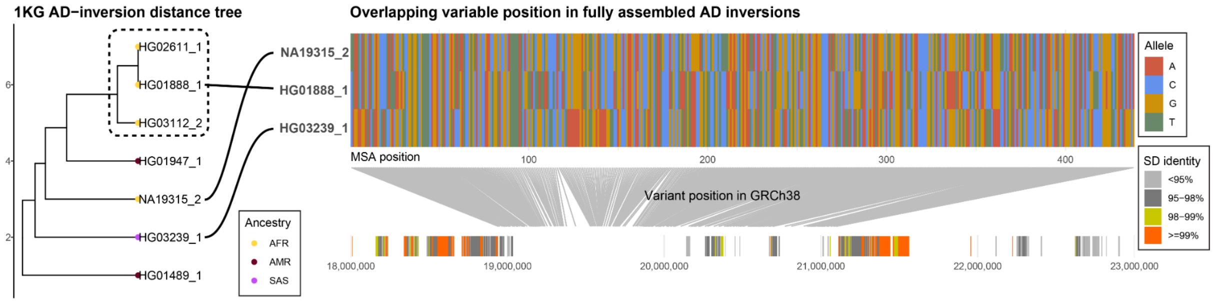


**Supplementary figure 32: Variable positions specific for each unique A-D inversion.**
Left: UPGMA distance tree distance tree constructed from phased SNVs using assemblies and Strand-seq data. Right: Visualization of variable SNV positions among three fully assembled AD inversions projected on GRCh38 assembly.

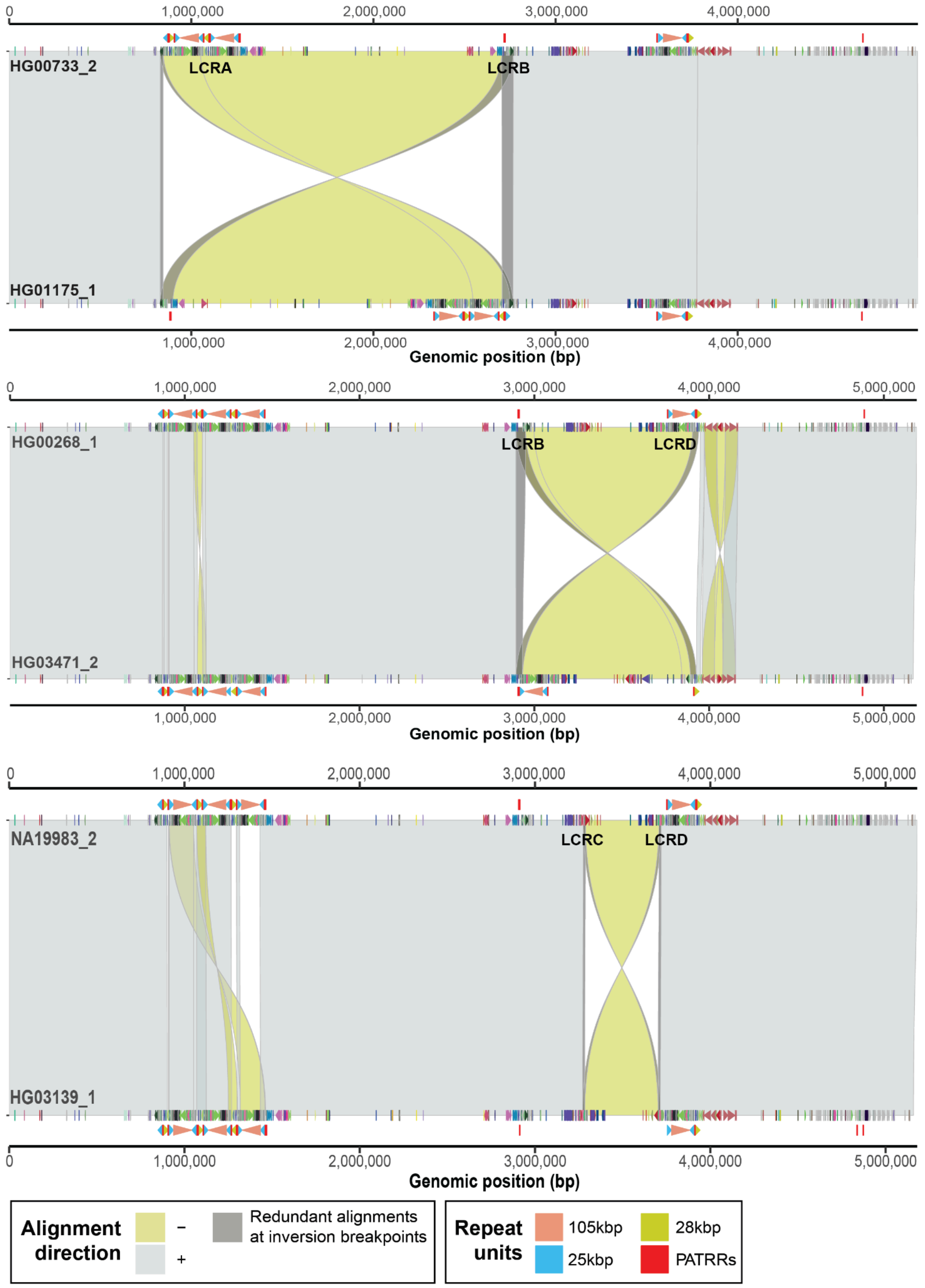


**Supplementary figure 33: Large-scale inversions mediated by various pairs of LCRA-to-D.**
Miropeats-style plot showing the alignments (direct '+' - gray, inverted '-' - yellow) between direct (target at the top) and inverted (query at the bottom) haplotypes for inversions between LCRA-B, LCRB-D, and LCRC-D. There is a duplicon annotation specific to query and target sequences shown as directional arrowheads colored by unique duplicon ID. Positions of 105 (red), 25 (blue), and 28 (yellow) kbp long repeat units are shown below the

query and above the target sequence along with PATRRs shown as red rectangles. Redundant (overlapping) alignments at the inversion boundaries are highlighted in dark gray color.

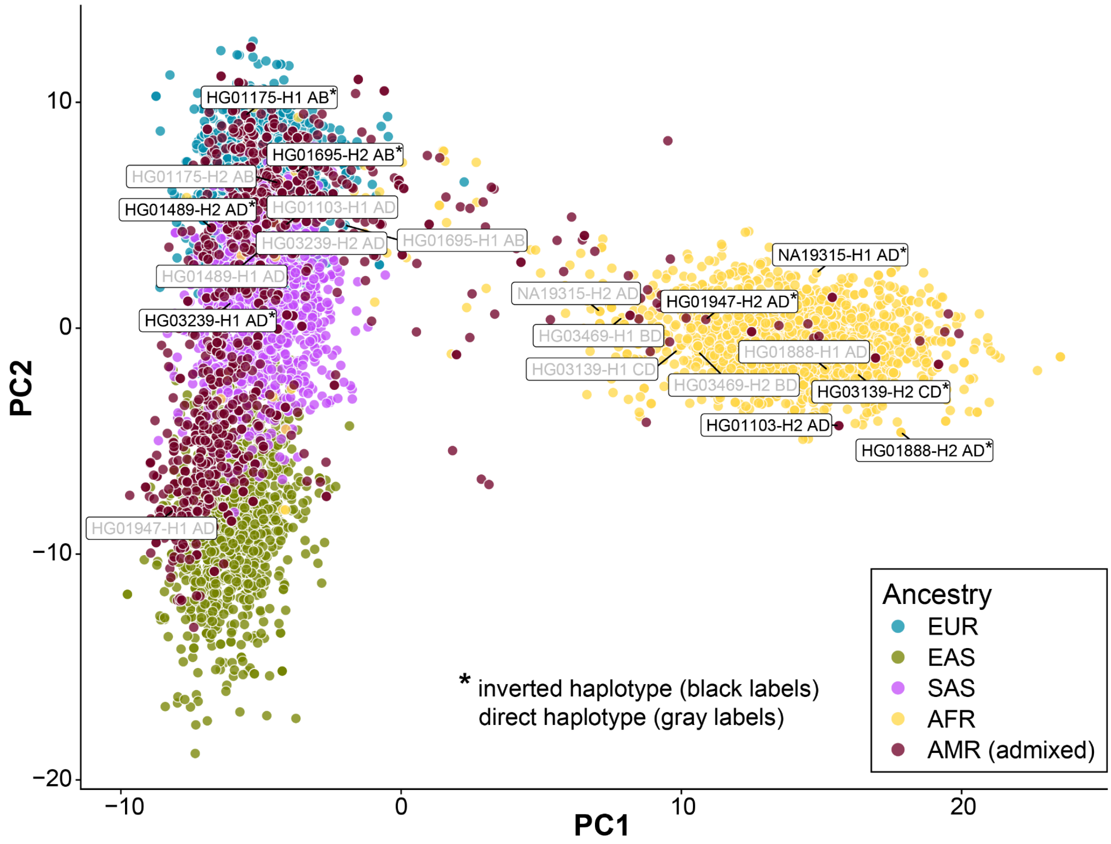


**Supplementary figure 34: Population ancestry of inverted haplotypes.**
Ancestry-specific grouping based on PCA. Inverted haplotypes in the 1KG panel were marked based on the concordance with assembly- or Strand-seq-based phasing. Inverted haplotypes are highlighted using black samples labels with asterisks. Each inversion is marked by LCRs in between which it occurred (AB, BD, CD, and AD).

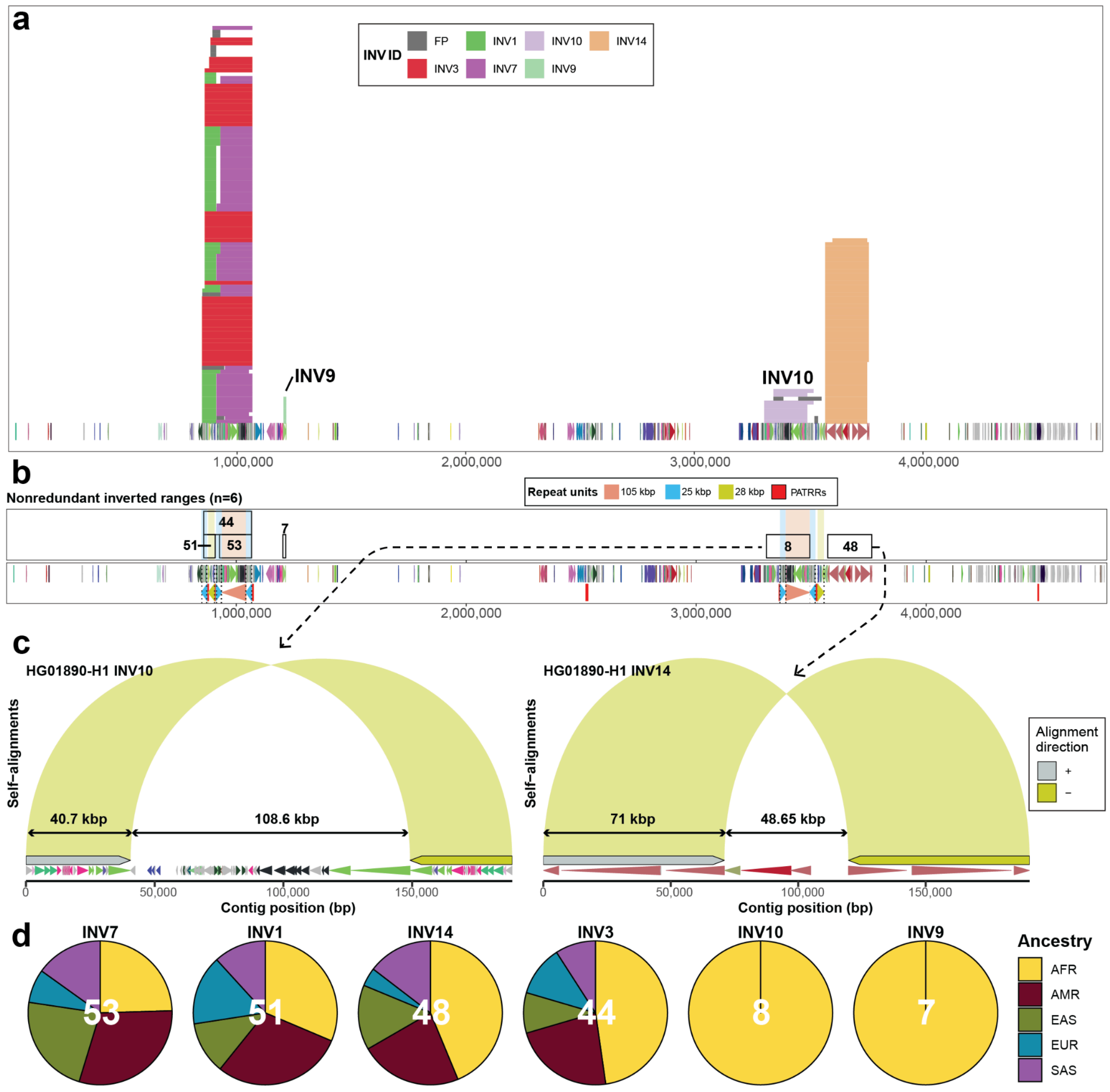


**Supplementary figure 35: Inversion detection within LCRA and D.**

**a**) Visualization of all inverted ranges across all 133 human haplotypes with respect to a single human reference of African ancestry (HG01890-H1) with a single copy of the 105 kbp segment in both LCRA and D. Each inverted alignment defined among all 133 haplotypes is stacked on top of each other and is colored by an alignment group defined by 60% reciprocal overlap (**Methods**). At the bottom there is a duplicon annotation shown as directional arrowheads colored by a unique duplicon ID. **b**) Visualization of nonredundant inversion ranges defined for each alignment group from (a) (**Methods**). Each range is marked with the number that reports the number of inverted alignments in a given group. Below is a duplicon annotation shown as directional arrowheads colored by a unique duplicon ID as well as annotation of 105 (red), 25 (blue) and 28 (yellow) kbp long repeat units shown as directional arrowheads. We also show PATRR positions as red rectangles. **c**) Zoomed-in view of two inversions from LCRD showing the inverted repeats at their flanks (yellow colored arcs). Below them we show duplicon annotation for a given region shown as directional arrowheads colored by a unique duplicon ID. We also highlight the predicted size of the inversion and its flanking repeats on top of two-directional arrows. **d**) Ancestry frequency for each of the six inverted duplications**.**

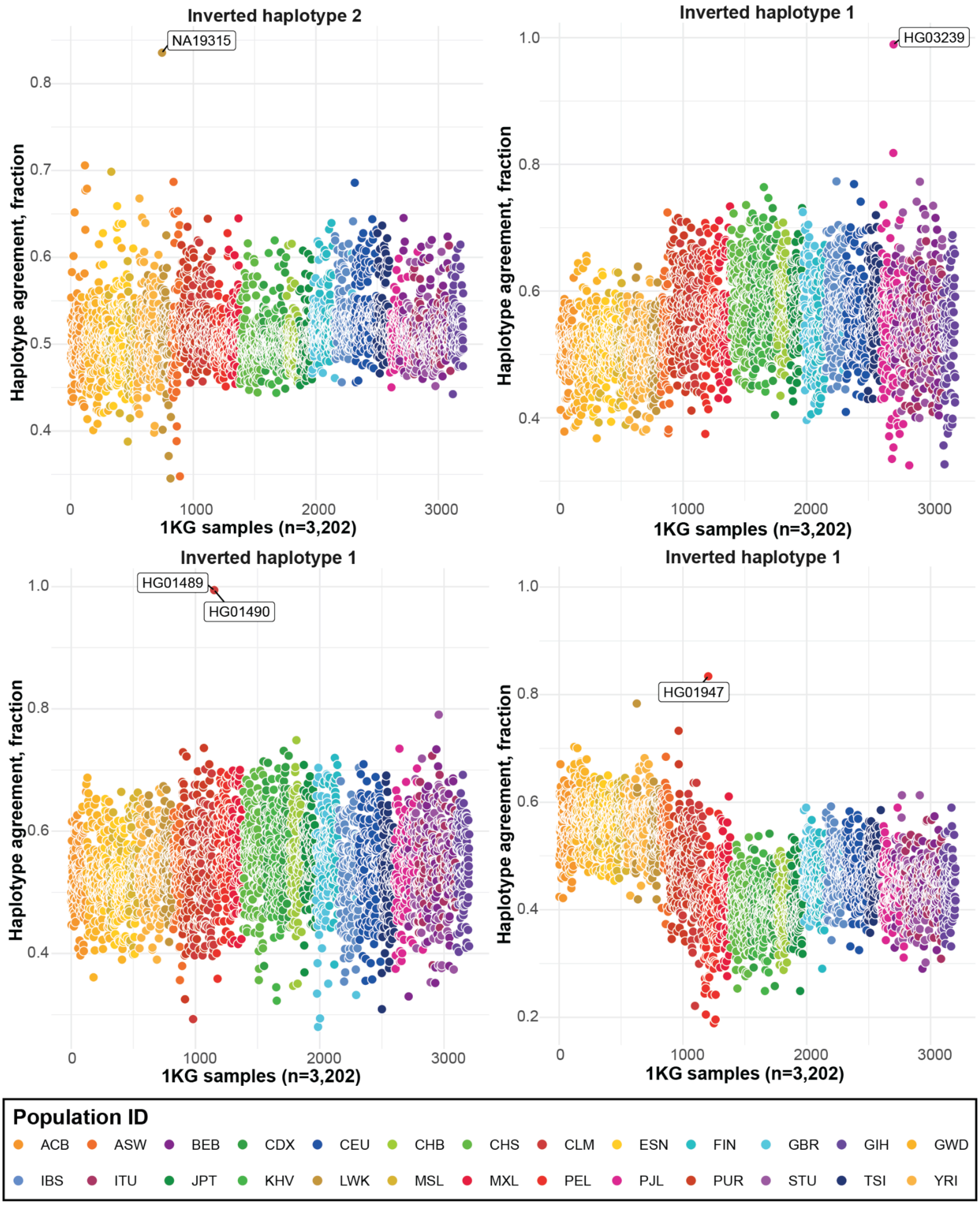


**Supplementary figure 36: LCRA-D inversion genotyping in NA19315, HG03239, HG01489, and HG01947.** Haplotype agreement (y-axis, fraction of matching alleles) between inverted haplotype with respect to 1KG sample panel (n=3,202, GRCh38 coordinates). Each dot represents a single sample colored by a population ID of five major ancestries (African - shades of orange, American - shades of red, East Asian - shades of green, European - shades of blue, and Southeast Asian - shades of purple). Given the observed haplotype agreement between inverted haplotypes and 1KG samples, we do not report any putative inversion carrier among the 1KG panel.

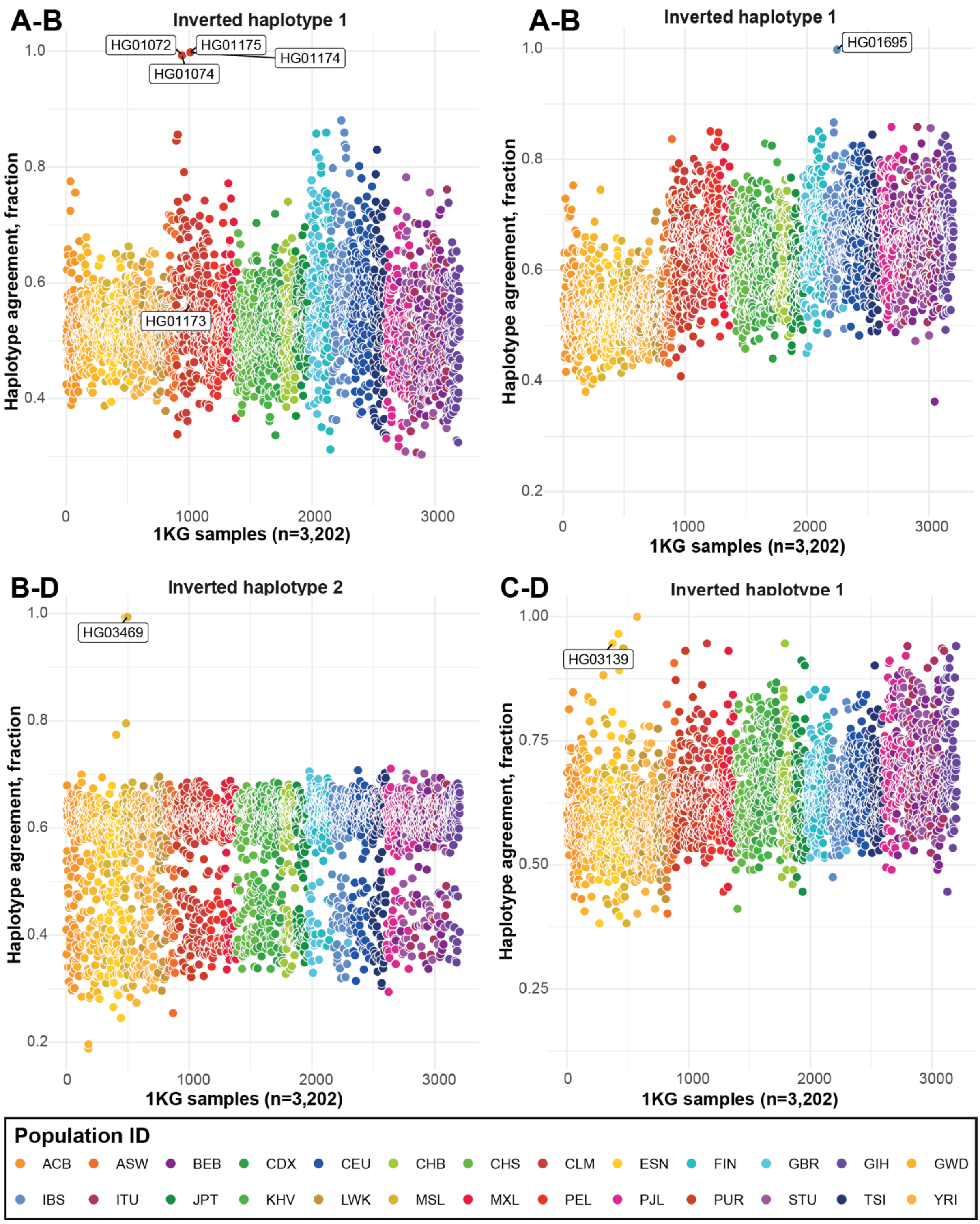


**Supplementary figure 37: LCRA-B, LCRB-D, and LCRC-D inversion genotyping.**
Haplotype agreement (y-axis, fraction of matching alleles) between inverted haplotype with respect to 1KG sample panel (n=3,202, GRCh38 coordinates). Each dot represents a single sample colored by a population ID of five major ancestries (African - shades of orange, American - shades of red, East Asian - shades of green, European - shades of blue, and Southeast Asian - shades of purple).

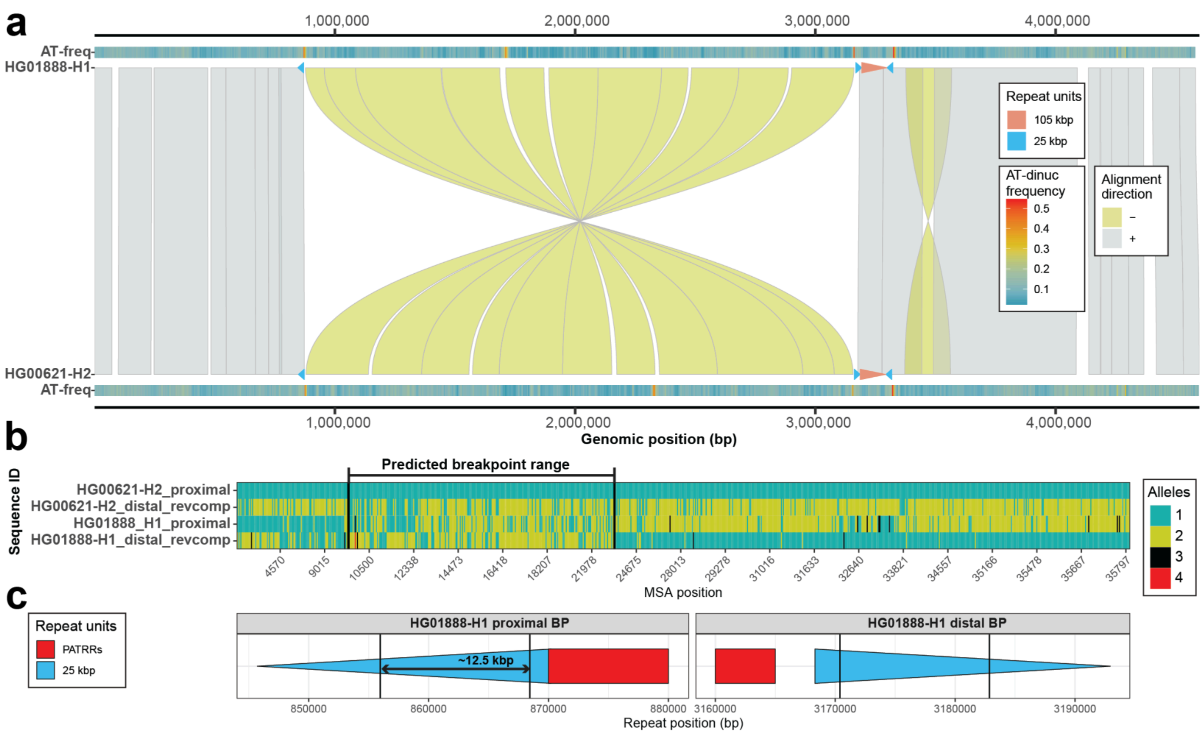


**Supplementary figure 38: Inversion breakpoint mapping in sample HG01888.**

**a**) Miropeats-style plot showing nucmer alignments (direct - gray, '+' and inverted - yellow, '-') between query (bottom) and target (top). Query and target sequences were aligned using (mummer v4.0.0) and its function 'nucmer' with `-mum` parameter. Alignments of 105 (red) and 25 (blue) kbp long repeat units are shown on top of the query and target sequences. Below the query and above target there is a heatmap projecting the frequency of AT-dinucleotides ('AT-freq') in query and target sequences. **b**) MSA visualization between inversions flanking 25 kbp repeat units, including adjacent PATRRs from direct (HG00621-H2) and inverted (HG01888-H1) haplotypes. Paralog-specific variants (PSVs) from the proximal (dark green) and distal (dark yellow) LCRs are colored separately. Gaps in the MSA are colored white and PSVs not present in the proximal and distal LCRs are shown in black and red, respectively. Vertical solid lines depict detected change points where inverted haplotype (HG01888-H1) matches proximal PSVs (dark green) and then switches to match distal PSVs (dark yellow) of the direct haplotype (HG00621-H2) and vice versa. **c**) Position of the narrowed down breakpoint with respect to proximal (LCRA) and distal (LCRD) 25 kbp repeat unit.

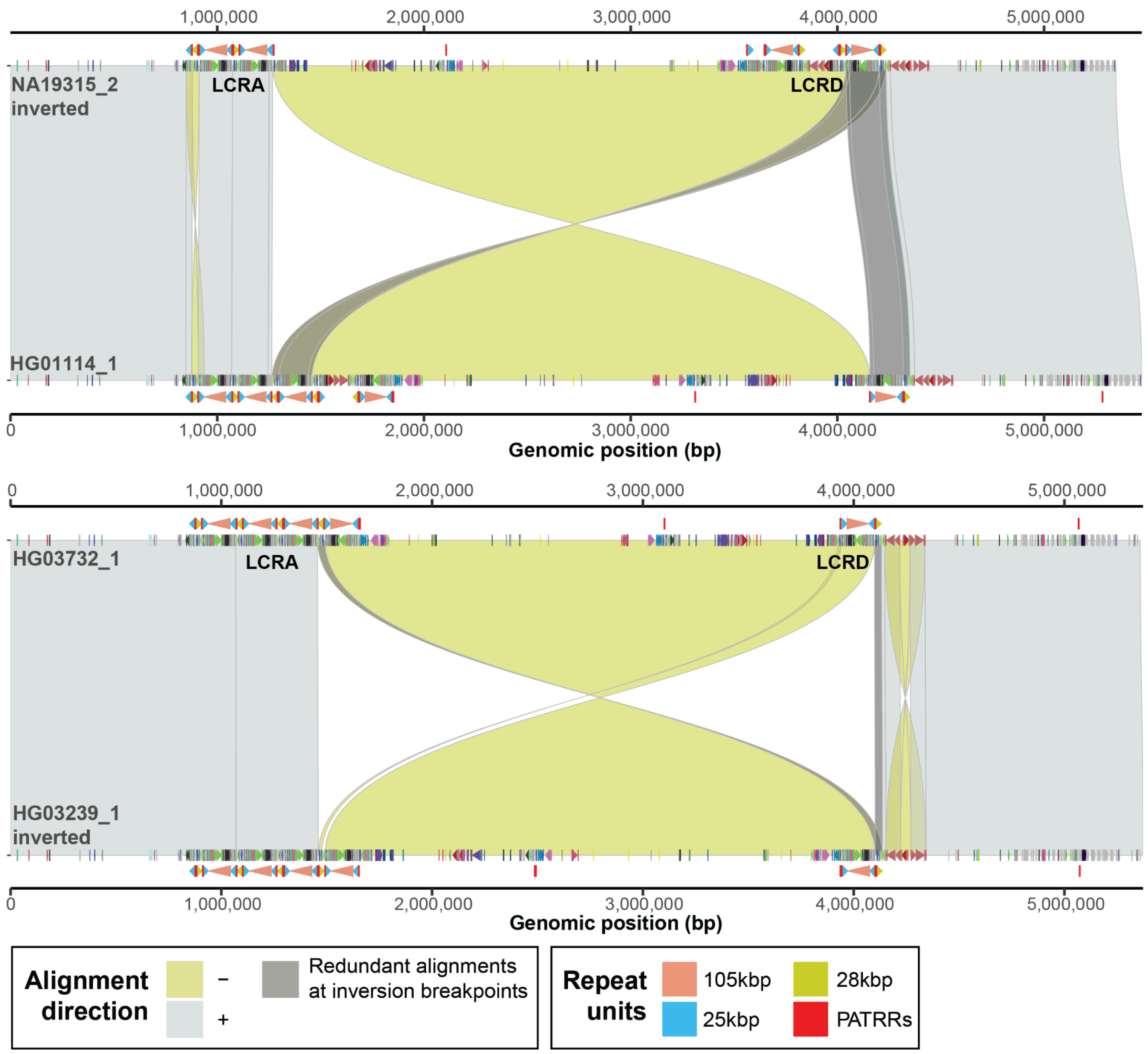


**Supplementary figure 39: A-D inversions in sample NA19315-H2 and HG03732-H1.**
Miropeats-style plot showing the alignments (direct '+' - gray, inverted '-' - yellow) between direct and inverted (marked as so in the plot) haplotypes for A-to-D inversions in sample NA19315-H2 and HG03239-H1. There is a duplicon annotation specific to query and target sequences shown as directional arrowheads colored by unique duplicon ID. Positions of 105 (red), 25 (blue), and 28 (yellow) kbp long repeat units are shown below the query and above the target sequence along with PATRRs shown as red rectangles. Redundant (overlapping) alignments at the inversion boundaries are highlighted in dark gray color.

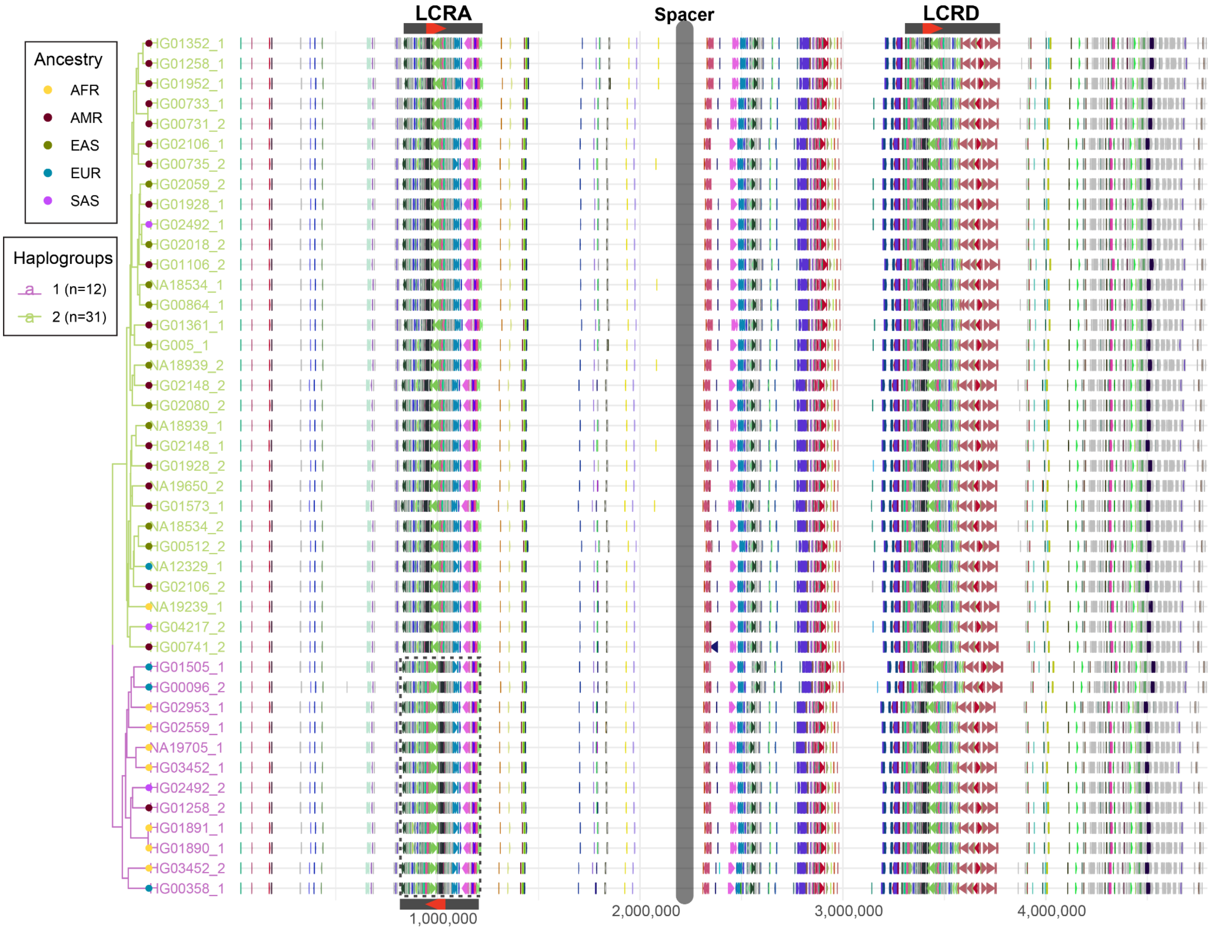


**Supplementary figure 40: Haplotypes with a single copy of 105 kbp segment in LCRA and D (n=43).** **Left**: UPGMA tree constructed based on DupMasker distance between all pairs of haplotypes. The tree is cut into two major haplogroups (‘1’ - purple; ‘2’ - green) and each haplotype is marked by a colored point marking its ancestry. **Right**: Visualization of DupMasker annotation for all 43 haplotypes clustered by the UPGMA tree. We highlight LCRA with a dotted rectangle to show an inversion of the 105 kbp repeat unit in comparison to haplogroup 2.

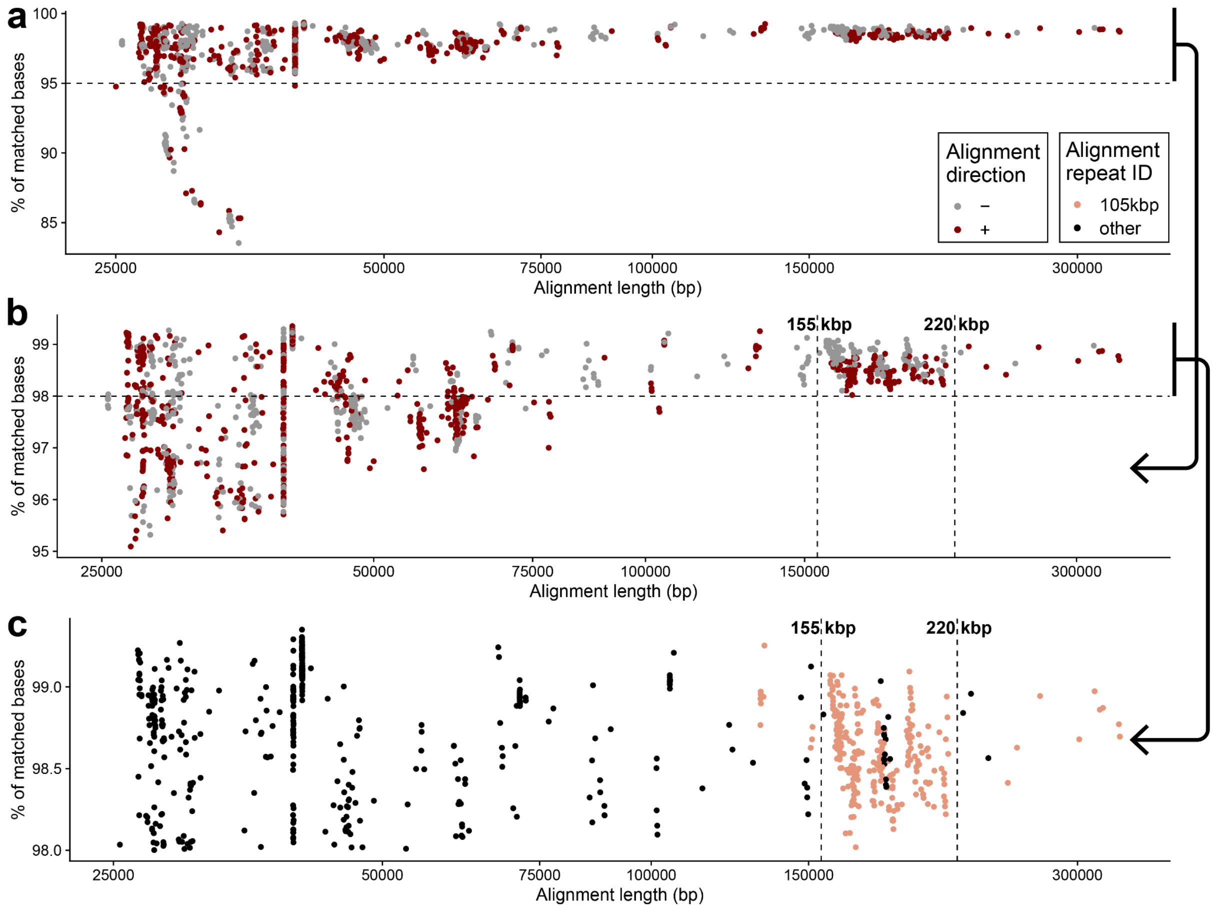


**Supplementary figure 41: Length and sequence identity of self-alignments within each haplotype (n=133).** **a**) Scatterplot showing the percentage of matched bases (y-axis), after removal of large SVs (≥50 bp), with respect to alignment length (x-axis) between LCR and D within each haplotype. Direction of each self-alignment is marked by a color (gray - reverse, red - direct) **b**) Same as (a) but here only self-alignments with ≥95% matching bases are shown. We also highlight a large group of the longest (155-200 kbp) and most identical alignments by vertical dashed lines. **c**) Same as (a) but here only self-alignments with ≥98% matching bases are shown. Self-alignments are colored by the presence (light red) or absence (black) of the 105 kbp repeat unit within each alignment.

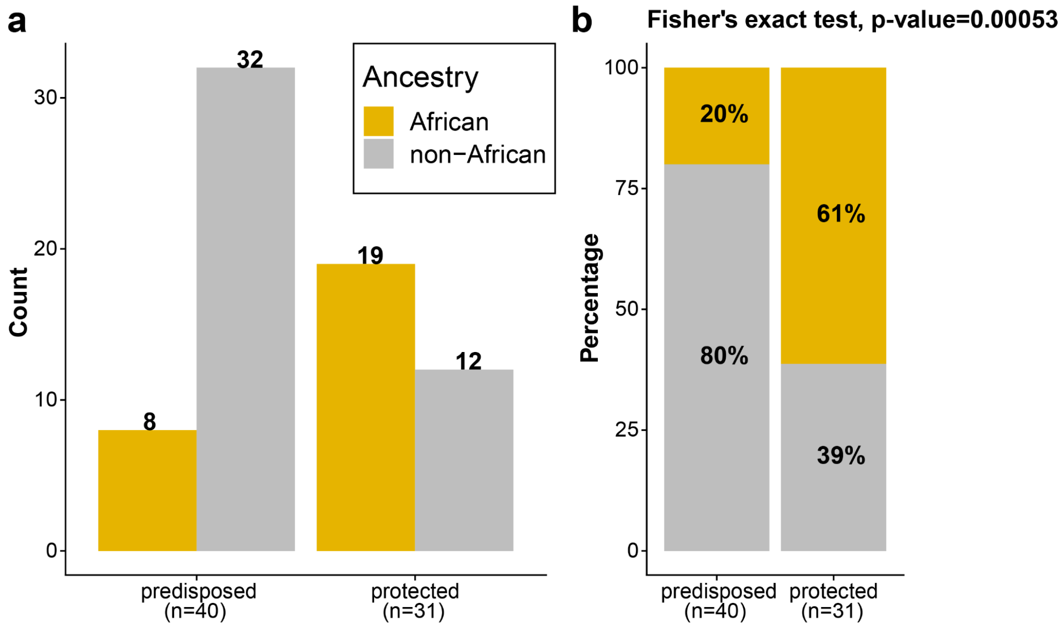


**Supplementary figure 42: Population characteristics of 22q11.2DS predisposed or protected haplotypes.** **a**) Barplot showing the raw counts of protected and predisposed haplotypes among samples of African and non-African ancestry. **b**) Barplot showing the proportions of protected and predisposed haplotypes among samples of African and non-African ancestry. (Fisher’s exact test, two-sided, odds ratio: 0.1627054)

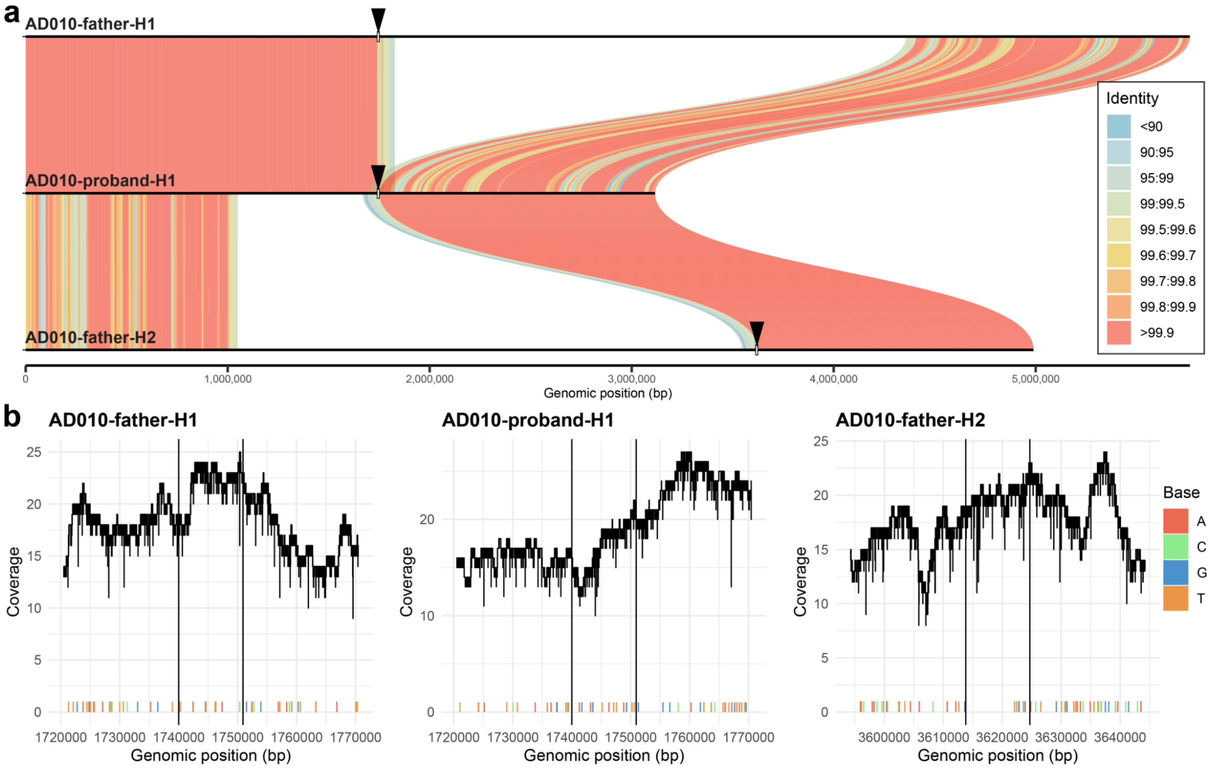


**Supplementary figure 43: Evaluation of haplotype switches in family duo #1.**

**a**) Visualization of binned (bin size: 10 kbp) sequence identity between proband (AD010, proband) haplotype 1 (H1) with respect to both haplotypes (H1 and H2) of a transmitting parent (AD010, father). Proband haplotype is in the middle while parental haplotypes H1 and H2 are at the top and bottom, respectively. Alignments are colored by sequence identity. Haplotype identity transition points are highlighted with black arrowheads.

**b**) Evaluation of sample-specific HiFi reads mapped back to each assembled haplotype from (a). Vertical black solid lines highlight the haplotype transition point from (a). The y-axis shows overall HiFi coverage along with a cumulative coverage of mismatches between HiFi reads and respective haplotype assembly. Low profile of mismatched bases suggests that HiFi reads are in good agreement with the assembly, meaning that no large phasing errors are expected.

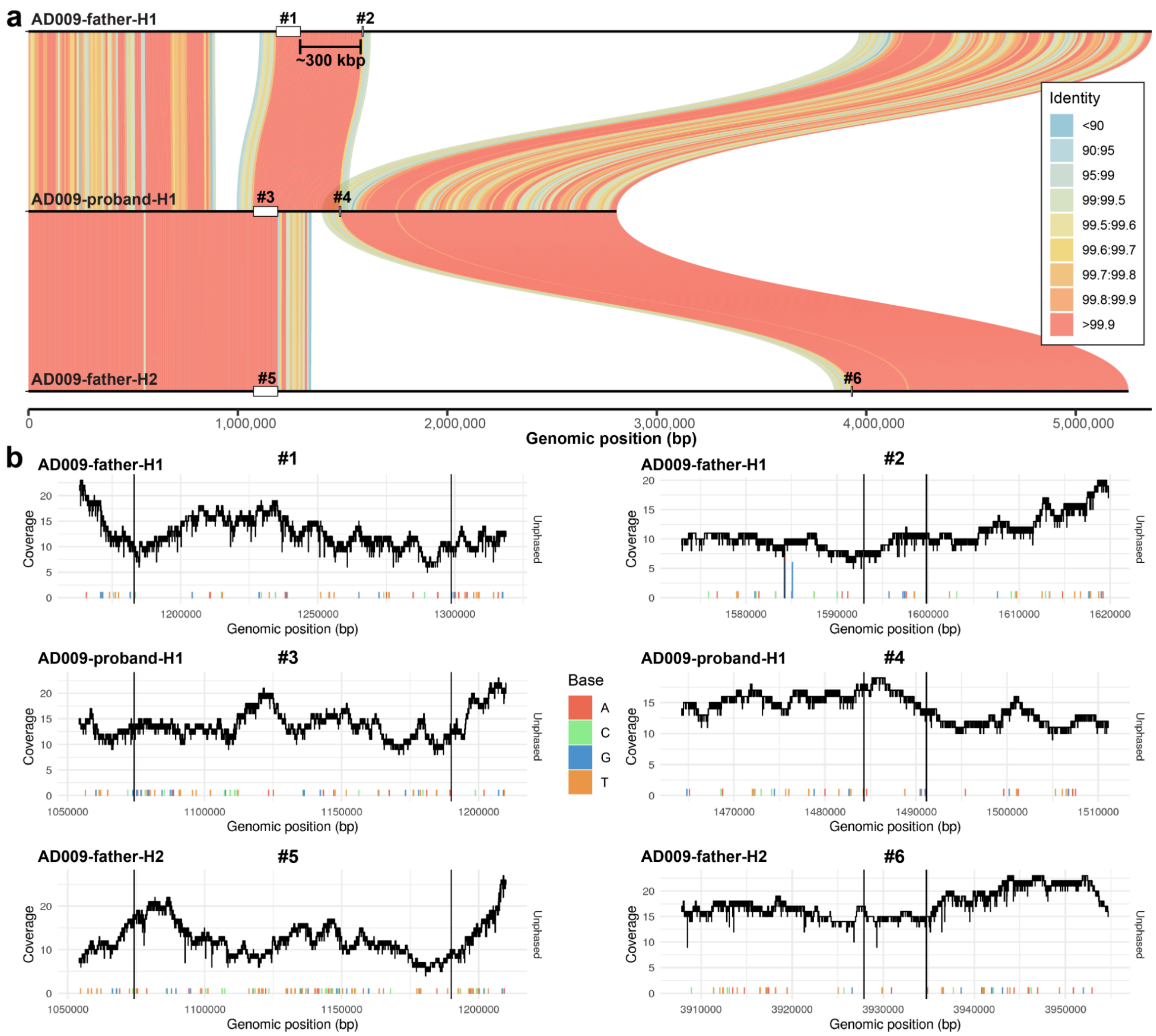


**Supplementary figure 44: Evaluation of haplotype switches in family duo #2.**
**a**) Visualization of binned (bin size: 10 kbp) sequence identity between proband (AD009, proband) haplotype 1 (H1) with respect to both haplotypes (H1 and H2) of a transmitting parent (AD009, father). Proband haplotype is in the middle while parental haplotypes H1 and H2 are at the top and bottom, respectively. Alignments are colored by sequence identity. We highlight the haplotype identity transition points breakpoint IDs #1-6. **b**) Evaluation of sample-specific HiFi reads mapped back to each assembled haplotype from (a). Vertical black solid lines highlight the haplotype transition point from (a). Each breakpoint is labeled by breakpoint ID #1-6. The y-axis shows overall HiFi coverage along with a cumulative coverage of mismatches between HiFi reads and respective haplotype assembly. Low profile of mismatched bases suggest that HiFi reads are in good agreement with the assembly, meaning that no large phasing errors are expected.

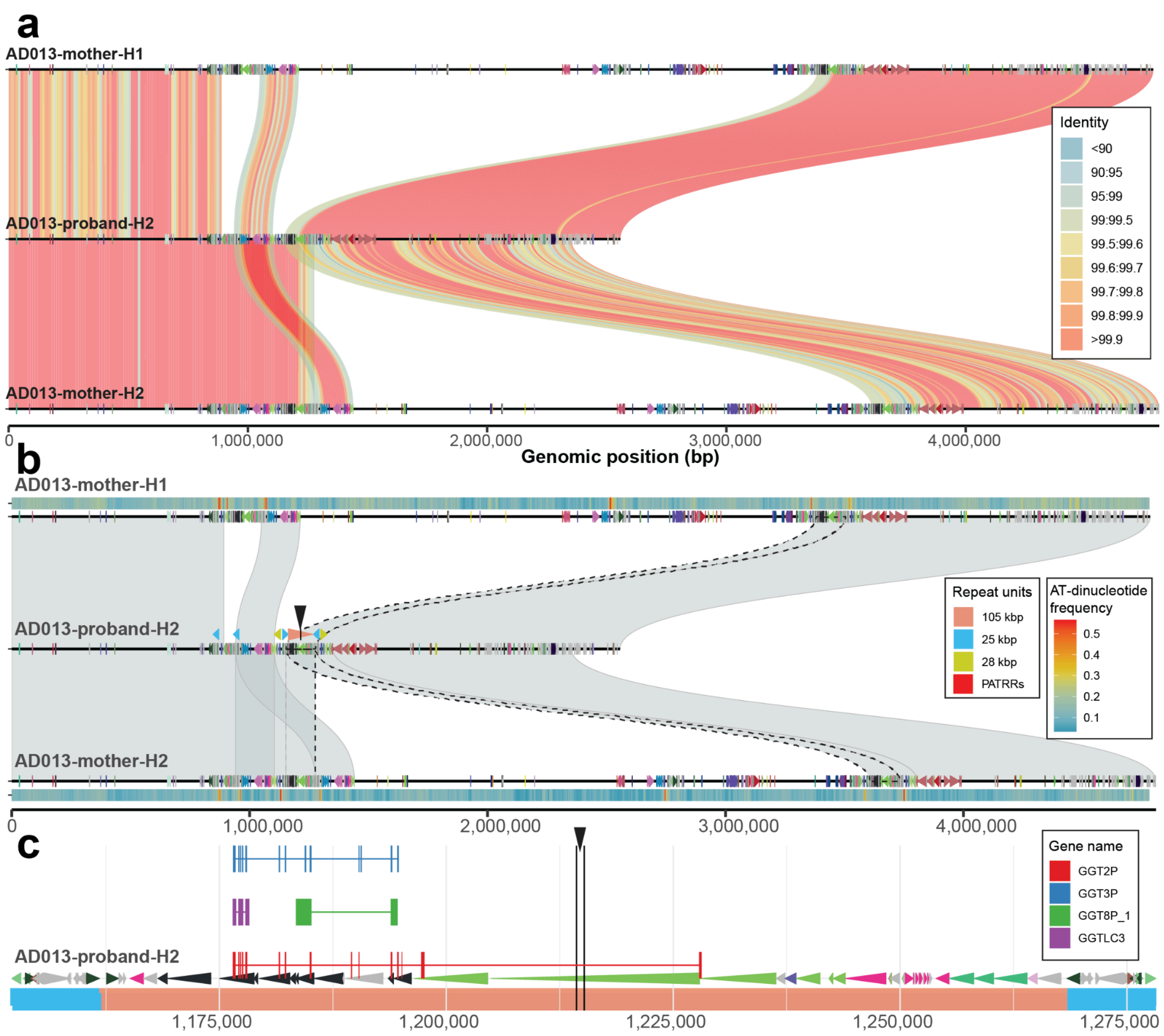


**Supplementary figure 45: Mapping of A-D deletion within a family duo 3.**

**a**) Visualization of binned (bin size: 10 kbp) sequence identity between probands of family 3 with respect to both haplotypes (H1 and H2) of a transmitting parent. Proband haplotype is in the middle while parental haplotypes H1 and H2 are at the top and bottom, respectively. Alignments are colored by sequence identity. On top of each haplotype is a DupMasker annotation depicted as arrowheads colored by unique duplicons. **b**) Similar to a) we show alignments between proband and transmitting parent colored by alignment orientation. Only alignments ≥150 kbp in length are shown. Redundant alignments at the deletion breakpoint are highlighted by black dashed lines. As in a) there is DupMasker annotation on top of each haplotype. For the proband we also show the annotation of common repeat units. Mapped breakpoint within the 105 kbp long repeat is shown by a black arrowhead. On top of parental haplotypes we also show the binned density (bin size: 5 kbp) of PATRRs for a given haplotype. **c**) Zoomed-in view of the duplication structure at the narrowed down deletion breakpoint. Duplicons are shown as colored arrowheads. On top there are mapped exons of the GGT gene family. The predicted breakpoint is marked by vertical black lines with black arrowhead.

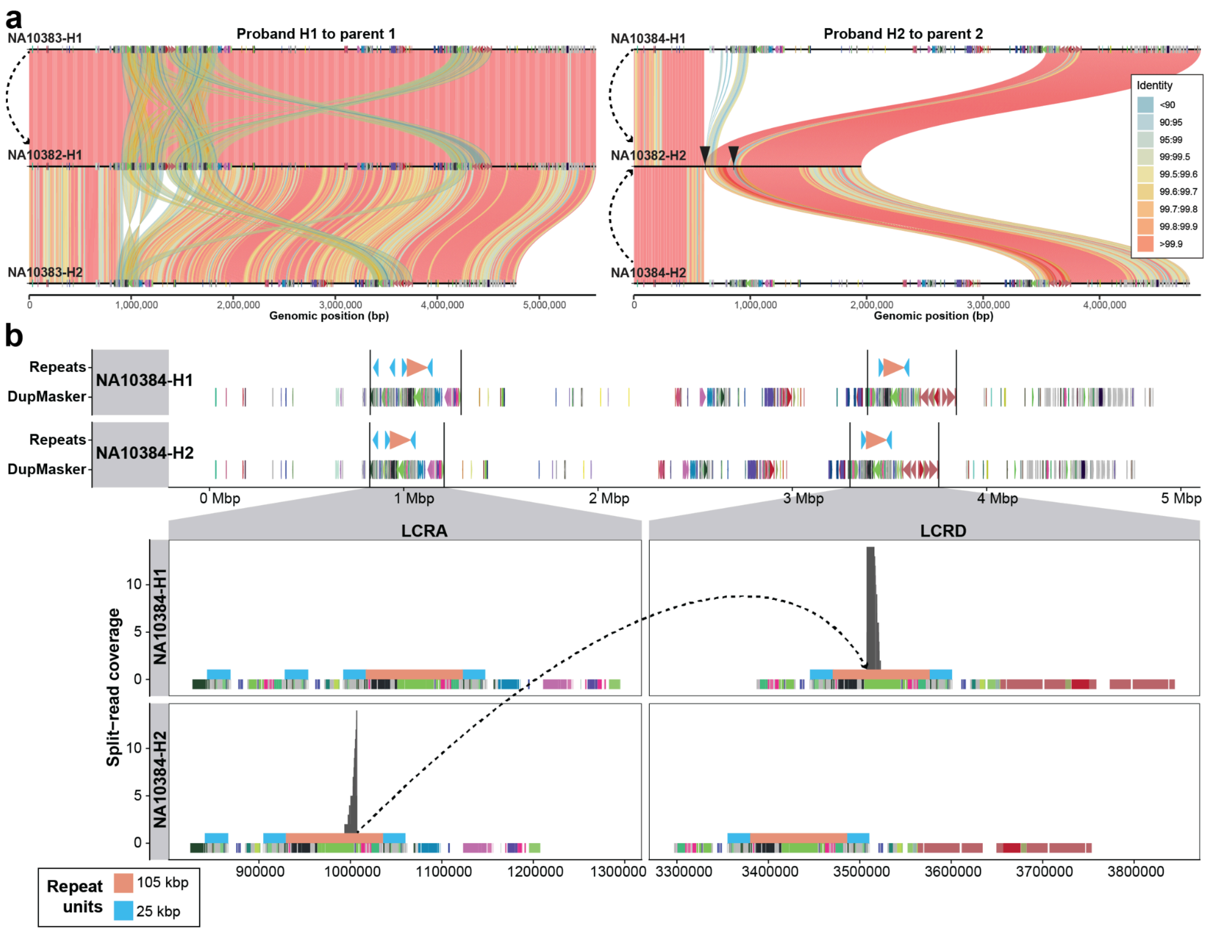


**Supplementary figure 46: Mapping of A-D deletion within a family trio.**

**a**) Visualization of binned (bin size: 10 kbp) sequence identity between proband (NA10382) haplotype 1 (H1) and haplotype 2 (H2) with respect to both haplotypes (H1 and H2) of parent 1 (NA10383, father) and parent 2 (NA10384, mother). Proband haplotype is in the middle while parental haplotypes H1 and H2 are at the top and bottom, respectively. Alignments are colored by sequence identity. Given this, we conclude that proband H1 (NA10382-H1) is inherited from father (NA10383-H1) while the proband haplotype carrying the A-D deletion is inherited from both maternal haplotypes (NA10384 H1 and H2), see dashed arrows. **b**) Visualization of DupMasker structure (colored arrowheads) along with positions of common 105 kbp (red) and 25 kbp (blue) repeat units on top. Boundaries of LCRA and D blocks are highlighted by the vertical solid lines. Below we show a zoomed-in view of the defined LCRA and D regions to show detailed DupMasker annotation along with common repeat positions. We also show coverage HiFi split read mappings as gray rectangles on top of the common repeat annotation. The dashed line shows split read mappings between LCRA on NA10384-H2 and LCRD on the other haplotype (NA10384-H1).

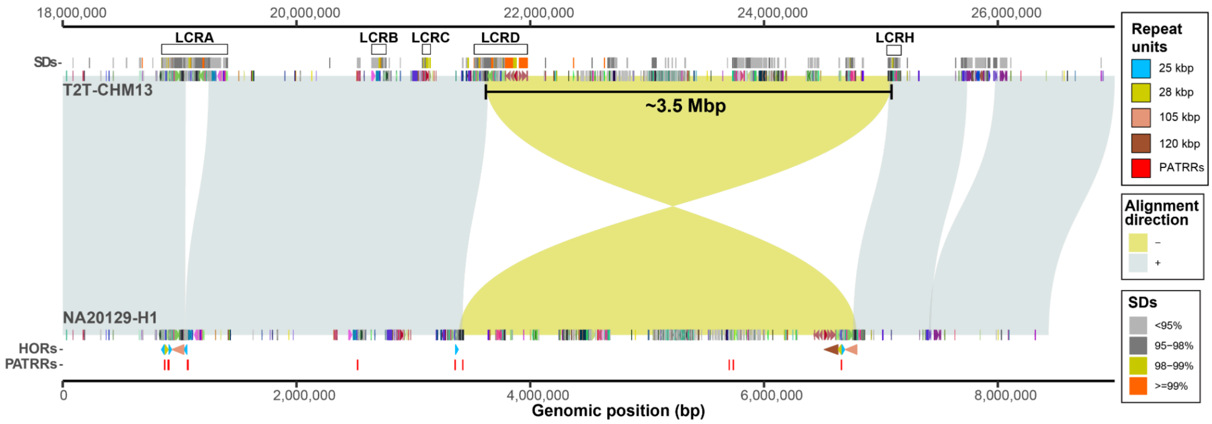


**Supplementary figure 47: Large-scale inversion between LCRD and distal LCR.**
Miropeats-style plot showing the alignments (direct '+' - gray, inverted '-' - yellow) between direct (T2T-CHM13 reference) and inverted haplotypes (NA20129-H1) for large 3.5 Mbp inversion that extends from LCRD to a downstream located breakpoint. There is a duplicon annotation specific to query and target sequences shown as directional arrowheads colored by unique duplicon ID. Positions of 105 (red), 25 (blue), and 28 (yellow) kbp long repeat units are shown below the query sequence along with PATRRs shown as red rectangles. SD annotation of the target sequence is shown on top colored by sequence identity. Lastly, on top we show positions of LCRA-to-D and LCRH regions on the target sequence.

### Supplementary Notes

**Gene annotation at 22q11.2.** There are a number of protein-coding genes and transcribed pseudogenes overlapping our defined repeat units (25 and 105 kbp). This includes protein-coding genes (*GGTLC3*, *USP18*) and transcribed pseudogenes (*GGT2P*, *GGT3P*, *GGT8P_1*, *POM121L15P*, and *POM121L8P*) overlapping the 105 kbp long repeat unit while the 25 kbp repeat unit is occupied by long noncoding RNA (lncRNA) represented by *FAM230* genes. There are also two newly characterized genes (LOC*) at 22q11.2 in the T2T-CHM13 reference (**Supplementary Fig. 14**).

We performed *de novo* gene annotation and expression analysis on selected human haplotypes of the 22q11.2 region using a database of 1.4 billion Iso-Seq reads (Dishuck et al. 2025). We compared alignments of Iso-Seq reads from *FAM230*, *POM121*, and *GGT* gene family members within the 22q11.2 region to the T2T-CHM13 reference versus the rest of the assemblies (**Supplementary Fig. 48**). Highly divergent Iso-Seq reads generally correspond to structural variants (SVs), with transcription extending across SV breakpoints. For example, the >10% divergent *GGTLC3* and *GGT3P* reads correspond to *PI4KAP2-GGTLC3* read-through fusion transcripts. We did not observe any novel protein-coding gene models for these families but did observe usage of an isoform of *CLTCL1* corresponding to the *Pan paniscus* gene model XP_063459430.1, which had not been documented in humans and encodes an additional 44 amino acids relative to the canonical human gene model.

*FAM230* lncRNA paralogs co-localize with the 25 kbp long repeat unit and adjacent palindrome AT-rich regions (PATRRs) (**Supplementary Fig. 14**) are associated with structural variation, including the most common non-Robertsonian translocation, constitutional t(11;22) (Vervoort et al. 2024; Tong et al. 2010). We observe preferential expression of *FAM230B*, *E*, *F*, *G*, *H*, and *J* in testis and sperm (**Supplementary Fig. 49**), raising the possibility that paralog transcription may modulate SV frequency as has been documented in yeast (Xie et al. 2024), given the bias towards parental origin for this translocation (Ohye et al. 2010).

With Iso-Seq data, we find embryonic stem cell and fetal striatum expression of *USP41* in LCR22B (**Supplementary Fig. 50**, **Supplementary Fig. 51**), and despite its annotation as a pseudogene (Rubino et al. 2020), it encodes a 295 amino acid open reading frame. Previous annotations may have been misled by extension into SD sequence for *FAM230G*, preventing unique short-read alignment. Our Iso-Seq data indicate that *USP41* has an additional 70 amino acids in three coding exons on its 3' end, relative to previous lncRNA annotations and its ancestral paralog, *USP18* (**Supplementary Fig. 52**). Recent work provides proteomic support for *USP41* translation and suggests a function distinct from *USP18* (Bonacci et al. 2024), which may increase breast cancer cell migration (Yoon et al. 2023). Investigating the phylogeny of this gene, we predict that *USP18* and *USP41* were likely to be present in African great ape ancestors, although the chimpanzee *USP41* was lost (**Supplementary Fig. 51b-c**). Constructing the phylogenetic tree, we observed that the expansion of USP genes indeed predates speciation of chimpanzees and gorillas (**Supplementary Fig. 53**). However, we find that the divergence time between *USP18* and *USP41* is 6.68 [5.5-7.81] MYA, which is earlier than the reported speciation time of gorilla (8.6 [8.3-9.3] MYA (Kumar et al. 2022)). Predictably, the divergence times of the USP gene tree are underestimated, considering relatively short divergence times to gorilla (5.99 MYA) and chimpanzee (4.64 MYA) orthologs, which is approximately one-third shorter than the expectation. Performing a test of positive selection, we find that the rate of nonsynonymous mutation is greater than the rate of synonymous mutation at *USP18*, *USP41*, and the African great ape ancestor branches; however, we also note that the test was statistically insignificant (**Supplementary Fig. 54**).

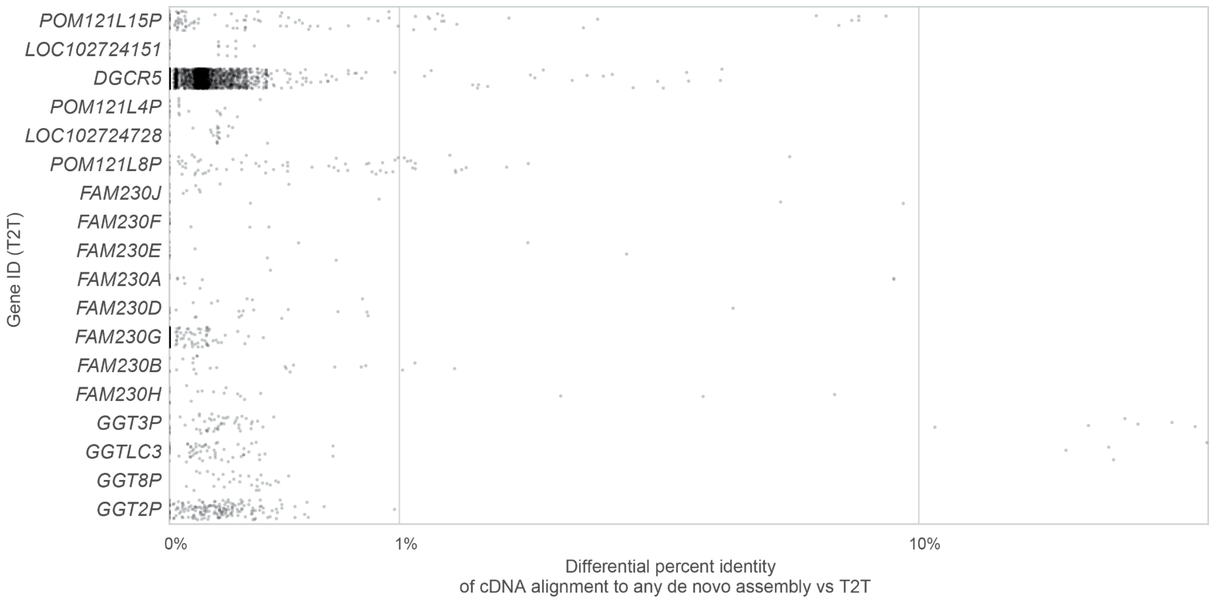


**Supplementary figure 48: Improved alignment of cDNA reads to population assemblies.**
The percent identity differential of the mapping of full-length Iso-Seq transcripts from chr22q11.2 *POM121*, *FAM230*, and *GGT* paralogs to T2T-CHM13 compared to population assemblies. Each point represents an individual Iso-Seq read, and the x-axis indicates the improvement in alignment identity using population assemblies instead of the T2T-CHM13 reference.

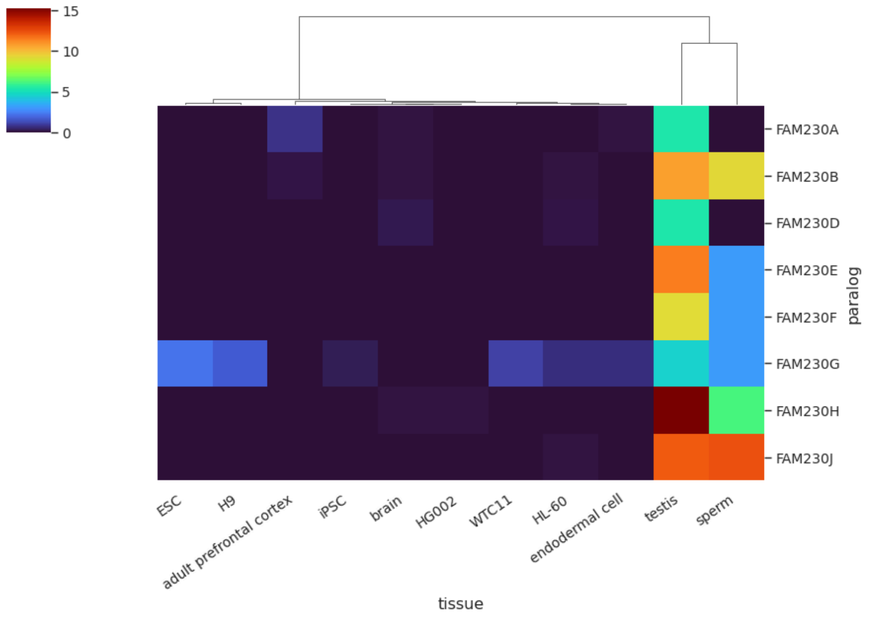


**Supplementary figure 49: Tissue specificity of Iso-Seq expression for *FAM230* paralogs.** Estimated Iso-Seq expression of *FAM230* paralogs as defined in T2T-CHM13, in reads per million. Only tissues/cell types with at least ten reads from the *FAM230* family are shown.

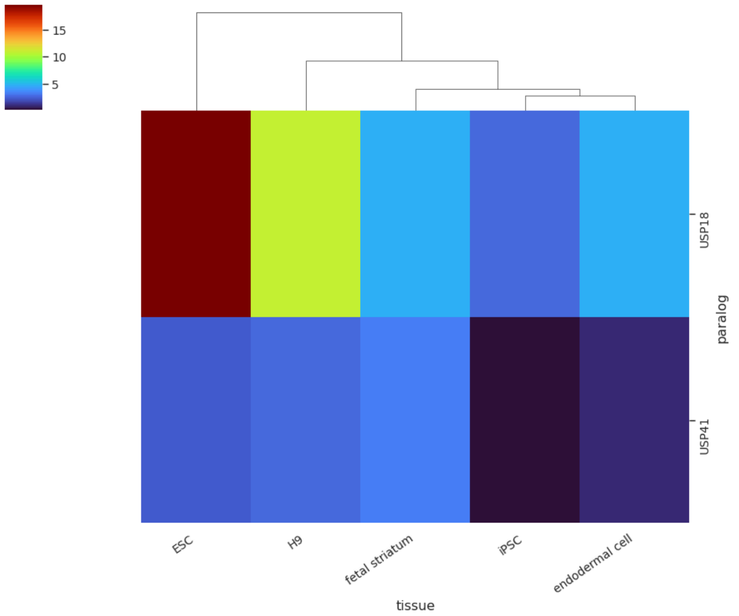


**Supplementary figure 50: Tissue specificity of Iso-Seq expression for *USP18* and *USP41*.** Estimated Iso-Seq expression of *USP18/41* paralogs as defined in T2T-CHM13, in reads per million. Only tissues/cell types with at least ten reads from *USP41* are shown.

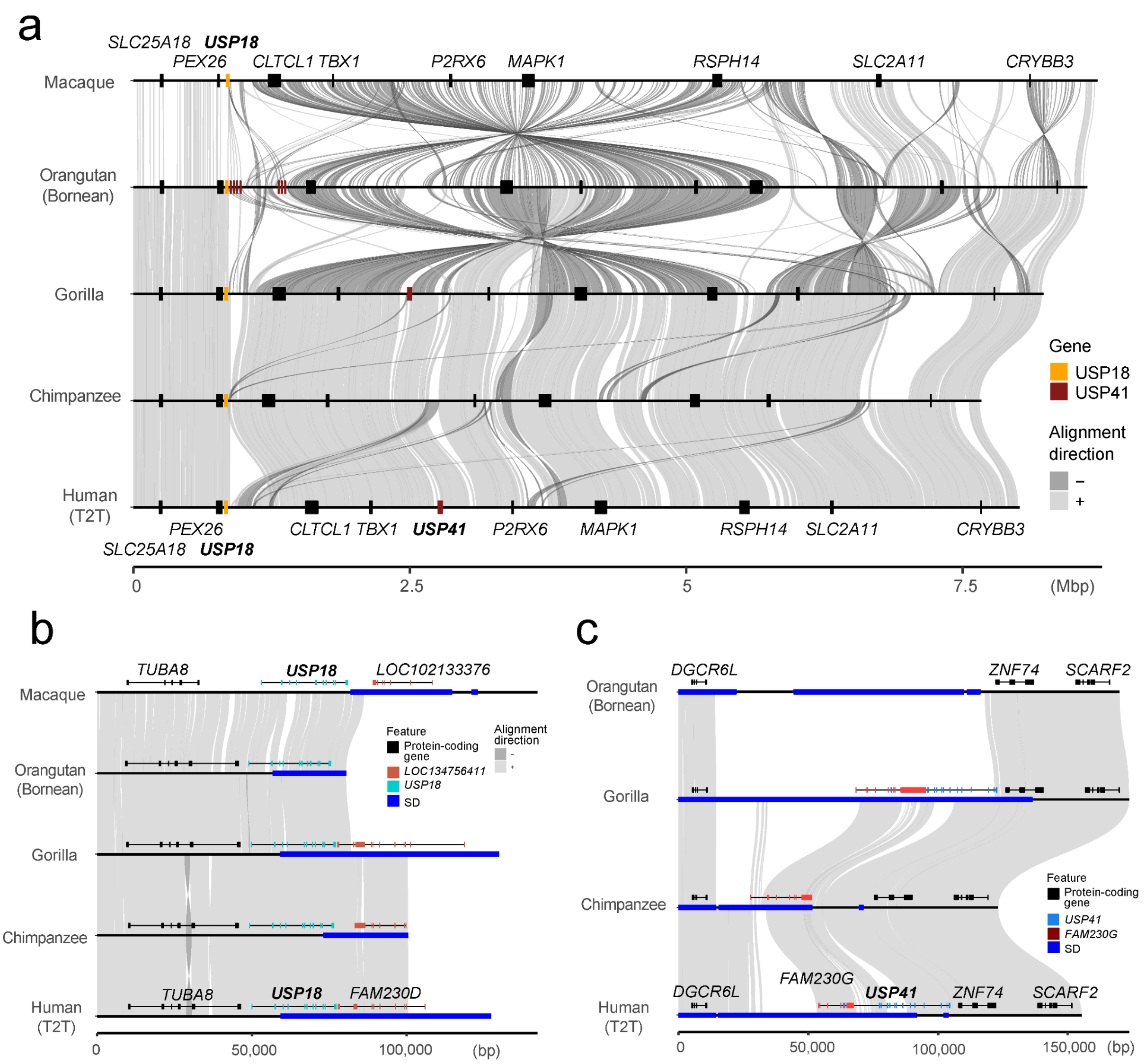


**Supplementary figure 51: Genomic position of *USP18* and *USP41*.**
**a**) Position of all USP gene copies with respect to the 18-26 Mbp sequence of T2T-CHM13 reference genome. Zoomed-in view near **b**) *USP18* conserved across the primate lineage and **c**) *USP41* present only in human and gorilla genomes.

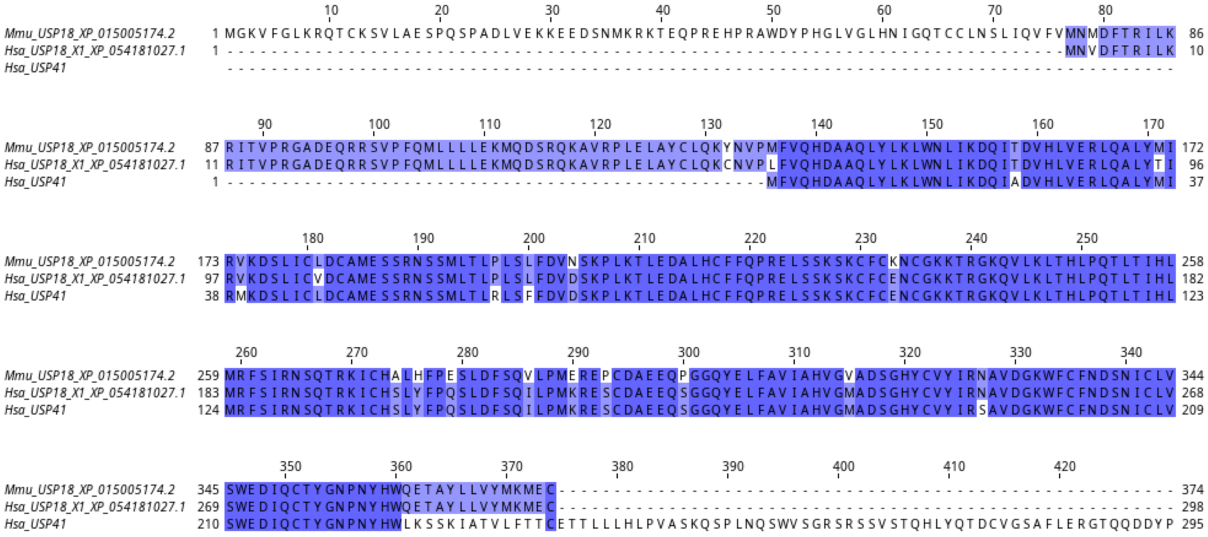

**Supplementary figure 52: MSA of USP41 protein sequence.**
Reference annotations of human (HSA) and rhesus macaque (MMU) USP18, compared to human USP41 open reading frame predicted from Iso-Seq.

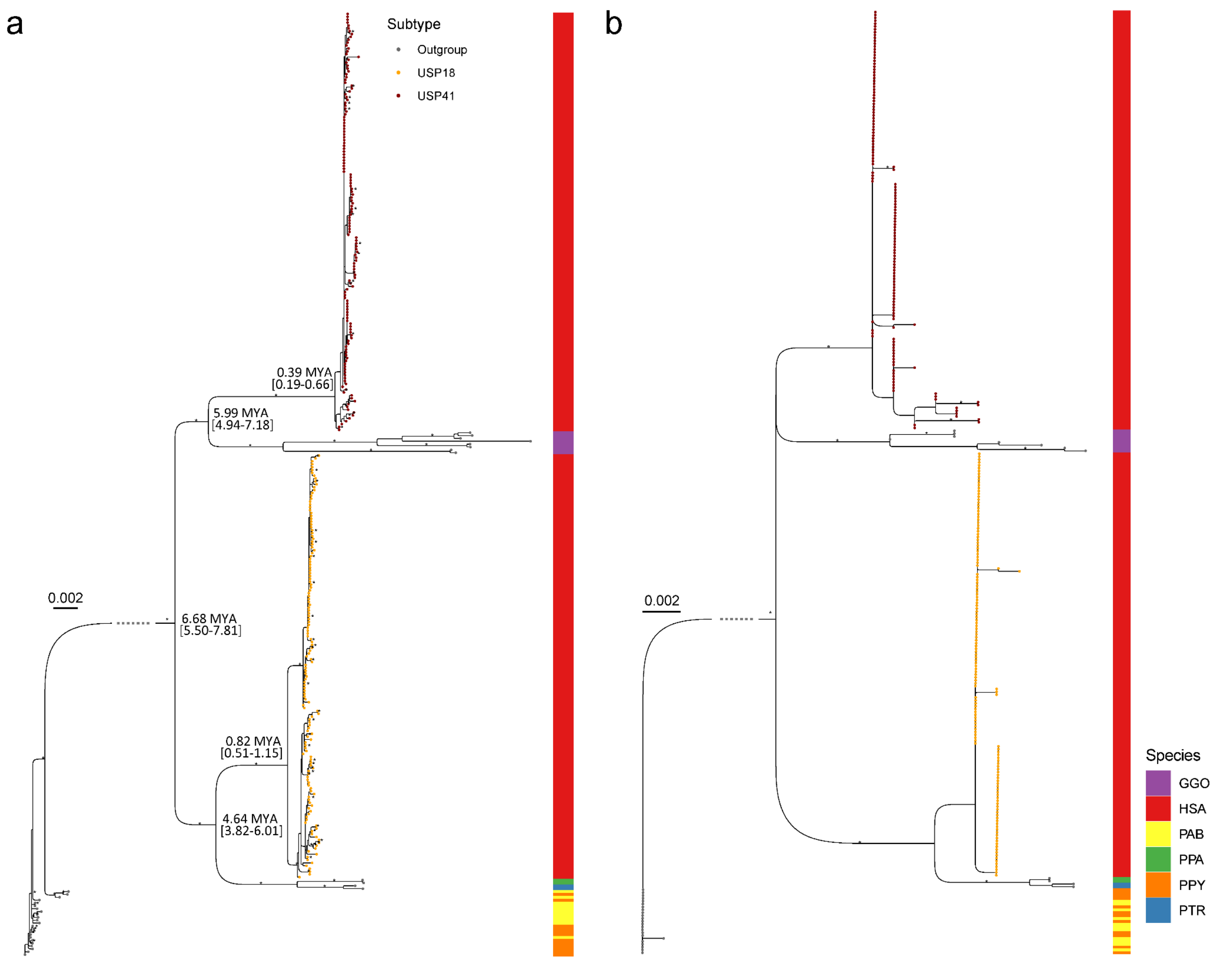


**Supplementary figure 53: Phylogeny of *USP18* and *USP41*.**
Maximum likelihood tree of concatenated **a**) intron and **b**) exon sequences of USP gene copies. The tree was outgrouped with orangutan orthologs. The copies corresponding to *USP18* and *USP41* are indicated at the tip nodes of the trees in orange and red, respectively, while the nonhuman orthologs are indicated in gray. The corresponding species is indicated on the right annotation track: chimpanzee (PTR), bonobo (PPA), human (HSA), gorilla (GGO), Sumatran orangutan (PAB) and Bornean orangutan (PPY). Notable divergence times among the copies are indicated as million years ago (MYA) with the 95% confidence interval. The bootstrap value of >95 is indicated as an asterisk.

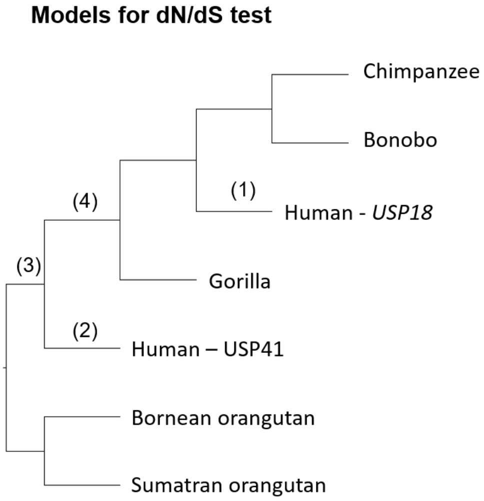


| Model | (1) | (2) | (3) | (4) |
|---|---|---|---|---|
| Dnds | 1.0 | 61.9 | 2.7 | 999.0 |
| h0lnL | -950.3 | -950.0 | -949.6 | -949.6 |
| h1lnL | -950.3 | -949.9 | -949.4 | -949.1 |
| LRT | 0.000 | 0.266 | 0.309 | 1.080 |
| p-value | 1.000 | 0.606 | 0.578 | 0.299 |
| Putative sites under selection (BEB posterior)* | NA | 27 P (0.646)<br>30 L (0.640)<br>157 N (0.638) | 56 R (0.602)<br>70 H (0.604)<br>71 R (0.591)<br>80 F (0.605)<br>119 T (0.590)<br>121 Q (0.619)<br>130 P (0.609)<br>134 F (0.613) | NA |

*site / amino acid / (probability)

**Supplementary figure 54: CODEML branch-site model test (dN/dS) of positive selection for the gene *USP41*.**
The modules (1-4) indicate the branch from which the dN/dS was estimated. On the right shows test statistics for individual models, including dN/dS (Dnds) ratio, log likelihood of null (neutral–fixing dN/dS to 1; h0lnL) and alternative (non neutral–allowing dN/dS ratio to vary; h1lnL) hypothesis, log likelihood test statistics (LRT) and p-value derived from the LRT. Putative sites under selection reported are summarized at the bottom row, with none of the sites showing Bayes Empirical Bayes (BEB) posterior probability >0.95.
